# Systematic evaluation of single-cell multimodal data integration for comprehensive human reference atlas

**DOI:** 10.1101/2025.03.06.637075

**Authors:** Mario Acera-Mateos, Xian Adiconis, Jessica-Kanglin Li, Domenica Marchese, Ginevra Caratù, Chung-Chau Hon, Prabha Tiwari, Miki Kojima, Beate Vieth, Michael A. Murphy, Sean K. Simmons, Thomas Lefevre, Irene Claes, Christopher L. O’Connor, Rajasree Menon, Edgar A. Otto, Yoshinari Ando, Katy Vandereyken, Matthias Kretzler, Markus Bitzer, Ernest Fraenkel, Thierry Voet, Wolfgang Enard, Piero Carninci, Holger Heyn, Joshua Z. Levin, Elisabetta Mereu

## Abstract

The integration of multimodal single-cell data enables comprehensive organ reference atlases, yet its impact remains largely unexplored, particularly in complex tissues. We generated a benchmarking dataset for the renal cortex by integrating 3’ and 5’ scRNA-seq with joint snRNA-seq and snATAC-seq, profiling 119,744 high-quality nuclei/cells from 19 donors. To align cell identities and enable consistent comparisons, we developed the interpretable machine learning tool scOMM (single-cell Omics Multimodal Mapping) and systematically assessed integration strategies. “Horizontal” integration of scRNA and snRNA-seq improved cell-type identification, while “vertical” integration of snRNA-seq and snATAC-seq had an additive effect, enhancing resolution in homogeneous populations and difficult-to-identify states. Global integration was especially effective in identifying adaptive states and rare cell types, including WFDC2-expressing Thick Ascending Limb and Norn cells, previously undetected in kidney atlases. Our work establishes a robust framework for multimodal reference atlas generation, advancing single-cell analysis and extending its applicability to diverse tissues.

## Introduction

Single-cell genomics is a fast-evolving field and provides tools to understand complex organs in incredible detail. Each single-cell method, whether for studying RNA or open chromatin (ATAC), offers a unique perspective into cell identity and function. While large-scale scRNA-seq datasets are the most prevalent, transcriptional profiles can be distorted during single-cell isolation, and cellular representation can be biased by incomplete tissue preparation^1,2^. Complementing scRNA-seq with protocols that provide full-length and nuclear RNA profiles is often adopted to minimize biases and improve data accuracy. Additionally, transcriptome profiling alone may not capture the full spectrum of cellular diversity while incorporating other modalities, like epigenetic measurements (e.g., open chromatin and DNA methylation) can enhance the completeness and resolution of an organ atlas. As such, researchers have increasingly employed multimodal, both paired and unpaired, single-cell measurements to generate reference atlases of human samples, as exemplified by large-scale efforts like the Human Cell Atlas (HCA)^3^. These approaches have significantly advanced the characterization of cellular heterogeneity and molecular diversity by enabling cross-validation of findings and elucidating functional relationships between different molecular layers^4,5,6,7^. However, the need for systematic benchmarking to evaluate the unique contributions of distinct modalities in resolving cell types and states has also emerged. While recent efforts in benchmarking single-cell approaches have primarily focused on comparing experimental methods within specific classes of single-cell assays, such as single-cell RNA sequencing (scRNA-seq)^8,9,10^ or single-cell ATAC sequencing (scATAC-seq)^11^, they have not addressed their integration. These studies have provided valuable insights into the efficiency and technical biases of each method in detecting molecules within cell types, offering a relevant but partial view of their unique ability to characterize complex tissues. Additionally, computational tools have been compared in their performance to correct batch effects and preserve biological variability in the integration of scRNA-seq datasets alone^12^ or in combination with scATAC-seq data^13^. However, these comparisons have not examined the advantages of joint integrations for defining cellular types based on multimodal data.

In the present study, we extend prior investigations by exploring the power of integrating multimodal single-cell omics data, offering a broader perspective on their combination in representing tissue complexity accurately and comprehensively. Using the kidney as an emblematic example of a complex organ, we conducted a multicenter comparative study, analyzing 33 samples, including 25 from donors with multiple samples (paired) and 8 from donors with only one sample (unpaired). This included single-cell 3’ and 5’ transcriptomic data, as well as multiomic single-nucleus RNA (snRNA-seq) and chromatin accessibility (snATAC-seq) data from the same cells. Additionally, we processed a subset of samples with Smart-seq2 and single-cell nucleosomal occupancy, DNA methylation and transcriptome sequencing (scNMT-seq) to further support the multimodal characterization. We systematically evaluated: i) how each modality contributes to the identification and characterization of specific cell types and ii) how the combination of multimodal data deepens our understanding of distinct cell types compared to any single modality. Our analysis includes three main components: 1) data harmonization, which standardizes and preprocesses single-cell data from different sources to ensure compatibility and minimize biases; 2) multimodal integration, which integrates and analyzes diverse data types; and 3) evaluation and benchmarking, which assesses the performance and efficiency of the integrated data in defining a wide range of cell types. Here, we introduce scOMM, a novel analytical framework designed to ensure a consistent projection of diverse data types onto a reference dataset while providing interpretable feature importance scores for cell type classification. Built on a supervised neural network strategy, scOMM learns cell identities and facilitates benchmarking across modalities, offering insights into their relative contributions to cell type identification and marker feature detection. We combine scOMM with unsupervised approaches, such as graph embedding followed by clustering, to explore the intrinsic structure of the data, offering supplementary insights into cell type diversity and relationships. To fully dissect distinct scenarios in cell atlas projects, we leveraged the established concept of anchors in single-cell multiomics integration^14^, which use shared elements to align and integrate different datasets. Using this framework, we compare different types of integrations: matching the same type of data from different sources (i.e., horizontal), combining different kinds of data from the same cells (i.e. vertical), and mixing different types of data from different cells, as well as incorporating paired cells when available (i.e. diagonal or mosaic).

Altogether, our work elucidates the synergistic value of integrating distinct single-cell data types, offering an integration framework that results in a more robust definition of kidney cell types and states. This framework can serve as a set of standards and guidelines for integrating multimodal data, highlighting the specific features of integration depending on the characteristics of the cell types being analyzed, and providing a roadmap for similar analyses in other complex tissues.

## Results

### Generation of a benchmarking dataset for single-cell multimodal characterization of the human renal cortex

Our study started with kidney tissue collection in the operating room from donors undergoing nephrectomy. Multiple aliquots from each normal tissue sample were frozen under different conditions selected to be appropriate for a variety of assays to profile them (see Methods). These aliquots were thawed and processed for 3’ scRNA-seq, 5’ scRNA-seq, and multiomics (snRNA-seq and snATAC-seq), which were central to our integrative framework (Fig. 1, Supp. Fig. 1). In addition, we also processed a small number of samples with Smart-seq2^15^ and scNMT-seq^16^, which are reported here separately because these assays were performed on a limited scale, generating insufficient data for our main analyses.

**Figure 1.**
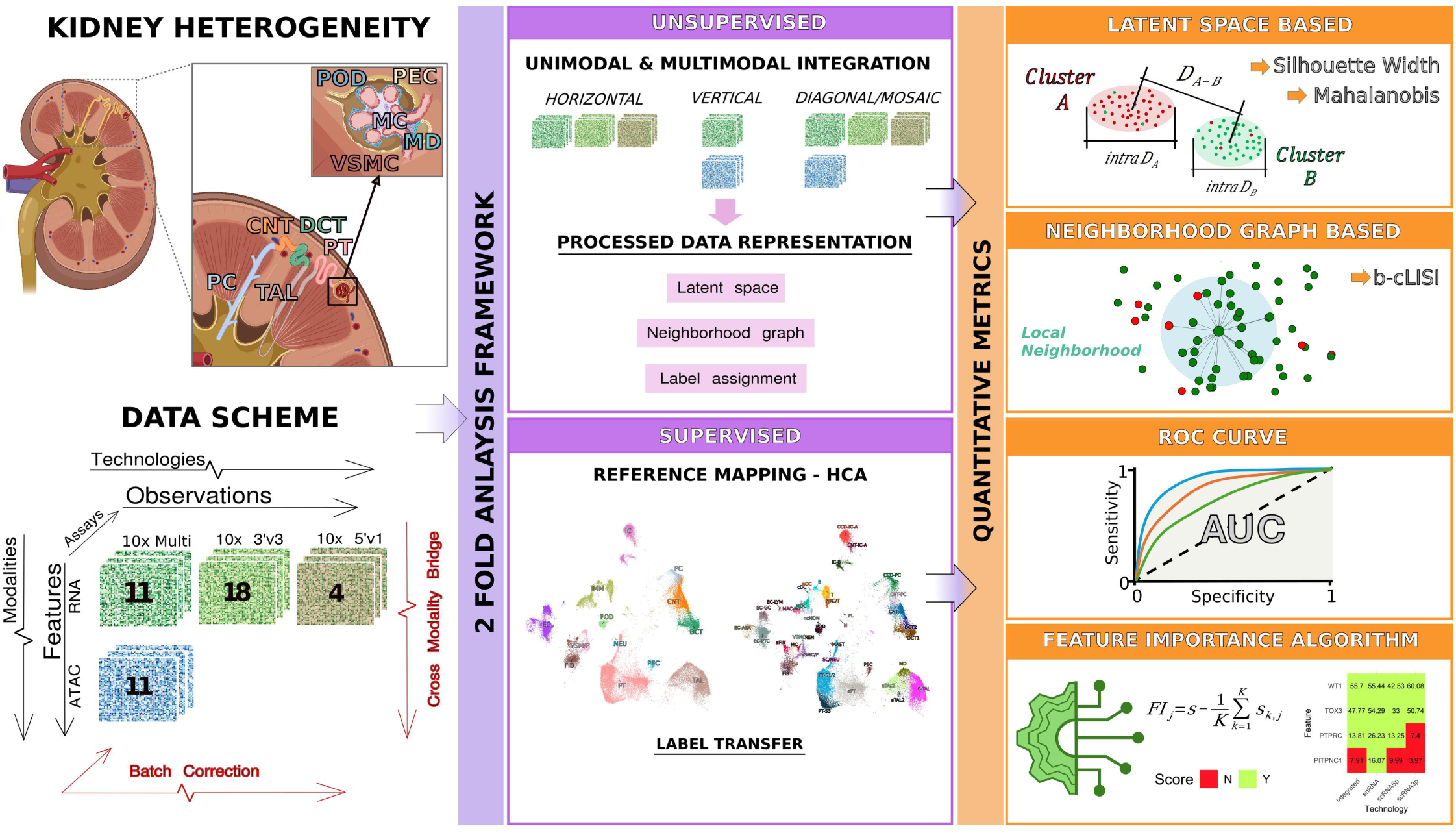
Experimental design and computational workflow. Overview of the study, integrating multimodal single-cell data from 33 kidney cortex samples from 19 matched and unmatched donors. Samples were analyzed using multiome snRNA-seq, multiome snATAC-seq, 3’ scRNA-seq, and 5’ scRNA-seq to create a comprehensive, multi-layered resource. Single-cell multiomics integration was conducted at different levels (with shared and unshared features) using unsupervised and supervised methods, such as graph embedding, clustering, and cross-modality bridging, to harmonize cell annotations and evaluate integration performance across cell types. Quantitative metrics, including latent space- and neighborhood graph-based, AUC, and feature importance, were employed to assess each modality’s contribution to cell type identification and the improved resolution achieved through multimodal integration.

This generated a multimodal benchmarking dataset for renal cortex (mBDRC) characterization, encompassing 119,744 high-quality cells/nuclei (see Methods) from 19 healthy donors (Fig. 2A-B, Supp. Fig. 2), representing diverse sex, age, BMI, and other clinical renal-associated characteristics (Table 1, Table 2). The mBDRC was anchored on previously established human kidney references^5,17^ through which we have obtained two main layers of annotations, 12 broad cell types (referred to as L1 annotation) and 39 cell types/states (referred to as L2 annotation), encompassing epithelial, endothelial, immune and other stromal subtypes (Fig. 2C, Supp. Fig. 3A-B, Table 3). This comprehensive dataset enabled us to perform a comparison and integration of multiple single-cell genomics data types, assessing technical biases, estimating statistical power, and exploring complementarity and redundancy in cell type/state identification, described in the following sections. We first harmonized and annotated cells from each technology individually (see Methods). Subsequently, we integrated the distinct protocols and modalities using the different integration strategies: horizontal, vertical, and diagonal/mosaic. For the mBDRC mosaic integration (Fig. 2B), we utilized MultiVI^18^ to generate a lower-dimensional latent space and correct for batch effects. We conducted a series of iterative benchmarking investigations throughout the analysis pipeline (see Methods), assessing the integration of individual technology samples and across sn/scRNA technologies, and ultimately the integration of snRNA with snATAC data in both paired and unpaired settings. These steps are discussed in greater detail in subsequent sections. This stepwise evaluation provided immediate feedback on the performance and consistency of the integrated datasets at each stage, ensuring that MultiVI was a well-validated choice. Additionally, to harmonize cell identities between technologies, we developed scOMM (see Methods), a machine learning tool specifically designed for supervised cell-type annotation and benchmarking of multimodal single-cell data. ScOMM mapped each dataset onto the external references by leveraging their curated cell annotations. Unlike existing tools such as Harmony^19^ and Seurat^20^, which focus on general reference mapping or integration, scOMM also addresses the unmet need for systematic benchmarking by facilitating direct comparison of distinct data types, offering flexible parameter tuning, and evaluating feature importance for cell-type predictability.

**Figure 2.**
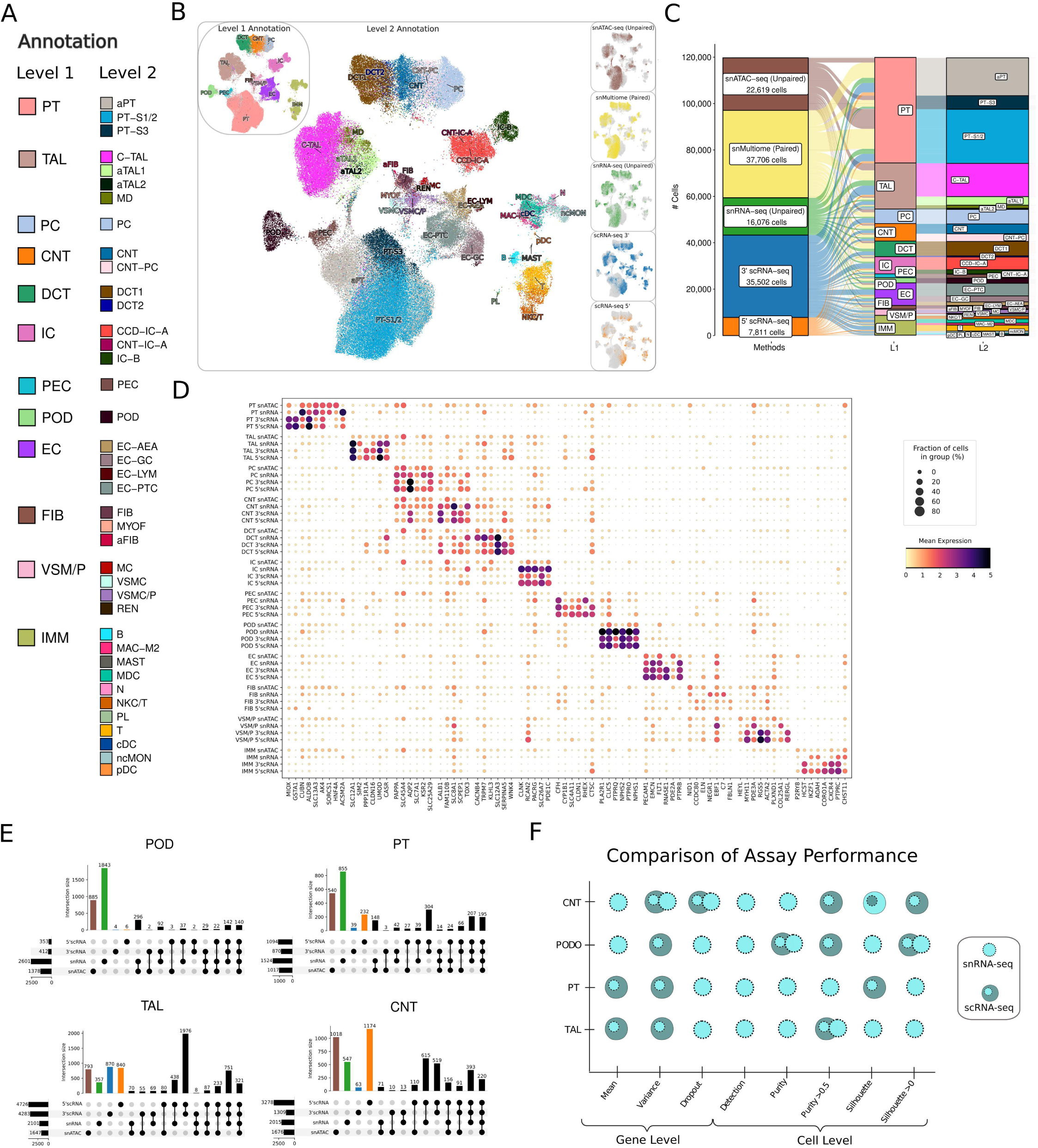
Overview of the multimodal Benchmarking Dataset for Renal Cortex (mBDRC). A) Color legend of the renal cortex cell types at the two levels of annotations. Level 1 (L1) represents 14 broad cell types, while Level 2 (L2) provides detailed annotations of 39 specific cell subtypes or states within these categories. Broad cell types and their subtypes are as follows: PT (Proximal Tubule) includes aPT, PT-S1/S2, and PT-S3; TAL (Thick Ascending Limb) includes cTAL, aTAL1, aTAL2, and MD (Macula Densa); PC (Principal Cells) includes PC; CNT (Connecting Tubule Cells) includes CNT and CNT-PC; DCT (Distal Convoluted Tubule) includes DCT1 and DCT2; IC (Intercalated Cells) includes CCD-IC-A, CNT-IC-A, and IC-B; PEC (Parietal Epithelial Cells) includes PEC; POD (Podocytes); EC (Endothelial Cells) includes EC-AEA, EC-GC, EC-LYM, and EC-PTC; FIB (Fibroblasts) includes MYOF, and aFIB; VSM/P (Vascular Smooth Muscle/Pericytes) includes MC, VSMC, VSM/P, and REN (Renin-producing cells); and IMM (Immune Cells) includes B, MAC-M2, MAST, MDC, NK/T, PL, T, cDC, ncMON, and pDC. B) UMAP plot of 119,744 nuclei/cells across different single-cell data modalities, including multiome snRNA, multiome snATAC, 3’ and 5’ scRNA. Colors indicate the refined cell-type annotations (L2), with broad cell-type annotations (L1) shown in the top-left squares.C) Alluvial plots displaying the distribution of cells analyzed across different protocols, with colors representing L1/L2 cell-type annotations. D) Dot plot showing consensus cell-type markers for the main renal cortex populations as detected by the different single-cell data types. E) Upset plot illustrating the overlap of detected markers identified by each technology for the primary epithelial populations, including POD, PT, DCT, and TAL. F) Summary Statistics for CNT, PT, POD and TAL, illustrating the best performing assay (snRNA-seq represented as a dark teal nucleus and scRNA-seq represented as a light teal cell) in terms of gene metrics: mean (the minimal median standard error of mean expression), variance (the minimal median variance of gene expression) and dropout (the minimal median dropout of gene expression); in terms of cell metrics: detection (the maximum median proportion of genes with nonzero expression per cell), purity (maximum median purity of k neighborhoods in a cell type), purity > 0.5 (maximal proportion of cells with 50% or more pure neighborhoods), silhouette (maximum median silhouette per cell type) and silhouette > 0 (maximal proportion of cells with silhouette index above 0). Both symbols are shown when they are essentially equivalent. See Supp. Fig. 6 for details underlying this summary.

Analyzing differentially expressed genes across the dataset revealed a consensus set of markers for each cell type (Fig. 2D), which demonstrated consistent co-expression patterns across platforms and modalities. Specifically, homogeneous populations, such as podocytes (POD), showed consistent detection and co-expression of canonical markers like *NPHS1* and *NPHS2* across all platforms, highlighting the robustness of these markers in defining well-distinguishable cell types. By contrast, populations such as distal convoluted tubules (DCT), connecting tubules (CNT), and principal cells (PC) displayed overlapping or gradient-like expression patterns, reflecting a spectrum of cellular states. This heterogeneity adds complexity to capturing these markers consistently across different technologies and emphasizes the value of integrating them. Indeed, comparing cell type-specific markers after downsampling cells and unique molecular identifiers (see Methods), showed that different technologies captured different markers more effectively (Fig. 2E, Supp. Fig. 3C, Table 4). For instance, *PARD3* and *COL4A5*, which are crucial for maintaining POD structure and function^21,22^, were identified as significant markers exclusively in the snRNA-seq data. Likewise, *SLC25A4*, a marker vital for the metabolic activity of CNT cells, was uniquely detected in the 5’ scRNA-seq dataset. Additionally, we performed Smart-seq2 and scNMT-seq analyses on four and two samples, respectively, yielding 597 and 1245 high-quality single-cell transcriptomes (Supp. Fig. 4A). Despite the limited sample size, these datasets align well with and reinforce the computational and manually curated markers identified using other protocols (Supp. Fig. 4B-C). For scNMT-seq, a small set of PT and DCT-CNT cells identified from transcriptomic data (Supp. Fig. 4A) was selected to match the DNA methylomes of these same cells (Supp. Fig. 5A). Since GpC methylation in scNMT-seq marks open chromatin, we further investigated the list of genes with open regions based on snATAC-seq data (see Methods; Table 3), which confirmed increased DNA accessibility at transcription start sites of those genes (Supp. Fig. 5B). Next, we investigated GpC methylation in promoter regions in genes detected as transcribed or non-transcribed on a pseudo-bulk level per cell type and provided evidence (see Methods) that non-transcribed genes were less accessible than transcribed genes (Pr(>*χ*^2^) < 0.01, Supp. Fig. 5C). Endogenous CpG methylation showed an overall reduction in promoter methylation which was more pronounced in transcribed than non-transcribed genes (Pr(>*χ*^2^) < 0.01, Supp. Fig. 5D-E).

### Empirical power analysis quantifies cell-type specific advantages of scRNA-seq and snRNA-seq

Before proceeding to the integration of different data types, we performed an empirical power analysis to quantify the advantages of scRNA-seq versus snRNA-seq, two widely used techniques in generating human organ atlases. Our mBDRC dataset was particularly well-suited for this analysis due to the unique advantage of profiling several donors using both protocols, enabling quantitative comparisons and specific recommendations for these assays. Using the scOMM broad cell type (L1) annotations and additional filtering steps to achieve balanced datasets for the different comparisons (see Methods), we first analyzed how much variance in expression levels can be explained by cell type (n=14), donor (n=5) and assay (3’ scRNA-seq and snRNA-seq). While assay (median 24%) and cell type (13%) explained most of the variance, also the interaction of assay and cell type explained a considerable proportion (6%; Supp. Fig. 3D). Hence, the different assays measured considerably different expression levels in different cells, exceeding differences among donors (3%). Nevertheless, gene expression differences among cell types were large and consistent enough to lead to similar distances among them, as reflected by the congruent cell type trees of scRNA-seq and snRNA-seq (Supp. Fig. 3E).

To compare the two assays on the gene level, we estimated mean, variance and detection rate of expression levels weighted by the number of cells. On the cell level, we estimated the proportion of genes per cell with non-zero expression levels weighted by number of reads and cells as well as the homogeneity of the cell type cluster based on its purity and silhouette (see Methods). We calculated values for all 14 cell types (Supp. Fig. 6) but restricted the visualization to the four main cell types for simplicity (Supp. Fig. 3F-G). In addition, we assessed reproducibility of metrics across our sampling cohort by calculating the Kolmogorov-Smirnov distance between assays within the same donor and between donors per assay (Supp. Fig. 6). We opted to rank the two assays based on eight summary statistics for the four main cell types (Fig. 2G). Intriguingly, in most cases snRNA-seq outcompeted scRNA-seq in terms of summary statistics as well as reproducibility. This is quantitative evidence for kidney tissue and probably also for tissues with similar cellular fragility that a) more information is gained when using snRNA-seq compared to scRNA-seq at both gene and cell levels; and b) that scRNA-seq and snRNA-seq modalities yields complementary information. Hence, their integration can improve cell-type characterization and provide a more comprehensive understanding of kidney complexity.

### Horizontal integration of sn/scRNA-seq data highlights differences in cell type identification accuracy across assays

To achieve a comprehensive transcriptional characterization of the cell types in the human kidney, we performed horizontal integration of sn/scRNA-seq datasets (Fig. 3A) using scVI^23^, which demonstrated the highest benchmarking scores (see Methods) for integration both within and across protocols (Supp. Fig. 7A, Supp. Fig. 8A). This integration approach effectively reduced technical noise (as indicated by the highest batch correction scores) and enhanced the detection of true biological signals (reflected in the highest biological conservation scores). Then, clustering produced cell-type annotations that were consistent with scOMM across datasets at both broad (L1) and fine (L2) annotation levels (Supp. Fig. 7B-C, Supp. Fig. 8B-C). To evaluate the relative contribution of each protocol to the transcriptional definition of each cell type, we first harmonized cell type annotations (see Methods) through the inspection of cluster-specific markers (Supp. Fig. 7B). The integrated data then served as a biological reference or “ground truth” for each cell type/subtype in subsequent comparison. We, thus, compared the annotations to the reference-based (scOMM) cell mapping obtained independently and from each sn/scRNA-seq dataset (Supp. Fig. 7D). By evaluating area under the curve (AUC) scores for each cell type/subtype (see Methods), stratified by protocol (Fig. 3B), we quantified how each dataset independently contributed to cell type definition, leveraging the strengths of integrated data while minimizing biases. High AUC scores for a specific cell type suggest strong agreement between the unsupervised clustering-based annotations and the supervised reference-based annotations (i.e., obtained by scOMM), indicating effective capture of the transcriptional cell type signature. Conversely, low AUC scores indicate discrepancies, reflecting a lower effectiveness for cell type identification. We focused on high specificity values (>0.9; see Methods) to prioritize the accurate identification with minimal false positive rates. We found that PT subtypes (e.g., PT-S1, aPT, PT-S3) were more accurately characterized by snRNA-seq, as indicated by higher AUC scores. Conversely, TAL groups (C-TAL, aTAL) were better identified with 3’ scRNA-seq data, while transcriptionally continuous and low-abundant populations (e.g., MD, DCT1, CNT) were better captured by 5’ scRNA-seq data (Fig. 3B).

**Figure 3.**
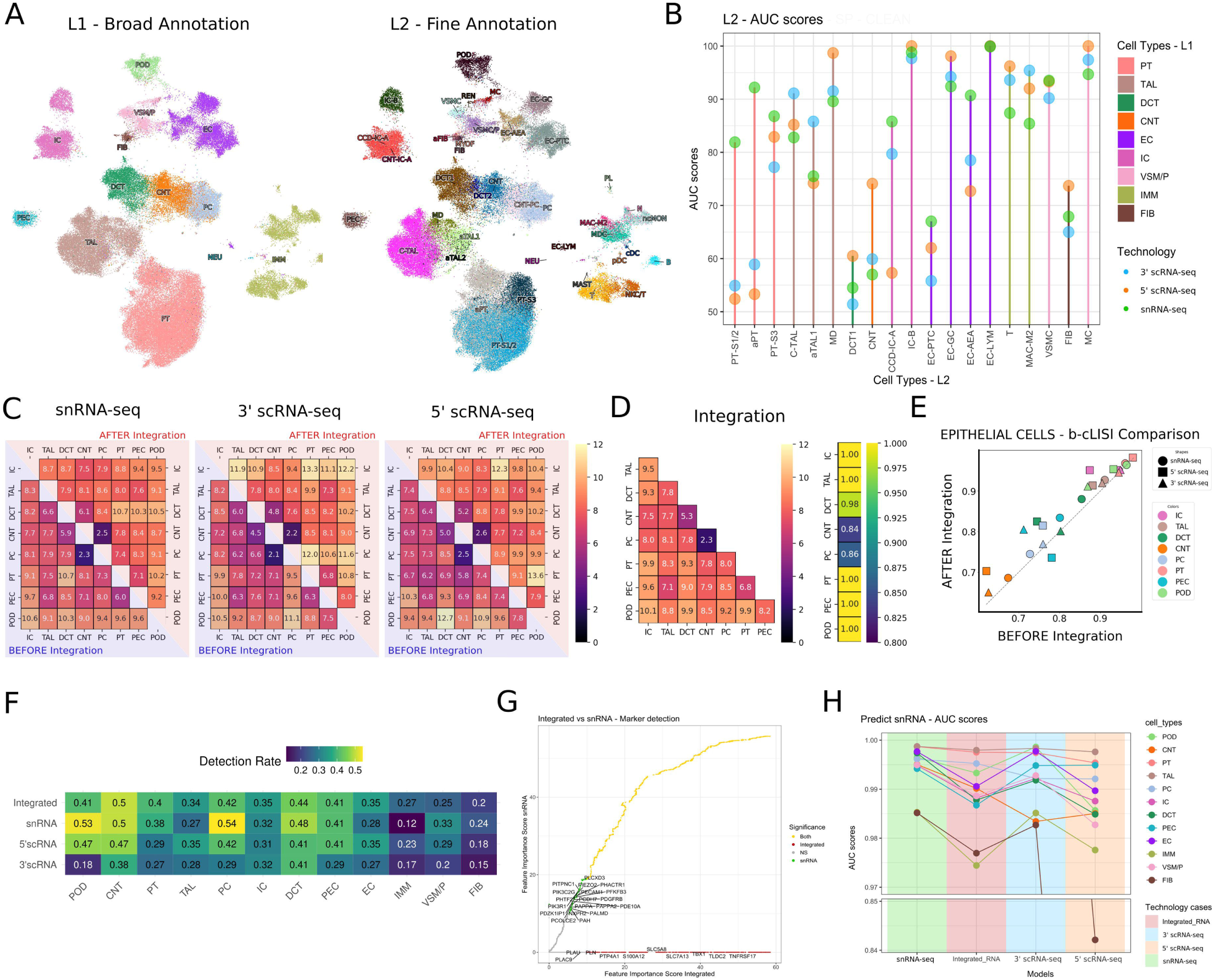
Horizontal integration of sn/scRNA-seq data. A) Horizontal integration of 97,125 nuclei/cells from 33 renal cortex samples. Colors represent the inferred annotations from HCA reference mapping, using broad L1 (left) and fine L2 cell types/subtypes (right). B) AUC scores from scOMM L2 annotations compared to clusters from the horizontal integration in A. Colored dots indicate modalities, while bars represent L2 cell groups, with color bars stratified by L1 cell groups. C) Pairwise Mahalanobis distances (see Methods) between epithelial populations within each technology before and after integration, where higher values indicate greater separation between populations. D) Mahalanobis distances between epithelial-derived cell-types in the embedding space (left). Summary of the cell type-specific Mahalanobis distances as overlap ratios derived from a Chi-squared distribution (right), with values closer to 1.0 indicating greater separability (lower overlap) between cell types, and values closer to 0 reflecting higher overlap (less separability). E) Binary cLISI (b- cLISI; see Methods) scores for L1 epithelial groups calculated before and after integration. F) Marker detection rate comparison based on the feature importance of L1 cell types across data models. G) Feature importance score comparison between snRNA-seq and the integrated dataset for markers that showed significant FI in the snRNA model but did not exhibit significant FI in the 3’ and 5’ scRNA models. H) AUC scores for the predictability of each data model in predicting the snRNA-seq data type for L1 cell-type annotations. The background is divided into shaded regions that correspond to different sn/scRNA technologies. The dots, connected by lines, represent the AUC scores for individual cell types, with distinct colors assigned to each cell type.

To further assess the resolution of the cell-type representations generated by the graph embedding and latent space approximation, we used the Mahalanobis distance (see Methods), which accounts for cell-type distribution variance and correlation structure, providing a robust metric of cluster separation in high-dimensional data. Calculating Mahalanobis distances independently for each dataset (before and after sample integration) revealed that individual protocols showed similar cell-type relationship, with consistently higher distance scores observed across all populations following the integration of samples, indicating improved performance in separating distinct cell types (Fig. 3C). While integration across protocols did not necessarily outperform individual datasets in all cases, it mitigated protocol-specific deficits, resulting in a more balanced and robust representation of cell populations, such as in IC, PT, and DCT, where distances were consistently greater than the minimum observed in individual protocols or near their average (Fig. 3D). Complementing this, the binary cell-type Local Inverse Simpson’s Index (b-cLISI; see Methods), which measures biological conservation and local diversity after integration, provided higher scores post-integration (Fig. 3E, values above the diagonal). This improvement was particularly notable in continuous and transitional populations, such as CNT, DCT and PC. Here, the DCT population, exhibited a clear improvement in both Mahalanobis (Fig. 3D, summary score 0.98) and b-cLISI (Fig. 3E, with values >0.8 after integration observed in snRNA and 5’ scRNA), reflecting its accurate separation from CNT after integration despite being a challenging case due to their biological proximity. Together, these results show how integration enhances both global separation (as assessed by Mahalanobis distances) and local diversity (as captured by b-cLISI), resulting in a comprehensive and nuanced view of cell population relationships.

### Integration of sn/scRNA-seq modalities boosts marker discovery and cell type predictability

Identifying marker genes that exhibit variable expression across populations is essential for classifying cell types in sn/scRNA-seq. However, this process can be biased by technical noise, dropouts, and batch effects, which may mask true biological signals and hinder accurate marker identification. By combining data from different sn/scRNA-seq protocols, we investigated whether this could improve the detection of cell type-specific markers. Also, we examined if this improvement varied depending on the characteristics of each cell type. We trained a scOMM model for each sn/scRNA-seq dataset, downsampling to 6,000 nuclei/cells per dataset to reduce biases associated with varying cell numbers. The models were trained using all potential marker genes (see Methods) identified independently within each dataset among their differentially expressed genes. To determine which genes were most important for distinguishing cell types, we applied a feature importance (FI) approach (see Methods) that perturbs the data and evaluates the impact on the overall cell type prediction. This method simulated the elimination of a gene’s contribution to the model by replacing the expression values of each tested gene with zeros. Genes with higher FI values were those that increased the model’s accuracy. We set a threshold of 10 for FI to indicate significance, meaning that when perturbing a gene resulted in a reduction of at least a relative 10 points in the model’s accuracy, the gene was considered critical for distinguishing cell types. Next, we also trained a model (see Methods) to assess the impact of combining data from different protocols on marker detection. The detected markers were compared to the cell-type markers listed in the HCA kidney reference, which integrates analyses from both single-cell and single-nucleus RNA-seq data, thereby minimizing assay-specific bias. We observed substantial differences in marker detection rates across protocols and cell types (Fig. 3F), with snRNA-seq consistently demonstrating the highest detection rates for several cell types, particularly POD, PC, and DCT, at around 0.5. This may be due to snRNA-seq avoiding issues associated with cell dissociation. The integrated dataset demonstrated strong performance across most cell types, again balancing the performance of individual dataset (Supp. Fig. 9A). Notably, the 5’ scRNA-seq dataset performed comparably well in several cell types, particularly in POD and TAL, where it showed the highest detection rates. On the other hand, the 3’ scRNA-seq dataset showed lower rates overall, with a very low score surprisingly observed in POD. The marker detection performance also highlights the unique, common, and partially shared markers across the different protocols (Supp. Fig. 9B-C), where snRNA-seq and the integrated dataset resulted in a higher proportion of unique markers, particularly in POD (e.g., *PCOLCE2* significantly detected only in snRNA-seq), epithelial cells (e.g., *SLC7A13* and *SLC5A8* specific to PT-S3, *PTP4A1* as a marker of CNT-PC significantly detected exclusively in the integrated data), and endothelial cells (e.g., *TBX1* in the integrated dataset and *PECAM1* in snRNA-seq) (Table 5). Importantly, assay-specific markers that were also detected as significant in the integrated model (Fig. 3G, Supp. Fig. 9D-E, referred to as “both” and highlighted by the yellow line in the figures) exhibited higher FI scores compared to those from their original protocol, indicating their relevance in distinguishing cell types. Furthermore, we compared the AUC scores of the integrated model across all cell types when predicting cells from each standalone assay-specific dataset (Fig. 3H, Supp. Fig. 9F-G), confirming that integrating data from multiple protocols not only improved marker detection but also provided a more robust framework for cell-type classification.

### Vertical integration of snRNA- and snATAC-seq data enhances cell subtypes identification beyond single-modality approaches

To perform vertical integration, we analyzed 11 multiome samples, generating simultaneous snRNA-seq and snATAC-seq data from 37,717 high-quality nuclei (Supp. Fig. 2; see Methods). To harness the full potential of both modalities, we employed two approaches specifically designed for single-cell multiomics analysis: multimodal spectral (multi-spectral) (Fig. 4A) as implemented in SnapATAC2^24^ and the Weighted Nearest Neighbor (WNN) from Seurat^7^. While both methods enable the generation of a joint embedding of RNA and ATAC data, WNN assigns cell-specific modality weights, prioritizing the most informative data type for each cell. This resulted in predominantly RNA-weighted data across all cell types (Supp. Fig. 10A), diminishing the contribution of snATAC-seq data to overall cell profiling. In light of this limitation, where RNA data was predominantly prioritized over ATAC data, we conducted additional analyses to evaluate the specific contribution of snATAC-seq data in renal cortex samples. We harmonized snATAC-seq samples (see Methods) similarly to RNA horizontal integration and assessed the performance of four widely used computational pipelines: SnapATAC2^24^ (referred to as Spectral MNN), Signac^25^ (referred to as LSI Harmony), PeakVI^26^, and PoissonVI^27^ (Supp. Fig. 11A). In agreement with recent benchmarking studies^28^, Spectral MNN best captured the complex structure of the kidney cortex, even at finer L2 resolution (Supp. Fig. 11B). Unsupervised clustering further supported this, showing a high level of agreement between snATAC cluster annotations and their RNA counterparts, with minimal mismatches in rare populations such as macula densa (MD) cells and transitional states like CNT-PC and PT-S1/2 (Supp. Fig. 10B-C). Based on these findings, we first optimized WNN by combining the best snRNA-seq and snATAC-seq embeddings derived from their respective horizontal integrations (Supp. Fig. 10D; See Methods). We then compared this WNN result with multi-spectral for vertical integration. Despite this optimization, a comparison of silhouette scores revealed that multi-spectral consistently achieved higher scores across several subtypes, resulting in greater average deviance, which indicates better-separated cell groups (Fig. 4B, Supp. Fig. 10E). This trend was particularly evident in populations such as POD and PEC, which are typically well-resolved through clustering. We hypothesize that WNN underrepresented complementary ATAC data, limiting its ability to capture subtle cellular differences, whereas multi-spectral enhanced sensitivity to chromatin accessibility, resulting in stronger cluster separability. Indeed, a comparison of average silhouette width scores from individual assays (RNA and ATAC) with those from vertically integrated data (i.e., joint RNA and ATAC) revealed a complementary effect, approximating an additive trend with the multi-spectral method (Fig. 4C, top table). This pattern was especially pronounced in homogeneous populations and extended to more complex subtypes, such as C-TAL and IC-B, underscoring the value of integrating both modalities. Rather than simply balancing the contributions of RNA and ATAC, the multi-spectral approach appears to optimally combine them, enhancing cell type identification by preserving the complementary strengths of each technology. At the local level, b-cLISI scores further demonstrated multi-spectral superiority in preserving neighborhood structure (Fig. 4C, bottom table). Higher b-cLISI values in challenging-to-identify cell states, such as aPT, CNT-PC, and DCT1, confirmed the benefits of multimodal integration. Altogether, these findings highlight the importance of leveraging both transcriptomic and chromatin accessibility data in parallel to achieve a more nuanced understanding of cellular heterogeneity, particularly in complex tissues.

**Figure 4.**
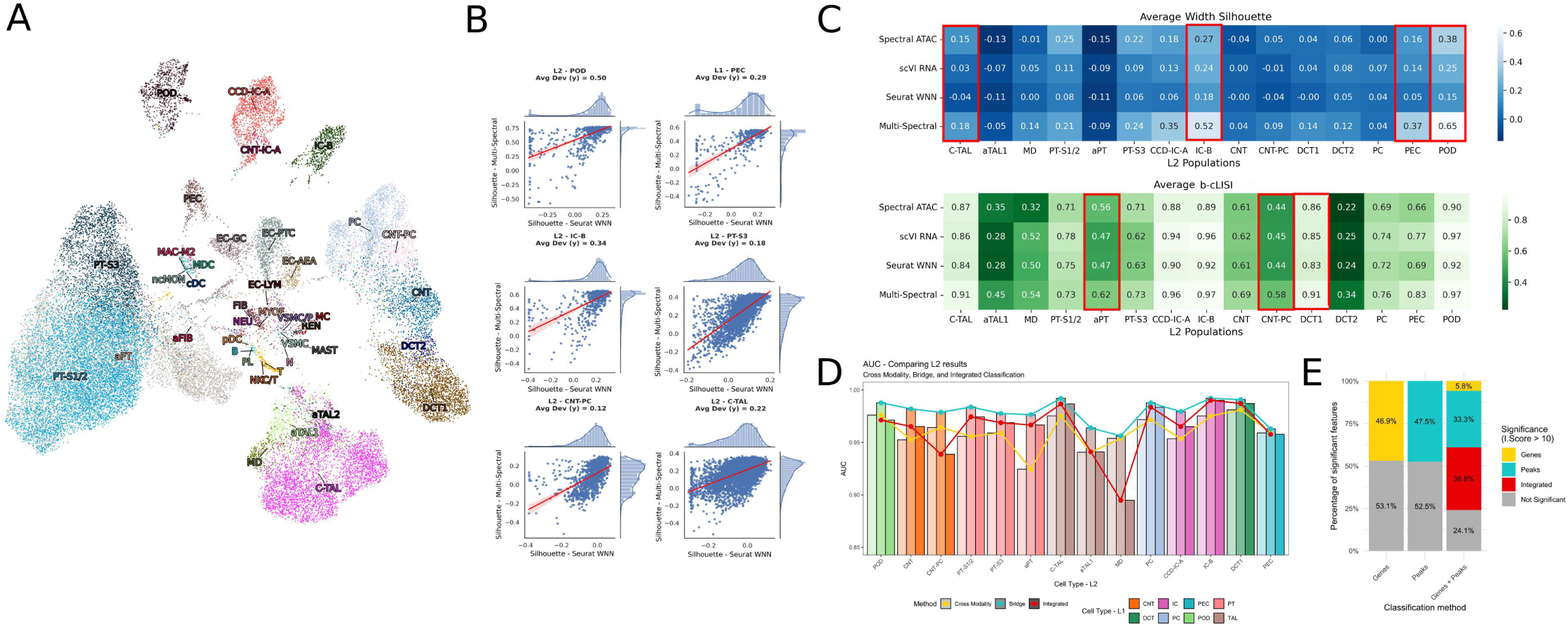
Vertical integration of snRNA/snATAC-seq data. A) Multiomics integration of 37,717 paired nuclei obtained with the multimodal spectral (multi-spectral) approach. B) Scatter plots compare silhouette scores from optimized WNN (X-axis) and multi-spectral (Y-axis) integration methods for various kidney cell types and subtypes. Each blue dot represents a single cell, while the red line indicates the trend, with the pink shading showing the confidence interval. Marginal histograms illustrate the distribution of silhouette scores along each axis. The “Avg Dev (y)” value represents the average deviation of the Y-axis (multi-spectral scores) for each cell type or subtype. C) The heatmaps compare four data integration methods across kidney L2 cell populations. Average Width Silhouette (AWS) are shown at the top, with darker blue indicating worse performance. B-cLISI scores are shown at the bottom, where darker green represents worse clustering consistency. Red boxes highlight key cell populations (C-TAL, aPT, CNT-PC, IC-B, POD) that exhibit improved values with vertical integration using the multi-spectral approach. D) Bar charts compare AUC scores across L2 cell types using three classification methods for predicting cell types in the paired snATAC-seq data: Cross Modality (yellow), Bridge (cyan), and Integrated (red). E) Stacked bar charts show the percentage of significant features (Genes, Peaks, and Genes + Peaks) across the three classification models with significance defined by a score threshold of 10. The colors represent the percentage of significant features for Genes (yellow), Peaks (cyan), and Genes + Peaks (red), while gray indicates the percentage of features that are not significant (Score ≤ 10).

### Evaluation of the contribution of chromatin accessibility to cell identity prediction and profiling consistency with snRNA data

To further examine the similarity between RNA and ATAC data and assess how well snATAC-seq captures regulatory features reflective of transcriptional identities, we trained two scOMM models (see Methods). These models transferred cell identities from snRNA to snATAC using either gene activity (i.e., peaks-associated genes are used as features for the model, also referred to as Cross-Modality, CM) as a proxy for gene expression or through bridge integration (referred to as Bridge), which directly incorporates the accessible peaks as model features. Additionally, we trained a third model (referred to as Integrated) that uses both types of features as input data to determine if combining the distinct features derived from each modality improves cell-type detection. In all models, cell type annotations from snRNA data served as the ground truth for performance evaluation. Model accuracy, which was measured by AUC, was highest for the Bridge model across most cell types, while the Integrated model displayed comparable performance for specific subtypes, such as PT, C-TAL, IC-B, DCT1, and PEC (Fig. 4D). Notably, both the Bridge and Integrated models improved sensitivity and classification rates compared to the CM model, whereas specificity remained similar across all three approaches (Supp. Fig. 11C-D). The drop in performance for the CM model may be attributed to discrepancies between gene activity (reflecting chromatin accessibility) and actual gene expression (Supp. Fig. 11E), as well as the lack of non-coding region information in snRNA-seq data. We also examined the contribution of the two different modalities to cell identity prediction by comparing the significance of peaks and/or genes across models. In the unimodal models (CM and Bridge), roughly half of the features showed high importance for cell-type prediction (Fig. 4E). In the Integrated model, which used both genes and peaks, peaks appeared to play a more dominant role than genes, likely due to the greater number of peaks relative to genes in the feature space. Interestingly, some features that were not significant in the unimodal models became relevant, exhibiting high feature importance in the integrated model (represented in red in the Genes + Peaks bar, Fig. 4E). However, despite the prominence of peaks and newly significant features in this model, overall cell type prediction accuracy did not outperform that of the Bridge model (Fig. 4D), suggesting that the added complexity, including the higher dimensionality introduced by peaks, may introduce redundancy or noise without providing substantial improvements in predictive power. Therefore, while integrating both data types uncovers new relationships, it does not necessarily enhance cell type predictability.

### Global data integration improves detection and characterization of rare cell types by refining signals across modalities

The combination of distinct single-cell data types can be approached through diagonal and mosaic integrations. In diagonal integration, distinct features (e.g., RNA and ATAC) are combined from unpaired cells, while in mosaic integration, these features are measured in paired or unpaired cells, providing a more comprehensive view. It is important to investigate: i) whether paired data significantly improves cell-type identification and characterization within global mosaic integration, and ii) whether integrating all data types in an unpaired fashion outperforms simpler combinations. To integrate all the data, we utilized MultiVI^18^ and GLUE^29^ (Supp. Fig. 12A-B), two models designed for single-cell multiomics integration. MultiVI is suited for mosaic integration, where paired data are leveraged for optimized integration, while GLUE handles diagonal integration using unpaired data. Interestingly, no significant differences were observed between these two approaches at both the local level, referring to the b-cLISI score (Fig. 5A), and the global level, corresponding to the silhouette score (Supp. Fig. 12C), with cell annotations obtained using scOMM for each individual dataset. Furthermore, global multimodal integration (mosaic and diagonal) did not outperform other multimodal combinations (Fig. 5A) in cell type identification, as reflected by the comparable cell-type b-cLISI scores in the renal epithelial compartment. This suggests that, depending on the heterogeneity of cell types or states, vertical or horizontal integrations may already offer sufficient resolution for accurate identification. For example, in POD, PEC, and IC subtypes, silhouette and b-cLISI values were comparable or slightly higher in the horizontal RNA and vertical integrations, indicating subtle but meaningful improvements in resolving these cell subtypes. However, in adaptive TAL cells, particularly in aTAL1, global integrations demonstrated better local resolution, as evidenced by increased b-cLISI scores, which improved significantly from 0.45 in vertical integration to 0.61 in the mosaic integration. We hypothesized that this adaptive state is better resolved in 3’ scRNA-seq datasets, displaying higher b-cLISI score in the unimodal integration (b-cLISI=0.55, Fig. 5A), suggesting that 3’ scRNA-seq captures biological variations that are less effectively represented in other modalities. Therefore, we re-clustered the TAL population and identified a subtype of aTAL1, referred to as aTAL1_0, which was predominantly enriched in cells from the 3’ scRNA-seq data (Fig. 5B). By analyzing genes enriched in this cluster, we found *WFDC2* to be the top marker for this population (Fig. 5C, Supp. Fig. 13A). *WFDC2* was also expressed in aTAL2 and MD cells, though at lower levels in both 3’ and 5’ scRNA-seq, but it was absent in snRNA-seq data in these populations (Supp. Fig. 13B). The identification of this *WFDC2*-marked subpopulations is intriguing, as *WFDC2*, which encodes Human Epididymis Protein 4 (HE4), has been proposed as a serum biomarker for lupus nephritis and chronic kidney disease^30^. Recent studies also suggest that *WFDC2* is a suitable pan-distal nephron marker in the human kidney^31^. We further compared additional markers of the aTAL1_0 population to better characterize this subpopulation with respect to the other aTAL1 groups, aTAL1_1 and aTAL1_2 (Fig. 5B). The differential detection of markers across these subtypes and protocols (Supp. Fig. 13C) suggests underlying biological, and potentially functional, differences rather than solely technical biases. *WFDC2* and *B2M,* top markers of aTAL1_0, are often linked to active immune or stress responses^30,32^.

**Figure 5.**
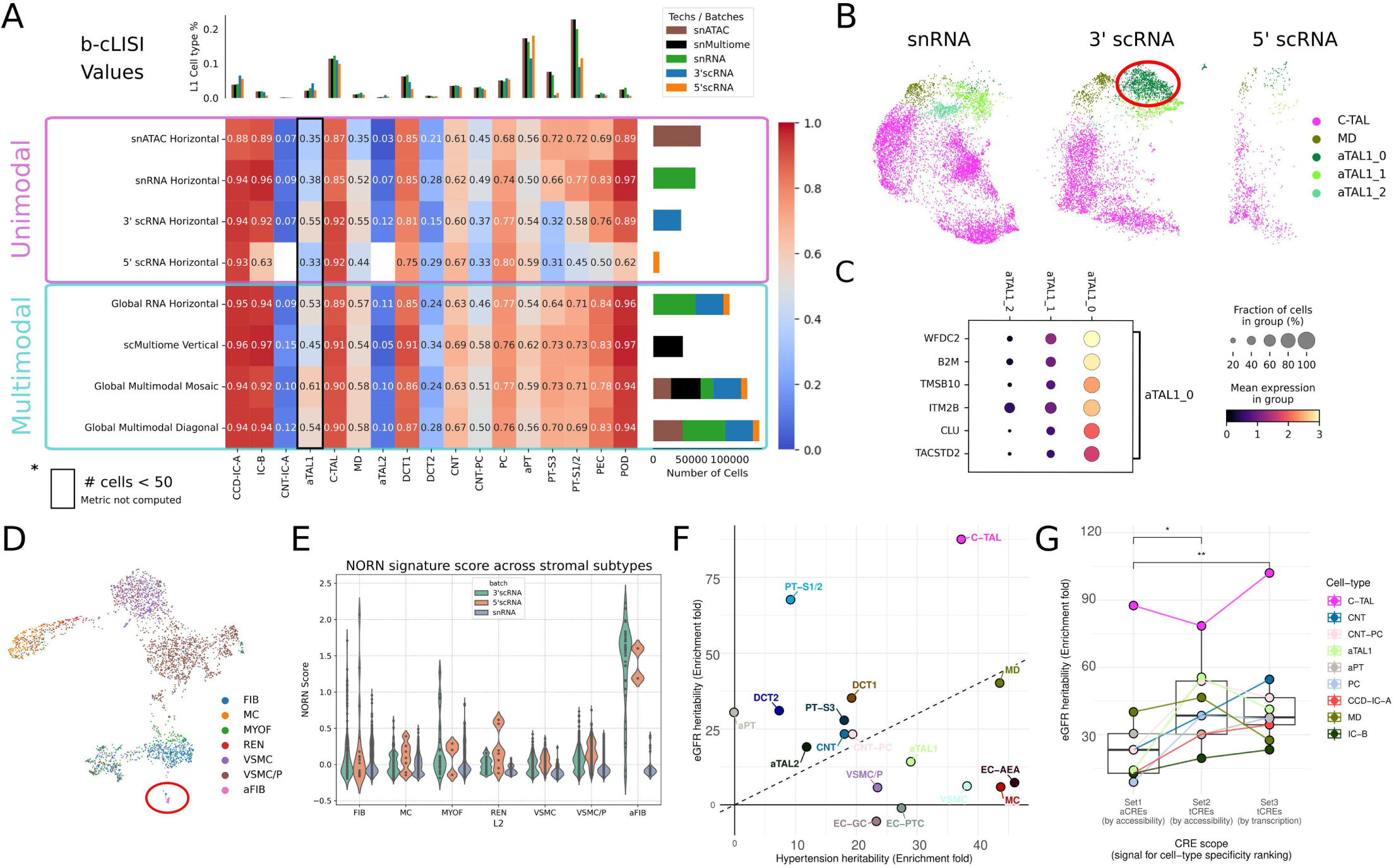
Global multimodal integration of paired and unpaired sn/scRNA/ATAC data. A) Heatmap showing the comparison of binary cell type LISI scores (L2 level) across unimodal (pink) and multimodal (blue) integration strategies for different kidney cell types. Each cell’s value represents the accuracy of cell type annotation, with red indicating higher accuracy and blue indicating lower accuracy. Bar plots at the top show the relative number of L1 cells per cell type. Cell types are listed at the bottom, and the right side provides a color key for the corresponding modalities and cell types. B) UMAP plots showing the re-clustering of the TAL1 subtype within the TAL compartment, split by snRNA, 3’ scRNA, and 5’ scRNA modalities. Notably, aTAL1_0 is enriched in the scRNA-seq protocols (3’ scRNA and 5’ scRNA), while aTAL1_2 is more specific to the snRNA-seq dataset, highlighting modality-specific differences in cell subpopulation detection. C) Dot plot illustrating the expression levels of key marker genes (*WFDC2*, *B2M*, *TMSB10*, *ITM2B*, *CLU*, *TACSTD2*) across aTAL1 subpopulations. Dot size represents the fraction of cells expressing the gene, and color intensity indicates log-normalized mean expression on each group. D) UMAP plot showing the re-clustering of stromal compartment cell types, including FIB (Fibroblasts), MC (Mesangial Cells), MYOF (Myofibroblasts), REN (Renin-producing cells), VSMC (Vascular Smooth Muscle Cells), and others. The red circle highlights the rare population of aFIB cells identified as the Norn cells. E) Violin plots showing the Norn signature scores across stromal subtypes splitter by sn/scRNA-seq protocols. F) Enrichment of hypertension and eGFR heritability in accessibility-defined cell type-specific CREs. Set 1 cell type-specific CREs were used for heritability enrichment analyses, and only cell types with an enrichment p-value < 0.1 for either trait are displayed. G) Comparison of cell type-specific CREs defined by accessibility and transcription. Only cell types present in all three sets and showing an enrichment p-value < 0.05 in at least one set are included. Asterisks indicate significance based on the Wilcoxon rank-sum test, whith * corresponding to p-values < 0.05 and ** corresponding to p-values < 0.01.

Applying a similar approach to the stromal compartment, b-cLISI scores highlighted higher values in the 3’ scRNA-seq data for the adaptive state of fibroblasts (aFIB; Supp. Fig. 12D), where reclustering of the stromal cells identified a rare subpopulation of erythropoietin (EPO)- producing cells, known as Norn cells^33^, characterized by the expression of DCN, TIMP1, and CFD (Fig. 5D). Typically, EPO is produced by peritubular fibroblast-like cells in the kidney, which respond to low oxygen levels by increasing EPO production to stimulate red blood cell generation, a process crucial for maintaining oxygen delivery throughout the body^34^. Norn cells were predominantly detected in 3’ and 5’ scRNA-seq datasets (Fig. 5E) but were not observed in the same samples analyzed by snRNA-seq, likely due to differences in method sensitivity. Identifying EPO-producing cells is especially relevant in the context of kidney disorders, where chronic conditions such as renal hypoxia can impair EPO production, leading to anemia. Understanding these cells’ behavior and regulation could provide valuable insights into treating anemia associated with chronic kidney disease and other conditions characterized by reduced oxygen levels. The differences in marker detection across these protocols highlight the biological complexity of these subpopulations, with each technology offering complementary insights into their cellular states.

### Integrative multimodal analysis reveals enhanced trait heritability in kidney cell-type-specific cis-regulatory elements

To further understand the impact of multimodal analysis in uncovering the molecular mechanisms driving cellular function and disease states, we focused on gene regulatory networks (Supplementary Material) and cis-regulatory elements (CREs) and their enrichment in Genome-wide association studies (GWAS) traits. GWAS have identified genetic variants associated with traits and diseases^35^, many of which are enriched within CREs, highlighting their critical role in gene regulation and trait heritability. Chromatin accessibility assays are commonly used to identify accessible CREs^36^. However, many distal CREs lack epigenomic features characteristic of active enhancers^37^. While some of these elements may function as insulators^38^ or silencers^39^, their broader roles in gene regulation remain poorly understood, making it challenging to annotate trait-associated variants based solely on chromatin accessibility. Emerging evidence indicates that transcriptional activity at distal CREs, as identified through 5′-end RNA-seq methods^40^, can serve as a marker of enhancer activity^41^. This suggests that transcriptional activity may offer a more informative and interpretable metric than chromatin accessibility for functional annotation of trait-associated variants. To demonstrate this, we systematically compared chromatin accessibility and transcription as metrics for assessing trait heritability in kidney cell-type-specific CREs. Specifically, we assessed heritability enrichment for hypertension^42^ and estimated glomerular filtration rate (eGFR)^43^ using accessibility-defined, cell-type-specific CREs (Fig. 5F; Table 6; see Methods). MD cells exhibited strong heritability enrichment for both traits, consistent with their role in regulating glomerular filtration rate and systemic blood pressure through the renin-angiotensin-aldosterone system^44^. Proximal tubular cells (PT-S1/2 and PT-S3) also showed enrichment for both traits, while adaptive proximal tubular cells (aPT) were enriched only for eGFR. This may reflect that genes activated in adaptive states (a potential failed-repair population)^5^ are enriched in eGFR-associated loci. Endothelial cells (EC-PTC, EC-GC, EC-AEA) and vascular smooth muscle/pericytes (VSMC, VSMC/P, MC) were enriched for hypertension heritability but not eGFR, emphasizing the role of blood vessels in hypertension^45^. These results imply that cell-type-specific CREs defined by accessibility are critical for interpreting cell-type-specific heritability enrichment. Next, we compared the utility of accessibility and transcription signals in defining cell-type-specific CREs for heritability enrichment (see Methods; Table 5). We found that either incorporating transcription signals in CRE selection (Set 2) or ranking CREs by transcription (Set 3) yielded significantly higher heritability enrichment (Fig. 5G, Wilcoxon rank-sum test, p < 0.05) compared to using accessibility alone (Set 1). This aligns with evidence suggesting that transcribed enhancers are more likely to validate in functional assays compared to epigenetically defined enhancers^41^. Altogether, these results highlight the advantage of integrating accessibility and transcription signals to enhance the sensitivity and interpretability of trait heritability analyses.

## Discussion

To fully harness the potential of multimodal data integration for understanding cellular and tissue function, benchmarking experimental methods and computational tools is crucial. Setting reliable standards helps ensure accuracy and shows the strengths and weaknesses of each modality. In this study, we investigated the integration of distinct single-cell protocols and multimodal data to create a detailed human organ reference focused on the kidney’s cortex. We aimed to address technical biases, improve tissue characterization, and assess how each single-cell data type contributes to identify cell types and states in both matched and unmatched samples. We also evaluated their complementarity and robustness, along with the reproducibility of data in samples from the same donors. The resulting mBDRC serves as a detailed reference for this complex organ, capturing a range of heterogeneous populations, from well-defined, homogeneous cell types to continuous, challenging-to-distinguish states. This diversity enabled us to explore key features relevant to cell type identification and marker detection across multiple integrative scenarios by using two complementary classes of approaches: supervised (i.e., label transfer-based, via scOMM) and unsupervised (i.e., graph embedding-based) methods (Fig. 6). Not all integration strategies were evaluated using both label transfer-based and clustering-based approaches. For instance, diagonal and mosaic integrations were only assessed with embedding-based methods, reflecting methodological constraints and the differing applicability of evaluation metrics across integration scenarios. In embedding-based approaches, both protocol-specific and multimodal integrations consistently demonstrated a clear positive impact on cell type identification, with RNA modalities outperforming ATAC in most cell compartments. Among the integration strategies, mosaic integration (i.e., including paired or unpaired data types) achieved the highest overall scores, while diagonal (i.e., all data are used but multiomics data are used as unpaired data) and horizontal integrations (same data type across samples) outperformed vertical integration (different data types within the same samples). Supervised classification using scOMM further emphasized the critical role of external references for accurate cell type prediction, as evidenced by the higher performance scores achieved for cell type identification compared to embedding-based approaches. This underscores the importance of reference-guided annotation in achieving robust and consistent cell type classifications. In this regard, the integration of diverse sn/scRNA-seq data demonstrated significant advantages in precisely identifying cell states, as different protocols contribute in varying ways to this process. For instance, snRNA-seq data provided higher resolution for proximal tubules (i.e., PT-S1/2, aPT, PT-S3), 3’ scRNA-seq excelled in resolving TAL subtypes, while 5’ scRNA-seq data were more effective in identifying smaller subpopulations, such as MD cells and subtypes within distal convoluted tubules and collecting ducts (Fig. 3C). Employing both horizontal and vertical integration strategies substantially improved data resolution, allowing for more accurate cellular mapping (Fig. 4C). Methods like multi-spectral integration preserved unique information from both RNA and chromatin accessibility data (Fig. 4B-C), offering refined insights into specific cell subtypes, such as podocytes (Fig. 4C, Fig. 5A), which exhibit distinct transcriptional and epigenetic profiles that make them particularly well-suited to benefit from multimodal integration.

**Figure 6.**
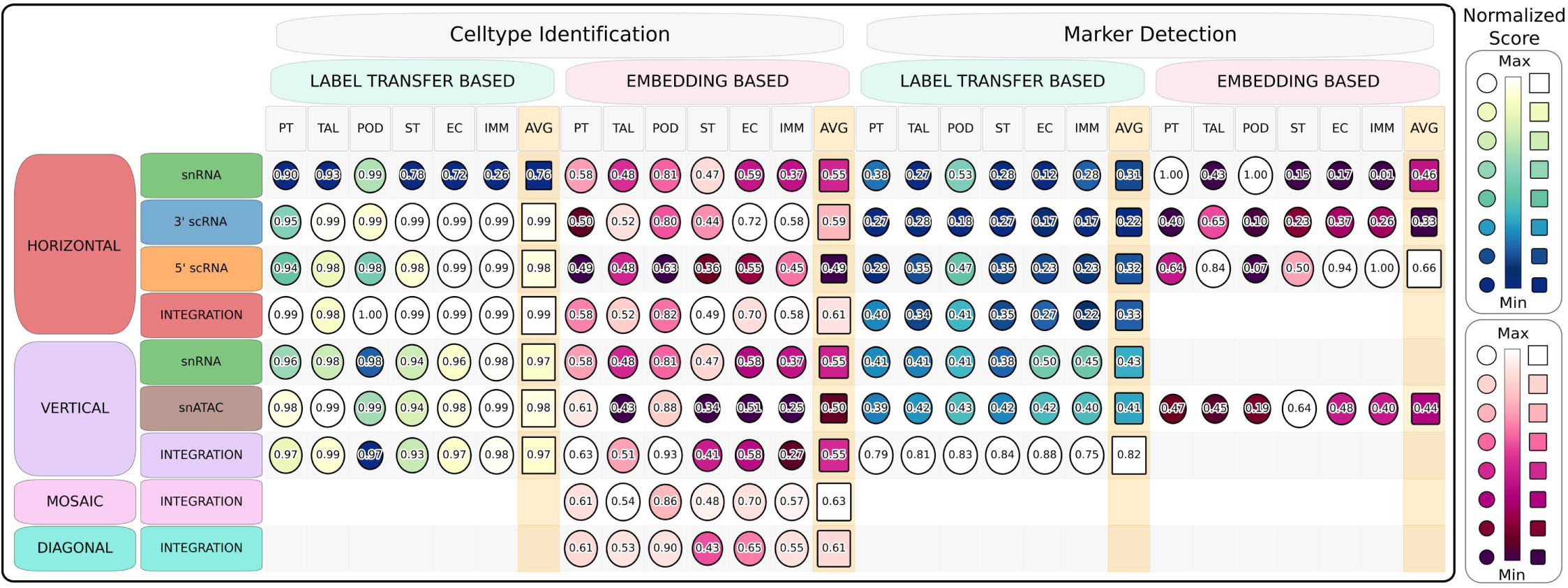
Benchmarking summary of the integrative scenarios for distinct single-cell data types. Different unimodal and multimodal integration strategies were evaluated based on customized scores (see Methods) for cell type identification and marker detection based on supervised (transfer learning-based via scOMM) and unsupervised (embedding-based) approaches. Performance scores are displayed for 6 kidney populations/compartments: PT (Proximal Tubule), TAL (Thick Ascending Limb), POD (Podocytes), ST (Stromal), EC (Endothelial Cells), and IMM (Immune Cells). AVG (Average) represents the arithmetic mean across shown cell groups in each evaluated approach and integration scenario. Shapes (i.e., circles for individual cell types and squares for the average across cell groups) and colors display columns with normalized scores, making the results comparable across different types of integration, while values are the original performance scores before normalization.

A key outcome of multimodal global integration was its ability to detect rare and clinically significant cell populations. For instance, we identified a unique TAL cell subpopulation expressing *WFDC2*, previously suggested as a lupus biomarker, detectable only in scRNA-seq data (Fig. 5B-C, Supp. Fig. 13A). Notably, we uncovered the very rare erythropoietin-producing Norn cells within the adaptive fibroblast population (Fig. 5D-E), a finding with potential therapeutic relevance for addressing anemia in kidney disease. These results clearly show the potential of combining distinct protocols to identify functionally and clinically relevant cell types that might be missed using a single modality.

Beyond cell type identification, integrating transcriptional and chromatin accessibility data revealed key regulatory mechanisms underlying trait heritability. We showed that transcriptional activity at CREs enriched in GWAS-associated traits more accurately reflects the functional relevance of trait-associated variants than chromatin accessibility alone (Fig. 5G). This was particularly evident for hypertension and eGFR, where heritability was enriched in CREs specific to proximal tubular, endothelial, and vascular smooth muscle cells. Overall, this work provides a systematic framework for creating comprehensive tissue atlases, establishing a valuable precedent for multimodal approaches that enable biologically meaningful insights into complex organs. The integration methodologies applied here demonstrate that multiomics data integration can significantly enhance the resolution and accuracy of cell type identification. Additionally, we are confident that our mBDRC will serve as an important resource for the scientific community, advancing both kidney research and next-generation protocols and tools for large-scale single-cell studies, such as those in the Human Cell Atlas.

### Study limitations

This study establishes a robust framework for integrating multimodal single-cell data and evaluating their individual and combined contribution to kidney cell biology. However, there are limitations that need to be considered. Some experimental protocols, including Smart-seq2 and scNMT-seq were not fully optimized for frozen kidney tissue, potentially affecting data quality and restricting their use to exploratory analyses. Similarly, we were unable to generate any useful data with MERFISH even with multiple attempts to optimize this method for our samples (unpublished data). The limited number of matched samples across protocols also introduces variability from donor heterogeneity, which could confound computational analyses such as gene regulatory network inference and transcription factor identification, detailed in the Supplementary material. Moreover, the exclusive focus on kidney samples could limit the extent of the generalizability of these findings to other tissues with distinct cellular compositions and regulatory mechanisms. On the other hand, computational challenges remain, particularly due to the lack of a ground truth for cell-type identification and marker detection, which limits the precision of evaluations. We minimized this bias as much as possible by consulting kidney experts for annotations and simulating realistic analysis scenarios to create a robust framework for evaluating cell type prediction performances.

## Authors contribution

E.M. and J.Z.L. coordinated the overall study. E.M., J.Z.L., H.H., P.C., W.E., and T.V. designed the original project, secured funding, and supervised the project, with contributions from E.F., M.B., and M.Kretzler. C.L.O. procured and processed human kidney tissue and performed pathological assessments. E.A.O. prepared human kidney tissue for scRNA-seq and oversaw quality control of its data. Single-cell experiments were conducted by X.A., D.M., G.C., R.M., T.L., P.T., and M.Kojima. Y.A. and K.V. coordinated parts of the study. E.M. conceived and guided the analysis framework for the study. M.A.-M. and J.-K.L. performed most of the computational analyses, with contributions from S.K.S., C.-C.H.H., B.V., M.A.M., R.M., T.L., and I.C. R.M., M.B., and M.Kretzler contributed to data interpretation. All authors contributed to writing the manuscript, read, and approved the final version.

## Data and code availability

All data analysis code used in this study is available at https://github.com/mereulab/Multimodal_single-cell_Benchmarking. The integrated multimodal single-cell data has been deposited and will be accessible via the CELLxGENE platform. ScOMM is publicly available as an R package at https://github.com/mereulab/scOMM. The empirical power analysis is available at https://github.com/bvieth. Raw sequencing data, including FASTQ files for gene expression datasets (scRNA-seq and snRNA-seq), will be released on GEO. Additional datasets, including chromatin accessibility (snATAC-seq) and multiome data, will be made available through their respective repositories upon publication.

## Acknowledgements.

We thank Felix Eichinger for analysis of bulk RNA-seq data, the Broad Institute Flow Cytometry group for cell sorting, the Broad Institute Genomics Platform for sequencing, MIT SuperCloud and Lincoln Laboratory Supercomputing Center for providing HPC resources that contributed to the research results reported, Cristin McCabe and Nathan Haywood for technical support, Anna Greka and Menna Clatworthy for guidance at early stages of the project, all members of the Enard/Hellmann lab for valuable input and support during initial stages of the project in particular Ines Hellmann, Zane Kliesmete and Philipp Janssen. We also acknowledge both current and former members of Mereu’s lab for their continuous feedback and valuable discussions throughout the duration of the project.

## Online Methods

### Human Tissue Procurement

Fresh normal tissue from the unaffected part of surgically removed kidney of patients undergoing total nephrectomy at the University of Michigan was obtained directly from the operating theater at Michigan Medicine. Patients were enrolled in the PRECISE cohort at the University of Michigan. The PRECISE study was approved by the institutional review board (IRB) of the University of Michigan (HUM00165536, HUM00052918) as previously described^46^. Patient data were obtained from each participant’s electronic medical record. Procured tissue specimens were a minimum of 1.5 cm x 1.0 cm x 1.0 cm and included all the kidney compartments (capsule, cortex, and medulla). In order to minimize cold ischemic time, tissue was obtained directly from the operating room and immediately processed on site using a 3D printed device (“PRECISE Pyramid”). Utilizing this device, nephrectomy specimens were simultaneously cut into approximately 25 cores of equal dimension that were closely related in space and resemble 16-gauge clinical kidney biopsy cores (Supp. Fig. 1). Biopsy cores were subsequently preserved across a variety of media, including CryoStor, RNALater, OCT, and LN_2_^47,48^. Biopsy cores were stored for later use at −80°C and shipped to interrogation sites on dry ice.

### 3’ scRNA-seq

We utilized 3’ scRNA-seq data sets from tumor-free kidney cortical tissue of nephrectomy specimens within the PRECISE cohort at the University of Michigan (IRB: HUM165536). Detailed tissue processing, single-cell isolation, and scRNA-seq generation protocols are available in the KPMP scRNA-Seq protocol (https://www.protocols.io/view/single-cell-rna-sequencing-scrna-seq-7dthi6n). Briefly, kidney biopsies preserved in CryoStor and frozen in liquid nitrogen were rapidly thawed and enzymatically dissociated into a single-cell solution using Liberase TL digestion for 12 minutes at 37°C. The solution was then filtered through a 30 μm strainer (Miltenyi Biotec) and washed in DMEM/F12 medium supplemented with 10% fetal calf serum and HEPES buffer. The cells (average 40,000 cells) were processed using the droplet-based platform with Chromium Single Cell 3’chemistry (v3.1, 10x Genomics). The cDNA libraries were prepared and sequenced on an Illumina NovaSeq 6000 platform, generating over 200 million reads (average 25,000 reads per cell) per sample using one of the following two conditions: 28 bases (Read 1), 10 bases (Index read), and 151 bases (Read 2) or 151 bases (Read 1), 8 bases (Index read), and 151 bases (Read 2) – see Table 2 for details. The Cell Ranger (v3, 10x Genomics) pipeline was employed to extract the cell x gene matrix from FASTQ files aligned to the GRCh38 genome reference (version 2020-A).

### Tissue dissociation and single-cell plate sorting

In this section, we describe the preparation of samples for 5’ scRNA-seq, Smart-seq2, and scNMT-seq. We placed the cryopreserved tissue containing tube in a 37°C water bath for 1 minute for quick thawing. We transferred the tissue to a plastic petri dish (diameter 5 cm) filled with 1 ml DMEM/F12/10% fetal bovine serum (FBS) prepared with 90% DMEM/F12/HEPES (Thermo Fisher Scientific, 11330057) and 10% heat inactivated FBS (Thermo Fisher Scientific, 16140-071) for 10 seconds to wash off the remaining DMSO at room temperature, followed by transferring the tissue to a new petri dish containing 1 ml DMEM/F12/10% FBS and incubating for 10 minutes at room temperature. We prepared 1 ml dissociation media by mixing 900 µl DMEM/F12 and 100 µl Liberase TL (Millipore Sigma, 5401020001, 2.5 mg/ml in H_2_O). We then minced the tissue in another new petri dish filled with 500 µl dissociation media for 1 minute (or 2 minutes for 5’ scRNA-seq) with a razor blade. We transferred the minced tissue to a 1.5 ml LoBind Eppendorf tube and rinsed the petri dish with the remaining 500 µl media and collected to the same tube. We incubated the tube at 37°C in a thermomixer for 12 minutes at 500 rpm, during which we triturated after 6 minutes 15 times with a wide bore 1 ml pipette tip. We stopped the reaction by adding 1 ml (or 500 µl for 5’ scRNA-seq) DMEM/F12/10% FBS (room temperature) and gentle mixing and incubating at room temperature for 1 minute. We passed the dissociated tissue through a 30 µm filter (Sysmex, 04- 004-2326 or Miltenyi Biotec, 130-098-458 for 5’ scRNA-seq) and into a 15 ml tube on ice. We rinsed the filter with 10 ml cold DMEM/F12/10% FBS. We then filtered the flowthrough again through a new 30 µm filter and collected into a new 15 ml tube. We rinsed the previous 15 ml tube with 1 ml cold DMEM/F12/10% FBS and passed through the second filter and collected into the same tube. We spun down the collection for 10 minutes at 200 g and 4°C. After supernatant removal, we resuspended the pellet with 200 µl (or 40 µl for 5’ scRNA-seq) DMEM/F12/10% FBS.

For 5’ scRNA-seq, we added 1 ml PBS/1% UltraPure BSA (Thermo Fisher Scientific, AM2616) and transferred the cells to a new 1.5 ml tube. We centrifuged for 10 minutes at 100 g at 4°C. After supernatant removal, we resuspended the pellet with 400 µl PBS/1% BSA. We then filtered the cells through a 20 µm filter (pluriSelect, 43-10020-40) and collected into a new 1.5 ml tube.

For Smart-seq2 and scNMT-seq, after cell counting with a Cellometer K2 (Nexcelom Bioscience) and AOPI stain (Nexcelom Bioscience, CS2-0106-5ML), we diluted the cell suspension to between 1 to 10 cells/µl with Phosphate Buffered Saline (PBS) containing 0.04% BSA (Miltenyi Biotec, 130-091-376) and 0.2 U/µl Recombinant RNase Inhibitor (Takara, 2313A). We stained the cells with 2 µl Propidium Iodide (PI, BioLegend, 421301) and 1 µl (1:10) CalceinAM (LifeTechnologies, C3099) per ml of cell suspension. To isolate single cells for the Smart-seq2 assay, we used a Hana cell sorter (Namocell) and gated on PE^-^/FITC^+^ to exclude dead cells and debris and sorted live single cells into 96-well plates with each well preloaded with 1 µl lysis buffer (Takara, 635013) containing 0.4 U Recombinant RNase Inhibitor. For the scNMT-seq assay, we sorted the above single cells into another set of special 96-well plates (Thomas Scientific, 1149V59 (4ti-0970/C)), in which each well was preloaded with 1.5 µl reaction buffer containing 0.25 µl 10x M.CviPI reaction buffer (New England Biolabs, M0227L), 2 U of GpC methyltransferase M.CviPI (New England Biolabs, M0227L), 0.5 µl of 800 µM SAM (New England Biolabs, M0227L), 0.025 µl 10% IGEPAL (Sigma, I8896-50ml), 2.5 U of Recombinant RNase Inhibitor. We spun down the Smart-seq2 plates at 2000 g for 2 minutes at 4°C and stored them at −80°C for later processing. We also spun down the scNMT-seq plates with the same conditions before a 37°C incubation for 15 minutes followed by adding 5 µl RLT buffer (Qiagen, 1053393) and freezing at −80°C.

### 5’ scRNA-seq

We did a final counting with Trypan Blue and loaded aiming to recover 10,000 cells per sample preparation on the Chromium Next GEM Chip G (10x Genomics, 1000127). We performed the subsequent library preparation with a Chromium Next GEM Single Cell 5’ v1.1 kit (10x Genomics, 1000167) following the vendor protocol with single indexing for the 5’ gene expression libraries. The 5’ gene expression libraries were sequenced on a NovaSeq6000 S2 200 cycle flow cell (Illumina) with 111 bases for read 1, 91 bases for read 2 and 8 bases for index 1.

### Smart-seq2

For four samples (lib_34, lib_36, lib_29, lib_676), we prepared cDNA using the SMART-Seq Single Cell Kit (Takara, 634471) following the vendor manual with 1/5x reduced reaction volumes and purified the cDNA with AMPureXP beads (Beckmann Coulter, A63881) with a 0.8x volume ratio. We quantified cDNA with a Wallac EnVision plate reader and normalized concentrations across all wells. We used 0.075 ng cDNA for each and prepared Illumina sequencing libraries with the NexteraXT kit (Illumina, FC-131-1096) with a 1/4x reduced reaction volume. We pooled individual libraries with equal volumes and sequenced on a NextSeq 500 flow cell (Illumina) with 50 bases for read 1, 25 bases for read 2, and 8 bases each for index 1 and index 2, and aimed for about 1 million reads per cell.

For one (lib_34) of those four samples, full-length single-cell RNA-seq libraries were also prepared separately using the SMART-Seq v5 Ultra Low Input RNA Kit for Sequencing (Takara Bio). All reactions were downscaled to one quarter of the original protocol and performed following the manufacturer’s thermal cycling conditions. Briefly, reverse transcription was performed using 2.5 µl of the RT MasterMix (SMART-Seq v5 Ultra Low Input RNA Kit for Sequencing, Takara Bio). cDNA was amplified using 8 µl of the PCR MasterMix (SMART-Seq v5 Ultra Low Input RNA Kit for Sequencing, Takara Bio) with 23 cycles of amplification. Following purification with Agencourt Ampure XP beads (Beckmann Coulter), product size distribution and quantity were assessed on a Bioanalyzer using a High Sensitivity DNA Kit (Agilent Technologies). A total of 140 pg of the amplified cDNA was fragmented using Nextera XT (Illumina) and amplified with double indexed Nextera PCR primers (IDT). Products of each well of the 96-well plate were pooled and purified twice with Agencourt Ampure XP beads (Beckmann Coulter). Final libraries were quantified and checked for fragment size distribution using a Bioanalyzer High Sensitivity DNA Kit (Agilent Technologies). Pooled sequencing of Nextera libraries was carried out using a NovaSeq 6000 (Illumina) to an average sequencing depth of 0.5 million reads per cell. Sequencing was carried out as paired-end (PE150) reads with library indexes corresponding to cell barcode.

### Nuclei isolation and single-cell multiome sequencing

A) For eight of the samples (lib_09, lib_10, lib_23, lib_29, lib_51, lib_54, lib_56, lib_57), we isolated nuclei from snap frozen human kidney issue based on a vendor (10x Genomics, CG000366_DemonstratedProtocol_SingleCellMultiome_Nuclei_EmbMouseBrain_Rev) provided protocol with these modifications. Briefly, we chopped the tissue in 0.75 ml cold lysis buffer containing 10 mM Tris-HCl pH 7.4, 10 mM NaCl, 3 mM MgCl_2_, 0.01% Tween-20 (Sigma, 11332465001), 0.01% NP40 (Sigma, 11332473001), 0.001% Digitonin, 1% BSA (Miltenyi Biotec, 130-091-376), 1mM DTT and 1 U/µl Recombinant RNase Inhibitor in a 1.5 ml Eppendorf tube with surgical scissors for about 3 minutes. We then transferred the chopped tissue together with 1.25 ml rinse with lysis buffer into a 2 ml dounce tissue grinder tube (Sigma, T2690). We ground the tissue with pestle A for about 15 strokes until resistance went away followed by 15 strokes with pestle B until smooth. We passed the tissue lysate through a 50 µm filter (Sysmex, 04-004-2327) together with another 2 ml tube rinse with the lysis buffer. We then passed the flowthrough through another 35 µm blue cap filter (Thermo Fisher Scientific, 08-771-23). We spun down the flowthrough at 500 g for 5 minutes at 4°C and resuspended the nuclei pellet in 1 ml cold wash buffer containing 10 mM Tris-HCl pH 7.4, 10 mM NaCl, 3 mM MgCl_2_, 0.1% Tween-20, 1% BSA, 1 mM DTT and 0.04 U/µl Recombinant RNase Inhibitor and repeated the centrifugation one more time. We resuspended the pellet in 1 ml cold wash buffer and counted the nuclei using the Cellometer K2 with the AO stain. We spun down the nuclei suspension at 500 g for 5 minutes at 4°C and resuspended the pellet with the diluted nuclei buffer containing 1x Nuclei Buffer (10x Genomics, 2000207), 1 mM DTT and 1 U/µl Recombinant RNase Inhibitor in a volume based on the previous counting and aiming for around 4,840 nuclei/µl. We did a final counting and loaded aiming to recover 6,000 nuclei per sample preparation.

We performed the tagmentation reaction followed by loading on the Chromium Next GEM Chip J (10x Genomics, 1000230) and subsequent library preparation with Chromium Next GEM Single Cell Multiome ATAC + Gene Expression kit (10x Genomics, 1000285) following the vendor protocol with single indexing for the gene expression libraries. We sequenced the ATAC-seq libraries on a NovaSeq 6000 SP 100 cycle flow cell (Illumina) with 34 bases each for read 1 and read 2, 8 bases for index 1 and 24 bases for index 2 loading together with 1% PhiX. We sequenced the gene expression libraries on a separate NovaSeq 6000 SP 100 cycle flow cell with 28 bases for read 1, 55 bases for read 2 and 8 bases for index 1.

B) For five of the samples (lib_15, lib_34, lib_36, lib_38, lib_55), nuclei were isolated from snap-frozen human kidney samples by following the Demonstrated Protocol for Single Cell Multiome Nuclei Isolation from Embryonic Mouse Brain (10x Genomics, CG000366) with the following modifications. Briefly, frozen tissues were transferred from –80°C to a petri dish on dry ice and cut into 2–3 smaller pieces. The samples were placed in a dounce homogenizer with 2 ml of 0.1x Lysis Buffer containing 10 mM Tris-HCl (pH 7.4), 10 mM NaCl (Invitrogen, AM9759), 3 mM MgCl₂ (Invitrogen, AM9530G), 0.01% Tween-20 (Sigma, 11332465001), 0.01% NP40 (Sigma, 11332473001), 0.001% Digitonin (Invitrogen, 10636033), 1% BSA (Miltenyi Biotec, 130-091-376), 1 mM DTT (Sigma, 646563-10X), and 1 U/µl RNase Inhibitor (Roche, 03335402001). The tissue was disaggregated with 30 strokes (approximately 15 strokes per pestle), then transferred to a 2 ml Eppendorf tube and incubated on ice for 5 minutes. A 70 µm strainer was pre-wetted with 0.5 ml of wash buffer containing 10 mM Tris-HCl, 10 mM NaCl, 3 mM MgCl₂, 0.1% Tween-20, 1% BSA, 1 mM DTT, and 1 U/µl RNase Inhibitor. The tissue lysate was then passed through the filter, and 1.5 ml of wash buffer was added to rinse the filter. The sample was centrifuged at 550 g for 5 minutes at 4°C using a swinging bucket rotor to pellet the nuclei. The pellet was resuspended in 1 ml of cold wash buffer, and the nuclei were counted using a TC20™ Automated Cell Counter (Bio-Rad) after staining with Trypan Blue (Gibco, 15250-061). Nuclei were washed a total of three times and finally resuspended in the appropriate volume of chilled Diluted Nuclei Buffer (10x Genomics) supplemented with 1 mM DTT and 1 U/µl of RNase Inhibitor to achieve a nuclei concentration suitable for a target recovery of 5000–7000 nuclei. If large clumps or debris were observed in the final suspension, the nuclei were additionally filtered with a 40 µm Flowmi Cell Strainer (Bel-ART, H13680-0040), before staining with Trypan Blue and performing final manual counting using a Neubauer chamber.

Nuclei transposition and library preparation were performed following the Chromium Next GEM Single Cell Multiome ATAC + Gene Expression User Guide (10x Genomics, CG000338). Transposed nuclei were partitioned into GEMs using the Chromium Controller with Chip J, aiming for a target recovery of 7000 nuclei per sample. After GEM incubation for mRNA reverse transcription and transposed DNA barcoding, the resulting cDNA and barcoded gDNA were purified and pre-amplified with 7 cycles, following the 10x Genomics protocol. After clean-up, 35 µl of the pre-amplified cDNA was amplified with 7 additional PCR cycles. The resulting cDNA was quantified on an Agilent Bioanalyzer High Sensitivity chip (Agilent Technologies), and 100 ng was used for library preparation. GEX libraries were indexed with 13 cycles of amplification using the Dual Index Plate TT Set A (10x Genomics; PN-3000431). In parallel, 40 µl of the pre-amplified DNA was indexed with 7 cycles of amplification using the Sample Index N Set A (10x Genomics; PN 3000427). The size distribution and concentration of full-length GEX and ATAC-seq libraries were verified on an Agilent Bioanalyzer High Sensitivity chip. Finally, sequencing of GEX libraries was carried out on a NovaSeq 6000 sequencer (Illumina) using the following sequencing conditions: 28 bases (Read 1), 8 bases (i7 index), 8 bases (i5 index), and 90 bases (Read 2), to obtain approximately 40,000 paired-end reads per nucleus. The ATAC-seq libraries were also sequenced on a NovaSeq 6000 sequencer using the following conditions: 50 bases (Read 1), 8 bases (i7 Index), 16 bases (i5 Index), and 49 bases (Read 2), aiming for a sequencing depth of >20,000 reads/nucleus.

C) For one sample (lib_34), we also followed a nuclei isolation method indicated for the snDropSeq method, as described in the Kidney Precision Medicine Project Tissue Interrogation Site Manual of Procedures (V7.0 – 2019). Briefly, a piece of tissue was placed in 1 ml of Nuclei Extraction Buffer (NEB) containing 20 mM Tris (pH 8), 320 mM sucrose, 5 mM CaCl₂, 3 mM MgAcetate₂, 0.1 mM EDTA, 0.1% Triton X-100, and 1 U/µl of RNase Inhibitor. The tissue was roughly disaggregated by pipetting 10–15 times using a wide-bore 1000 µl tip. The sample was then transferred to a dounce homogenizer (Sigma, D8938-1SET) and stroked 5 times with the loose pestle and 20 times with the tight pestle before being transferred to a 5 ml tube. An additional 2 ml of NEB buffer was used to rinse the dounce and added to the rest of the sample. After incubation on ice for 10 minutes, the sample was filtered through a 40 µm strainer (Pluristrainer, 43-10040-70). Then, 8 ml of PBS supplemented with 1 mM EDTA and 1 U/µl of RNase Inhibitor was added, and the sample was centrifuged for 10 minutes at 900 g at 4°C. The nuclei pellet was then resuspended in the appropriate volume of 1x NB buffer (10x Genomics), filtered with a Flowmi cell strainer, and stained with Trypan Blue for manual counting using a Neubauer chamber. Once the concentration was determined, nuclei were transposed and loaded onto the Chromium system following the manufacturer’s instructions as described above.

### scNMT-seq: gDNA and mRNA separation

The genomic DNA (gDNA) and mRNA were separated using oligo(dT)_30_VN beads on an automated liquid-handling robotics platform (Hamilton STAR) as described for single cell genome-plus-transcriptome sequencing^49^ with minor modifications. More specifically, three washes instead of two were executed using 15 µl instead of 10 µl G&T wash buffer. After separation, the gDNA plate was centrifuged for 1 min at 1,000 rpm at room temperature and stored at −80 °C until further processing, while the mRNA was immediately subjected to reverse transcription and PCR amplification.

### Modified Smart-seq2 and cDNA library preparation

Following separation, the mRNA was reverse transcribed and PCR amplified with an adapted Smart-seq2 protocol^49^. Single-cell samples were amplified for 24 cycles and subsequently purified with SPRI beads (Beckman Coulter) according to the manufacturer’s instructions at 0.8:1 bead:sample ratio and eluted in 25 µl nuclease-free water. Sample concentrations were measured using Quantifluor (Promega) according to the manufacturer’s instructions and fragment size was assessed by Bioanalyzer (Agilent). The cDNA length ranged from 500 to 2,000 bp, peaking around 1-1.5 kb, with a concentration of approximately 1-2 ng/µl. Libraries were prepared following the Nextera XT (Illumina) library preparation kit according to the manufacturer’s instructions, using quarter volumes. 96 single-cell libraries were pooled together and SPRI-purified according to the manufacturer’s instructions at a 0.65:1 bead:sample ratio and eluted in 50 µl elution buffer (Qiagen). The concentration of the library pool was measured using Qubit (Thermo Fisher Scientific) and the size was measured using a Bioanalyzer (Agilent). Expected library pool concentrations were approximately 30 ng/µl with an average size between 400 and 600 bp and smooth profiles. Libraries with expected profiles were 384-plex equimolarly pooled to 2 nM and 50 base paired-end sequenced on an NextSeq 2000 (Illumina) aiming for 1-2 million reads per cell.

### scBS-seq

The gDNA was first SPRI-purified at a 0.65:1 bead:sample ratio. After adding the beads, the samples were incubated 20 min at room temperature. Next, the plate was spun down and placed on a magnet for 20 min. The supernatant was discarded and the beads were washed twice with 80 % ethanol. Finally, the beads were resuspended in a 10 µl elution buffer (Qiagen). To the resuspended beads, 65 µl CT conversion reaction buffer (Zymo, Methylation-Direct MagPrep kit, D5044) was added and samples were incubated as follows: 8 min at 98°C, 3 hours at 65 °C and at most 20 hours at 4 °C. Bisulfite-converted samples were purified using the Methylation-Direct MagPrep kit according to the manufacturer’s instructions using half volumes. Samples were then subjected to single-cell bisulfite sequencing library preparation as described by Clark et al.^7^ Amplified libraries were pooled together and twice SPRI-purified at 0.8:1 bead:sample ratio and eluted in 50 µl elution buffer (Qiagen). The quality of the library pool was assessed using Qubit and Bioanalyzer (Agilent). The concentrations ranged from 10 to 20 ng/µl with an average fragment length of 400-600 bp. Libraries were 44-plex sequenced on an NextSeq 2000 in 150 base paired-end mode aiming for at least 5 million reads per cell.

### 3’ and 5’ scRNA-seq and snRNA/ATAC-seq multiome raw data processing

FASTQ files per sample from 10x Chromium 3’ scRNA-seq, 5’ scRNA-seq, and snRNA/ATAC-seq multiome protocols were processed using the 10x Genomics pipelines: Cell Ranger (v.5)^50^ for scRNA-seq data and Cell Ranger Arc (v.2) for multiome data. Both analyses utilized the GRCh38 (v. 2020-A) reference genome (hg38). Data processing included barcode processing, alignment to the reference genome, and quantification of gene expression for scRNA-seq data using standard parameters. For multiome data, processing also included quantification of accessible chromatin and sample aggregation. Specifically, for snATAC-seq data, sample aggregation was performed to identify a shared set of peaks across samples, with depth normalization set to *None*.

### Smart-Seq2 data and scNMT-seq transcriptomics analysis

FASTQ files for Smart-seq2 and scNMT-seq transcriptome data were processed with the Nextflow DSL1^51^ bulk pipeline, available at https://github.com/seanken/BulkIsoform/tree/main/Pipeline/BulkPipeline.strand.nf. Reads were mapped to the reference genome (used the GTF and FASTA file from the Cell Ranger refdata-cellranger-arc-GRCh38-2020-A reference to construct STAR and Salmon references) with STAR v 2.7.9a^52^, with arguments “--outSAMattributes NH HI AS nM --outSAMtype BAM SortedByCoordinate --readFilesCommand zcat --outStd BAM_SortedByCoordinate -- outSAMunmapped Within”. The resulting BAM file was used by PICARD v2.27.5^53^ to extract QC information with CollectRnaSeqMetrics and MarkDuplicates (default settings). Salmon v1.6.0^54^ was run with the arguments “-l A --posBias --seqBias --gcBias –validateMapping” which was then used for downstream analysis. Other tools were run by the pipeline though not used in this manuscript, so are not described.

Transcripts per million from Salmon were loaded into R with tximport v1.18^55^ and processed with Seurat v4.0.0^56^. Cells with <500 genes were filtered out. NormalizeData was used to normalize to TP10K, and variable genes were selected with FindVariableFeatures, both with default arguments. Data were scaled with the ScaleData command with default parameters and RunPCA was run with npcs = 60. Harmony v1.0^19^ was then used with the RunHarmony command with default parameters to correct for lab of origin differences. The UMAP was generated with the RunUMAP command, while the clustering was calculated with the FindNeighbors command followed by FindClusters. Defaults were used for UMAP and clustering, except with reduction=”harmony” and dims=1:25. Cell types were labeled based on marker genes and manual inspection based on the HCA reference atlas^5^.

### scNMT-seq epigenome data analysis

After sequencing and demultiplexing, adapter sequences were trimmed as described by Clark et al.^57^ using cutadapt v2.103. Trimmed sequences were aligned using Bismark v0.23.18 ^58^ in non-directional mode and the methylation state at individual CpG and GpC sites was extracted using Bismark Methylation extractor using the coverage2cytosine script with --NOMe and -- gc options. Endogenous DNA methylation was examined using CpG methylation, while GpC methylation was used to analyse chromatin accessibility. To eliminate potential biases, CpG methylation was assessed only in the ACG and TCG contexts, while GpC methylation was analysed in the GCH context, where H represents A, C, or T. Methylation levels were linked to genomic features, including gene bodies and promoters. Gene bodies were defined as the region from the transcription start site (TSS) to the transcription end site (TES), extended by 15 kb upstream and downstream. Promoter regions were defined as 1,200 bp upstream and 300 bp downstream of the TSS. Gene body and promoter regions were transformed into a percentage-based scale, where detected positions within a feature were normalised according to the length of the feature, taking strand orientation into account. This enabled conversion of specific genomic locations into relative percentages, representing their position within the feature. For each relative position, the average methylation percentage was calculated across overlapping features within a single cell. Subsequently, per-cell methylation values were pooled across all cells, and a best-fit was generated using a loess regression model to visualise the trends within promotor and gene body endogenous CpG methylation (%CpG) and chromatin accessibility (%GpC).

To determine whether accessibility and methylation levels at the transcription start site differed significantly, two nested regression models were compared. Both models employ a restricted cubic spline approach, with 15 knots for %GpC and 14 knots for endogenous %CpG. The number of knots was selected based on the Akaike Information Criterion (AIC)^59^. Model 1, without interaction effect, assumes that the distance (around TSS and gene body) (*X*_1_) and transcription status (*X*_2_, 1 = transcribed, 0 = non-transcribed) have independent effects on the average methylation across different cells (*Y*, measured as %CpG or %GpC methylation). No interaction effect between these variables was considered. The model was the following:

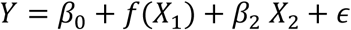

where *f*(*X*_1_) was a restricted cubic spline function describing the nonlinear relationship between distance and methylation, and where 𝜖 were i.i.d. from a normal distribution with mean 0 and residual variance *σ*^2^.

In model 2, with interaction effect, the effect of distance (*X*_1_) on the average methylation across different cells (*Y*) was assumed to depend on transcription status (*X*_2_), and vice versa. This was modelled by including an interaction term between distance and transcription.

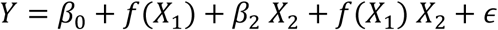

where the term *f*(*X*_1_)*X*_2_ modeled the interaction between distance and transcription, allowing for each transcription type to have its own specific curve. The number of knots for the restricted cubic spline models were determined based on the AIC^60^. The two models were compared using the likelihood ratio test (LRT), to determine whether adding the interaction term in Model 2 provided a significant improvement over Model 1, under the null hypothesis (*H*_0_) that Model 1 was sufficient and the interaction term did not contribute to the model, while the alternative hypothesis (*H*_*A*_) stated that there was a significant interaction term between distance and transcription on methylation levels.

### Quality control procedure for sn/scRNA-seq data

A consistent quality control (QC) procedure was applied across scRNA-seq and snRNA-seq protocols to ensure reproducibility. We began with cell barcodes that passed the 10x Genomics Cell Ranger (or Cell Ranger Arc) filters, followed by additional filtering steps for low-quality cell barcodes and doublet removal, as detailed below.

### Low-quality cell barcode removal

Low-quality cell barcodes were filtered in two stages:

1. Stage 1 - General QC Thresholds:

- Cell barcodes were excluded if they had fewer than 200 or more than 7,500 unique features.
- Mitochondrial content exceeding 70% and ribosomal content exceeding 40% were also excluded.
2. Stage 2 - Cell-Type Specific Filtering:

- Cell-type classifications were informed by scOMM label transfer from the reference data at L1 resolution.
- For immune cells, a mitochondrial content threshold of 10% was applied.
- For non-immune cells: A 20% mitochondrial threshold was applied for cell barcodes with fewer than 500 unique genes.
- Proximal tubule cells were an exception, with a relaxed mitochondrial threshold of 30%.

### Doublet detection

Doublets were identified using the DoubletDetection^61^ software (v.4.2), employing a BoostClassifier model with 30 components and Louvain clustering. Predictions were made on raw counts with a p-value threshold of 1×10^−16^ and a voter threshold of 0.5. Barcodes classified as doublets were excluded.

### Quality control procedure for snATAC-seq data

Similarly, for snATAC-seq data, cell barcodes passing the 10x Genomics Cell Ranger Arc filters were retained as an initial filter before additional low-quality cell barcode and doublet removal, as detailed below.

### Low-quality cell barcode removal

To assess cell barcode quality, metrics specific to snATAC-seq data were applied. Fragment size distribution was used to inspect nucleosome binding patterns and a nucleosome ratio value was computed. Transcriptional start site (TSS) enrichment score was used to confirm the accessibility enrichment around TSS compared to the accessibility on flanking regions. Atac module from Muon^62^ (v0.1.2) was employed to compute was employed to compute QC metrics (muon.atac.tl.nucleosome_signal with n=1e6, muon.atac.tl.tss_enrichment with n_tss=1000). QC threshold:

- Cell barcodes were excluded if they had fewer than 2500 or more than 30000 total features.
- Cell barcodes were excluded if the nucleosome ratio exceeded 4.
- Cell barcodes we filtered if the TSS enrichment score was lower than 2.

### Doublet detection

Doublets were identified using the AMULET^63^ (v1.1) with algorithm standard parameters and specifying the blacklist file for hg38 genome. Barcodes classified as multiplets were filtered out.

### Variance partitioning and cell type trees

For the quantitative comparison of sn/scRNA-seq, eight donors profiled with both technologies were considered (lib_09, lib_10, lib_15, lib_29, lib_36, lib_51, lib_55, lib_56) (Table 2). Broad cell type annotation of scOMM was used. After initial filtering for genes with at least 10% non-zero counts in any cell type and cells with reasonable UMI counts and number of detected genes (i.e. no more than 5 MADs computed with function *isOutlier()* in package scuttle version 1.14.0) per assay, the expression of 22,707 genes in 63,711 cells was kept.

Out of the initial list of eight donors, five donors (lib_09, lib_10, lib_15, lib_51, lib_55) had enough cells across the 14 cell types to achieve a balanced design for the gene-wise partitioning of variance. The 14 cell types were annotated using Muto et al.’s granularity, with alternative names from Lake e.t al.^5^ shown in parentheses: CNT, DCT, ENDO, FIB, ICA (CCD-IC-A), ICB (IC-B), LEUK (IMM), MES (VSM/P), PC, PEC, PODO, PT, TAL, and aPT. For that, pseudobulk samples per cell type, assay and donor were constructed by summing up all counts per cell type with function *aggregateAcrossCells()* in package scuttle (version 1.14.0). The contributions of assay, cell type, interaction of cell type and assay while controlling for donor identity in explaining the variance of normalized gene expression (function *voom()* in limma package version 3.60.4) were estimated with function *fitExtractVarPartModel()* in package variancePartition (version 1.34.0).

The same pseudobulk samples were used to draw dendrograms of cell types per assay based on hierarchical clustering using Euclidean distances and complete agglomeration method with function *buildClusterTreeFromPB()* in package dreamlet (version 1.2.1). The assay-specific dendrograms were compared using stepwise greedy forward selection algorithm by rotating the dendrograms until a local optimal solution of entanglement score (0 = no entanglement, i.e. similar; 1 = full entanglement, i.e. dissimilar) was found (function *untangle(method = “step2side”)* in package dendextend^64^ version 1.19.0).

### Metrics and statistics per cell type and assay

The following metrics were estimated per library: mean, variance and detection rate of gene expression as well as cellular detection rate (CDR, i.e. proportion of genes with nonzero expression per cell), purity (i.e. proportion of k neighboring cells with the same cell type annotation in a Nearest Neighborhood graph based on 500 highly variable genes and 50 principal components using function in package bluster^65^ version 1.14.0) and silhouette of cell type clusters (silhouette based on 500 highly variable genes and 50 principal components using function in package bluster version 1.14.0). For comparison of metrics, the metrics were weighted by number of cells per library for gene metrics, purity and silhouette and weighted by number of cells and UMI reads for CDR, respectively. To ensure comparability of the metrics, we restricted the representation of gene metrics distributions (Fig. 3G, Supp. Fig. 3F- G) to genes with nonzero values in both assays per cell type as well as to the 22,707 genes deemed to be expressed for the cellular detection rate, respectively.

### Summary statistics per cell type

To provide recommendations for researchers to choose a particular assay, we derived the following criteria of performance:

1. Gene Metrics

a. minimal standard error of mean gene expression (“Mean” in Figure 2G).
b. minimal variance of gene expression (“Variance” in Figure 2G).
c. minimal dropout of gene expression (“Dropout” in Figure 2G).
2. Cell Metrics

a. maximal cellular detection rate (“Detection” in Figure 2G).
b. maximal purity (“Purity” in Figure 2G).
c. maximal proportion of cells with a purity value > 0.5 (“Purity > 0.5” in Figure 2G).
d. maximal silhouette (“Silhouette” in Figure 2G)
e. maximal proportion of cells with a silhouette value > 0 (“Silhouette > 0” in Figure 2G).

In addition, we used the Kolmornov-Smirnov distance as a measure of reproducibility. In general, this metric quantified how (dis-)similar a pair of univariate distributions are, e.g., the mean gene expression between one snRNA-seq library compared to one scRNA-seq library, where a value close to zero indicated a high similarity. Given our experimental design where the expression was profiled in several patients using both scRNA-seq and snRNA-seq, we could compare the reproducibility of above-mentioned metrics per cell type within donors between scRNA-seq and snRNA-seq and across donors within one assay.

### Horizontal integration of sc/snRNA-seq and snATAC-seq modalities

Individual sn/scRNA protocols were preprocessed independently prior to integration, following consistent procedures as explained below. The Scanpy^66^ (v1.9.5) standard pipeline was employed to generate PCA-based latent spaces, using the top 50 Principal Components (PCs) as the latent representation. Sample batch correction within protocols was applied through the following methods:

- Harmony^19^ integration: Performed on the top 50 PCs using *run_harmony* function from *harmonypy* (v.0.4.7).
- Scanorama^67^ integration: Conducted using the *correct_scanpy* function (v1.7.3) according to the standard parameters as suggested in authors’ guidelines.
- scVI integration^23^: The scVI model (v.1.0) was configured with 256 nodes in the first hidden layer (*n_hidden*), two hidden layers in the encoder-decoder architecture (*n_layers*), and 30 nodes in the bottleneck layer (*n_latent*). For all latent spaces, the same set of 5,000 genes was used, identified using Scanpy’s ‘*cell_ranger’* algorithm.

For sn/scRNA integration between protocols, 5,000 highly variable genes were selected using ‘*seurat_v3*’, setting the protocol of origin as a variable to regress out. For tools supporting multiple batch-effect variables, samples of origin were also included beyond suspension type (nucleus/cell). This approach ensured consistency and effective integration across different sn/scRNA sequencing protocols.

For the snATAC-seq modality, the open chromatin regions data were represented in two ways for latent space computations:

1. Peaks (Variable-Length Windows): A peaks x nuclei matrix with a homogenous set of peaks across human samples was directly obtained from the *Cell Ranger ARC aggregate* pipeline as *filtered_feature_bc_matrix.h5*.
2. Bins (Fixed-Length Tiles): A bin feature matrix was created using the *add_tile_matrix* function from SnapATAC2^24^ (v0), based on the *atac_fragments.tsv* file provided by Cell Ranger ARC. The matrix consisted of fixed-size tiles for downstream dimensionality reduction tasks.

Depending on the requirements of the dimensionality reduction method, either the peak or bin matrix was selected.

- Latent Semantic Indexing (LSI): LSI was computed using Signac^25^ (v,1.14.0) with standard parameters. The first component of the LSI space was discarded, as typically associated with sequencing depth, and batch correction was performed using Harmony (via Seurat^20^ v5).
- scVI-Based Models: PeakVI^26^ and PoissonVI^27^ models from scVI (v.1.0) were configured with 512 nodes in the first hidden layer (n_hidden) and 30 nodes in the bottleneck layer (n_latent). Features were filtered to retain those present in at least 1% of cells. Training continued until the elbow loss converged (early_stopping=True). For additional analysis, the PoissonVI model was trained on a fragment matrix approximation derived from the peak matrix.
- Spectral Embedding: Based on Laplacian Eigenmaps, unimodal spectral dimensionality reduction was computed following SnapATAC2 guidelines, using a tile matrix with a 500-bin size and the top 200,000 features. Batch effect correction was evaluated using Mutual Nearest Neighbor^68^ (MNN) and Harmony with standard parameters.

### Benchmarking of computational tools for horizontal integrations in sn/scRNA and snATAC-seq data

For each protocol and modality integration described in the horizontal integration section, a benchmarking pipeline from scIB^12^ (v1.1.3) was executed for the computational tools tested. The benchmarking was performed using the two levels of annotations defined via label transfer. For individual sn/scRNA protocols and data harmonization across RNA protocols, all metrics included in the standard *scIB* Benchmarker function were applied. In the snATAC-seq scenario, the benchmarking pipeline was modified by removing the *Isolated_labels* metric from the *BioConservation* set and the *pcr_comparison* metric from the *BatchCorrection* set. These metrics were excluded as they were not informative and produced similar values across the different tools. Additionally, for the snATAC-seq horizontal integration, nuclei failing quality control in the snRNA modality were filtered out to include only high-quality nuclei from both modalities in the multiome experiment. Labels transferred from snRNA were propagated and used in the snATAC-seq data. For each protocol, the sample of origin was set as the batch variable. For integration across sn/scRNA-seq protocols, the batch variable was defined as the protocol of origin.

### Cell-type annotations across single-cell data types with scOMM

To achieve consistent and robust cell annotations across datasets and modalities, we developed scOMM (https://github.com/mereulab/scOMM/tree/master), a versatile, reference-based classifier designed for the automatic classification of single-cell multimodal data. Built within a unified architectural framework, scOMM features implementations in Keras, R, and Python, ensuring accessibility and reproducibility across diverse computational workflows. ScOMM employs a sequential neural network architecture with adjustable parameters, including the number of layer nodes, activation functions, dropout rates, and training settings, providing flexibility for various applications. The basic workflow of the scOMM annotation is the following:

#### Data Preparation

The *ds_prepare_data* function aligns the reference (*R*) and query (*Q*) datasets and extracts relevant features. Marker genes (*M*) are either user-specified or determined using the *FindAllMarkers* function (Seurat v.5.1.0). A gene *g* is retained if it satisfies:

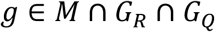

where *G*_*R*_ and *G*_*Q*_ represent the gene sets of the reference and query datasets, respectively.

The resulting expression matrices *X*_*R*_ ∈ ℝ^|*M*|×*nR*^ and *X*_*Q*_ ∈ ℝ^|*M*|×*nR*^, where *n*_*R*_ and *n*_*Q*_ are the number of cells in *R* and *Q*, are processed alongside their associated cell-type annotations (*C*_*R*_, *C*_*Q*_) for downstream tasks.

##### Data Splitting

The *ds_dplit_data_dnn* function divides the reference dataset *X*_*R*_ into training (*X*_*train*_) and test (*X*_*test*_) subsets while maintaining class balance. For each celltype *c* in *C*_*R*_:

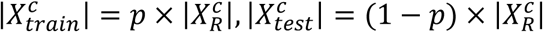

Where *p* is the proportion of data allocated to training, and 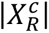 is the number of cells of type *c*.

Labels *C*_*train*_ and *C*_*test*_ are one-hot encoded into matrices *Y*_*train*_𝜖{0,1}^*ntrain*×*k*^ and *Y*_*test*_ 𝜖{0,1}^*ntest*×*k*^, where *k* denotes the number of cell types.

#### Model Architecture and Training

The deep neural network (DNN) model is constructed and trained using the *ds_dnn_model* function. Its architecture includes:

1. An input layer with |*M*| nodes.
2. *L* hidden layers, each with user-defined nodes (*h*_*l*_ for layer *l*), *ReLU* activation, and optional dropout. The transformation at layer *l* is expressed as:

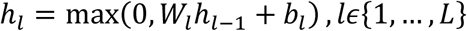 where *W*_*l*_ and *b*_*l*_ are the weight matrix and bias vector, respectively.
3. An output layer with *k* nodes and softmax activation:

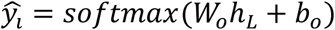

where 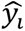 represents the predicted probability vector for cell *i*.

The model optimizes the categorical cross-entropy loss:

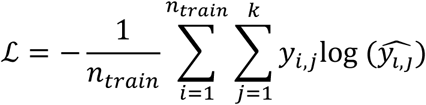

Where *y*_*i*,*j*_ and 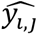 are the true and predicted probabilities for class *j* in cell *i*.

The Adam optimizer is used, with early stopping employed to prevent overfitting. Hyperparameters such as the learning rate (*η*), batch size (*b*), and dropout rate (*r*) are configurable.

#### Classification

The *ds_dnn_classify* function applies the trained model to the query dataset *X*_*Q*_, producing a probability matrix *P*_*Q*_ 𝜖[0,1]^*nQ*×*k*^:

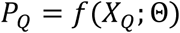

where *f* denotes the DNN and Θ its learned parameters. Classification is determined by:

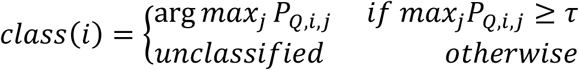

where *τ* is a user-defined threshold.

While originally developed for scRNA-seq data, scOMM is highly adaptable and extends its functionality to snATAC-seq data and transformed gene activity values, making it suitable for a wide range of omics datasets. This extension enables its application to cross-modality models, where the snRNA-seq data serve as reference for training the model to predict the query gene activity values inferred from snATAC-seq, and vice versa. A notable strength of scOMM is its bridge-like classification approach, which leverages the relationships between snRNA and snATAC profiles within the same individual nuclei. This approach enables the transfer of cell labels from chromatin accessibility profiles in multiomics datasets to unpaired ATAC data using the same basic workflow, replacing gene expression with chromatin accessibility features. By preserving consistency in cell annotations across both modalities and datasets, scOMM facilitates comprehensive, scalable, and integrated single-cell multiomics analysis.

### Label transfer from external references

For the cell-type annotation, we relied on scOMM. Feature selection for model training was based on cell-type markers, identified using the Wilcoxon test on log-normalized data. The model architecture shared several common elements, including *ReLU* as the activation function, batch normalization, weight regularization, and dropout. The initial learning rate for training was set to 0.001.

Two reference single-cell datasets from human kidney studies were used in order to guarantee precise cell type labeling and enable cross-referencing. One dataset contained snRNA-seq data, whereas the other contained both snRNA-seq and scRNA-seq data. One dataset captured finer cell states (Lake et al.), while the other provided broader annotations (Muto et al.).

#### 1. snRNA/snATAC-seq reference (Muto et al., 2021)

The reference dataset was obtained through the public data in Muto et al.^17^. Features were selected using the “*author_cell_type*” metadata via Scanpy^66^ software. For training, the top 400 non-overlapping gene markers per cell type were used. The model architecture consisted of three hidden layers with 256, 128, and 68 nodes, respectively, and a dropout probability of 0.5.

#### 2. Human Kidney Atlas (Lake et al., 2023)

The reference dataset was obtained through the public data in Lake et al.^5^, and was subsetted to include only cortex-associated states, either healthy or adaptive, from the reference samples.

- Level 1 Label Transfer (Broad Annotation): The top 500 non-overlapping features per cell type were selected as input for the model. The model architecture included two hidden layers with 1024 and 256 nodes, respectively, and training was performed with a dropout probability of 0.1. Predictions were made without applying an unclassified threshold, across all RNA-based protocols.
- Level 2 Label Transfer (Fine Annotation): Level 2 label transfer was conducted iteratively from Level 1, with one model trained per Level 1 cell type to predict the Level 2 sub-states. Each model used two hidden layers with 512 and 128 nodes, respectively. The top 300 non-overlapping features per cell type were selected as input. Training was performed with a dropout probability of 0.3.

### External references annotation matching

The Human Kidney Atlas (Lake et al., 2023) provides hierarchical annotations for kidney biology. From the publicly available dataset, we selected *subclass.l1* metadata as Level 1 (L1) annotation and *subclass.l3* metadata as Level 2 (L2) annotation. The kidney cortex was resolved into the following stratified categories: For PT (L1), this included PTS1/2, aPT, and PT-S3 (L2). TAL (L1) was annotated as C-TAL, aTAL1, MD, and aTAL2 (L2). PC (L1) comprised PC (L2). CNT (L1) included CNT and CNT-PC (L2). DCT (L1) was further resolved into DCT1 and DCT2 (L2). IC (L1) was divided into CCD-IC-A, IC-B, and CNT-IC- A (L2). PEC (L1) was annotated as PEC (L2). POD (L1) included POD (L2). EC (L1) was categorized into EC-PTC, EC-GC, EC-AEA, and EC-LYM (L2). FIB (L1) comprised FIB, MYOF, and aFIB (L2). VSM/P (L1) included VSMC/P, VSMC, MC, and REN (L2). Finally, IMM (L1) encompassed T, MDC, NKC/T, ncMON, MAC-M2, B, N, PL, cDC, pDC, and MAST (L2).

The snRNA/snATAC-seq reference (Muto et al., 2021) provides a single annotation layer, defined in the publicly available dataset as *author_cell_type* metadata. This annotation broadly matches the Level 1 annotation from the Human Kidney Atlas, with increased resolution for certain populations. Here, we provide the annotation match between these two external datasets: CNT (L1), DCT (L1), ENDO (L1, corresponding to EC), FIB (L1), ICA (L2, corresponding to CCD-IC-A under IC), ICB (L2, corresponding to IC-B under IC), LEUK (L1, corresponding to IMM), MES (L1, corresponding to VSM/P), PC (L1), PEC (L1), PODO (L1, corresponding to POD), PT (L1), TAL (L1), and PT_VCAM1 (L2, corresponding to aPT under PT).

### Cell identity harmonization following integration based on clustering

To harmonize cell identities after data integration, we performed clustering followed by manual annotation using known cell type-specific markers. Clustering was performed to define two main annotation levels, L1 and L2, as outlined in the supervised label transfer approach with scOMM, which was applied independently for each assay to minimize biases. Leiden^69^ clustering was conducted on a k-nearest neighbors (k=50) graph, using a range of resolution values (0.3, 0.6, 1, 1.4, 2, 6, 10). Final cluster labeling was determined by majority voting based on labels transferred at each annotation level, with resolution level 6 used as the primary reference. Results were then visually inspected for cluster consistency, and marker-based supervision was applied for accurate cell type annotation. If needed, cluster assignments were refined by adjusting the resolution to balance cluster structure and cell identity.

### Population reclustering

Reclustering is performed with Leiden^69^ algorithm on a k-nearest neighbors (k=10) graph derived from the embedding computed on the whole dataset. No recomputation of highly variable genes or embedding was performed.

### Contribution of sn/scRNA-seq protocols to cell type identification

To evaluate the contribution of each sn/scRNA-seq protocol to cell type annotations, partial area under the receiver operating characteristic curve (pAUC) by specificity was employed using the R package pROC^70^ (v.1.18.5), focusing on the high-specificity range (values > 0.9) to minimize false-positive assignments. Partial AUCs were calculated separately for Level 1 (broad annotation) and Level 2 (fine annotation) label transfers. The curated manual clustering annotation, derived after sn/scRNA-seq horizontal integration, was used as the ground truth for cell-type annotation.

### Vertical integration of snRNA- and snATAC-seq data

Integration of paired snRNA and snATAC modalities from the multiome dataset uses nuclei (observations) as anchors to generate a joint embedding across modalities. A joint embedding can be generated either directly from the raw feature spaces of both data types (snRNA-seq and snATAC-seq) or by using pre-computed embeddings specific to each modality. Preprocessing settings and highly variable feature selections are imported and fixed from horizontal unimodal integration. Final vertical embedding representation was performed with the SnapATAC2 module through Laplacian Eigenmap computation on both modalities simultaneously (i.e., *snapatac2.tl.multi_spectral*), with linear space and time complexity, to obtain the joint latent space. MNN was applied to mitigate the batch effect. UMAP projection was computed on a 50 kNN graph.

### Mosaic integration

Integration of paired and unpaired sn/scRNA-seq and snATAC-seq datasets was achieved using scVI tool (v.1.0) through the MultiVI^18^ model. This model was trained on the previously selected highly variable features, with protocol and sample origin included as categorical covariates for batch effect correction. Clustering of the joint embedding was computed on a k-nearest neighbors (k=50) graph using the Leiden^69^ algorithm. Differential expression analysis (Wilcoxon test) on the sn/scRNA-seq data was used to detect and exclude low-quality cells and outlier populations inconsistent with the kidney cortex biology. The final UMAP projection was performed on a 50 kNN graph.

### Diagonal integration

To explore global integration potentials, we simulated fully unpaired datasets by decoupling snMultiome modalities, thereby testing diagonal integration. For this purpose, we utilized GLUE^29^ as a representative method for state-of-the-art unpaired multimodal integration. Preprocessing and feature selection from both sn/scRNA- and snATAC-seq horizontal integrations. The GLUE model was trained according to the authors’ guidelines. During dataset configuration (via *scglue.models.configure_dataset*), cells from each modality were assigned a batch effect variable defined as a combinatorial factor of sample origin and protocol. The best-performing embedding from horizontal integration task was used as the guidance embedding for each modality.

### Systematic evaluation of vertical integration embedding

To evaluate vertical integration embedding, we employed the Weighted Nearest Neighbor (WNN) algorithm from Seurat as a baseline. WNN leverages unimodal latent spaces to assign weights to each cell and modality, generating a final multimodal latent space. We computed WNN embeddings twice: (1) using standard unimodal embeddings as per the authors’ vignette (Standard WNN), and (2) using the best-performing embedding from each modality obtained during horizontal integration (referred as to optimized WNN).

The performance of the optimized WNN embedding was systematically compared to that of the multi-spectral MNN-corrected embedding (described in Vertical Integration of snRNA-seq and snATAC-seq Data) across every L2 population. Silhouette width and b-cLISI scores were computed for each cell in unimodal and multimodal embeddings using scOMM labels as the cell-type reference. The distributions of cell-wise Silhouette values for multimodal embeddings were compared via scatter plots (generated with Seaborn^71^ v0.13.0). Average deviations from the diagonal (referred to as Avg. Dev.) toward a specific axis was used to quantify the performance gap between embeddings in detecting and isolating cell types or states. It was computed by averaging pairwise distances between the two distributions. To provide a broader comparison perspective, a heatmap was generated showing the average metric values per L2 population for multiome cells with high-quality data in both RNA and ATAC modalities.

### Systematic evaluation of diagonal and mosaic integration embeddings

To systematically evaluate the ability of unimodal and multimodal approaches to detect and isolate cell types and states, silhouette width and b-cLISI scores were computed using scOMM labels. For each population category, cell-wise metric values were averaged and displayed in a heatmap, with integration methods as rows and populations as columns.

### Systematic evaluation of single-cell data integrations using scOMM for supervised label transfer

ScOMM was employed as a benchmark tool to systematically evaluate the integration of the distinct data types, focusing on how different strategies preserved biological information, maintained data quality, and achieved consistent cell-type annotations. Horizontal and vertical integration were assessed by quantifying modality-specific contributions and performance using metrics such as cell-type identification accuracy and marker detection efficiency.

### Cell type predictability of sn/scRNA-seq assays before and following horizontal integration

To quantify cell type predictability (L1 granularity) for each individual sn/scRNA-seq protocol, we trained three distinct scOMM models, one for each protocol. For model training, ∼6,000 nuclei/cells per dataset were used, representing the maximum number that could be equally included across all three protocols. Additionally, a combined model was constructed by randomly sampling ∼2,000 nuclei or cells from the training data of each protocol, resulting in a combined dataset of ∼6,000 nuclei/cells. The remaining cells were allocated as testing data. Each model was subsequently used to predict cell types in the other datasets. Model performance was evaluated using the total area under the curve (AUC), with AUC scores computed using the *bench_calcAUC* function in scOMM. Predictions from the *ds_dnn_classify* scOMM function were compared to the original protocol-specific cell annotations. The visualization of this comparison was generated via the *bench_plotTileAUC* function in scOMM.

### Comparative analysis of snRNA- and snATAC-seq data modalities in cell type identification (label transfer-based)

To compare snRNA- and snATAC-seq data in cell type identification under a supervised, label transfer-based scenario, three scOMM models were evaluated using cell type annotations from snRNA-seq data as the ground truth:

1. Cross-Modality model: trained on snRNA-seq data as the reference and tested on gene activity scores derived from snATAC-seq.
2. Bridge model: both training and testing were conducted within the snATAC-seq modality, using peak data.
3. Integrated model: combined features from the training data of both previous models to assess whether integrating RNA and chromatin features improves cell type resolution.

To ensure comparable cell composition across models, 70% of nuclei (∼26,400 cells) were sampled as the training set for the gene-based, peak-based, and integrated models, with the remaining 30% used for testing. Sampling was performed using the *ds_split_data_dnn* function, and training was conducted using the *ds_dnn_model* function, both implemented in scOMM. Performance evaluation focused on two key metrics:

i) Cell type prediction accuracy: to assess the relative contributions of RNA and ATAC modalities in resolving cell types, AUC scores were computed using the *bench_calcAUC* function. Predictions from the *ds_dnn_classify* function were compared against the ground truth.
ii) Identification of cell type-specific features: significant markers were identified through feature importance scores and marker detection rates, evaluated using the *ds_feature_importance* and *bench_MarkerDetect* functions, respectively, in scOMM.

### Feature importance in scOMM

Feature importance (FI) scores were calculated from the scOMM models using a permutation importance algorithm as implemented in the *ds_feature_importance* function. This approach evaluates the relative contribution of individual features, such as genes, peaks, or their combination, to the model’s overall cell type prediction performance. The algorithm works by systematically setting the values of a single feature to zero across all cells and measuring the resulting impact on model accuracy. Features that cause a drop of more than 10% in accuracy are assigned the highest importance scores.

### Marker detection rates

The marker detection rate comparison is conducted using the function *bench_MarkerDetect* (in scOMM), where each marker detection rate was calculated based on the feature importance scores generated by each model. For each cell type, marker detection rate was defined as the proportion of significant features (importance scores ≥ 10) relative to the predefined reference list of markers derived from the Human Kidney Atlas. This allowed the quantification of each model’s capacity to detect biologically meaningful markers across sn/scRNA-seq, snATAC-seq, and their integrated datasets. For models using genomic sites (peak-based and peak-integrated models), the *ClosestFeature* function from Signac^25^ (v1.14) was employed to map each genomic site to its nearest gene.

### Embedding-based metrics for systematic evaluation of data integration

The primary metrics employed in this evaluation include the Mahalanobis distance and width silhouette, which serve as global measures of cell type separability, and the binary version cell type Local Inverse Simpson’s Index (b-cLISI), which provides a local assessment of cell-type preservation and accuracy. For all these metrics, calculations were performed at the level of cell types, where cell types were inferred using scOMM.

### Multidimensional Mahalanobis Distance

This metric calculates the distance between two distributions, one as a reference (*V*_*i*_) and the other as a query (*V*_*j*_), in an N-dimensional embedding space. The distance (*D*_*ij*_) is computed as the separation from the reference distribution *V*_*i*_ to the centroid of *V*_*j*_, measured in units of dispersion of *V*_*i*_. The formula is as follows:

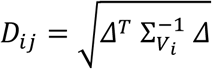

where:

- *Δ* = *μ*_*V_i_*_ − *μ*_*V_j_*_
- 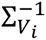 is the inverse of the covariance matrix of *V*_*i*_, computed using the Ledoit-Wolf shrinkage adjustment (Scikit-learn v1.4.0)
- 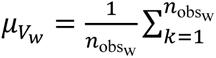 *V*_*wk*_ is the centroid of *V*_*w*_

Each cell type defines a distribution in the N-dimensional embedding space, allowing for Mahalanobis distances to be calculated between cell types. Distances greater than 3 indicate good separation in low-dimensional embeddings. For larger embeddings or precise evaluations, the chi-squared distribution can serve as a guide (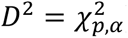), where *p* is the dimensionality of the embedding and α is the overlap ratio). As this measure is not symmetric (*D*_*ij*_ ≠ *D*_*ji*_), the harmonic mean is used to summarize distances between two populations.

### Accuracy of cell-type identification in the embedding-based (unsupervised) scenario

Cell type identification performance was assessed using the following embedding-based metrics. The evaluation combined the median values of b-cLISI and Silhouette Width (SW) scores for each cell type, averaging these values across all observations in the category to produce a final score. The SW metric evaluates label identity assignments within N-dimensional embeddings. It calculates the ratio between intra-category distances and inter-category distances (to the nearest distinct category) for individual observations. Implemented in the scIB^12^ package, the *silhouette_samples* function was used to compute sample-wise scores, providing insights into how well clusters are separated in the embedding space.

### Binary Cell-Type Local Inverse Simpson’s Index (b-cLISI)

The Binary Cell-Type Local Inverse Simpson’s Index (b-cLISI) score is a variant of the cLISI to assess local neighbor diversity in a neighborhood graph, considering only the label of the central node’s category. This modification enhances interpretability by focusing on intra-category consistency. For a node *i* with neighborhood *N*(*i*) and category *j*, the formula is:

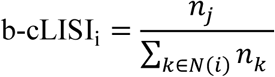

where *n*_*j*_ is the number of nodes in *N*(*i*) belonging to category *j*. The b-cLISI metric ranges between 0 and 1, where higher scores indicate better local clustering and consistency for the same cell type.

## Differential expression and accessibility analyses

Differential analyses were conducted for each individual protocol and modality using annotation labels derived from the label transfer tasks at the two levels of granularity. For sn/scRNA-seq protocols, differential expression analysis was performed using Scanpy’s^66^ *rank_genes_groups* function with the Wilcoxon method on log-normalized data. For the snATAC-seq modality, cell type-specific narrow peak calling was conducted using the *macs3* function in SnapATAC2 with standard settings. Differentially accessible regions (DARs) were identified on a cell type basis from peak calling results using the *diff_test* function in SnapATAC2, with thresholds set to *min_lfc=0* and *min_pct=0.01*. Accessible regions were mapped to “accessible genes” by extending 2000 bp upstream and 500 bp downstream from the transcriptional start site, based on the GRCh38 genome reference.

Differential accessibility results were propagated to genes in the following way:

1. P-values for accessible regions were aggregated to corresponding genes using Stouffer’s Z-score method. This was implemented via custom Python scripts utilizing the *ppf* and *cdf* functions from the *scipy.stats.norm* module (v1.11.4).
2. Log Fold change is approximated by averaging the linear fold changes of individual features assuming all features contribute equally to the aggregate:

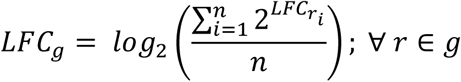

A significance score (*w*_*r*_(*g*)) was calculated as:

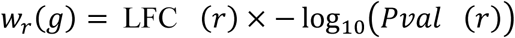

These scores were averaged across regions associated with the same gene. Statistically significant expressed genes were defined using thresholds of a minimum log fold change of 0.25 and an FDR-corrected p-value of 0.05. For accessible genes, significance scores were thresholded at the 0.8 quantile, enabling comparisons with statistically significant differentially expressed genes across protocols.

### Identification of cell type-specific markers

We implemented a postprocessing step on differential expression results to identify markers that are specific to individual populations within a given batch (i.e., protocol or modality). These markers were further validated to ensure they were not statistically significant in any other population within the same batch. Filtering was performed using thresholds for adjusted p-value, log-fold change, and mean expression level, which could be customized to meet the user’s desired level of statistical rigor.

For each batch:

1. Candidate markers for each population were compared against genes identified in other populations within the same batch, excluding overlapping or non-significant genes.
2. Markers unique to each population were identified, ranked by significance scores, and compiled into a sorted list for each population.

This approach enabled the precise identification of population-specific markers, facilitating robust and reliable downstream analyses. This was used for the discovery of consensus sets of markers across protocols.

### Norn signature scoring

A Norn signature score was calculated in each nucleus/cell using a reference set of marker genes (DCN, PDGFRA, GSN, TIMP1, CFD) by using the *score_genes* function from Scanpy^66^ with default parameters. This method assigns a score to each cell based on the average expression of the provided marker genes subtracted with the average of a reference set of genes drawn randomly from all expressed genes. This score enables the identification and quantification of cell-type-specific activity across the single-cell RNA-seq dataset.

### Downsampling for the comparison of assay-specific cell type markers

Custom downsampling workflow was developed for sn/scRNA-seq data to ensure proportional representation of cell types and preservation data distributions while standardizing number cells and counts depth across dataset, seeking to avoid technical biases on the comparison of marker features. Data from each protocol were first downsized to 7,500 cells. Its implementation on Python was achieved through *StratifiedShuffleSplit* function from sklearn^72^ (v1.4.1) model_selection module, providing the cell type L1 labels to maintain the population proportion on the downsized object. Checks were done for the maintenance of total counts and number of features per cell distribution quantiles within their original values. Once objects were downsized, *downsample_counts* function from Scanpy^66^ preprocessing module was employed, setting an upper threshold of 10,000 total counts per cell.

### Benchmarking summary scores

We focused on two primary analytical tasks: cell type identification and marker detection, each implemented through label transfer–based (supervised) and embedding-based approaches (unsupervised). To facilitate a concise comparison of results, a single metric was computed derived from the analyses within each analytical arm, providing an integrated perspective on the overall performance of the different methodologies and cell types.

- Cell-type identification in label transfer-based scenario: population wise AUC predictions for each scenario scOMM model were averaged across testing sets to obtain the final population score.
- Cell-type identification in embedding-based scenario: cell-wise width silhouette and b- cLISI scores were averaged per cell group, then their values were averaged across populations to obtain the average score.
- Marker detection in label transfer-based scenario: final score was drawn proportion of reference marker sets provided in Lake et al.^5^ detected as significant features for each scenario scOMM model based on feature importance score.
- Marker detection in embedding-based scenario: marker detection scores were based on differential expression analysis statistical results from each individual dataset/modality.

### Marker detection metric

A composite marker detection score was used to evaluate the uniqueness and statistical significance of marker features across cell types and protocols. For a given technology (*t*) and cell type (*c*), a set of marker genes were computed (*G*_*t*,*c*_ = {*g*_1_, *g*_2_, …, *g*_*n*_ }), with their associated statistics (*LFC*_*tc*_(*g*), *Pval*_*t*,*c*_(*g*)), the following scores were computed:

1. Uniqueness score (*U*_*tc*_): Is computed as the summatory over the *LFC*_*tc*_ of markers (*G*_*tc*_′) for *c*, found only in *t* and not in any other technology (*t*’). In the case that a gene is considered marker (*G*_*tc*_′′) for any other cell-type (*c*’) in any other technology, a penalty (*P*_*t*,*c*_(*g*)) is applied by subtracting *LFC*_*t*′*c*′_.

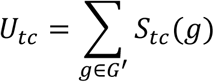

Where:

- 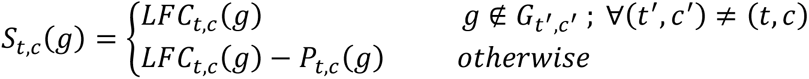
- 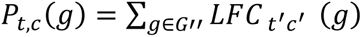
- 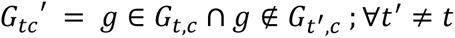
- 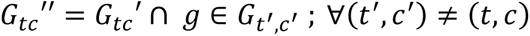

2. Power score (*W*_*tc*_): For each gene, a significance score (*w*_*tc*_(*g*)) was computed based on (*LFC*_*tc*_(*g*), *Pval*_*t*,*c*_ (*g*)). Power score is computed as the summatory of the significance score over markers (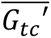) for c, found in at least more than one technology.

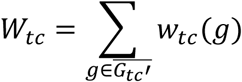

Where:

- 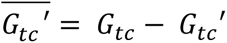
- 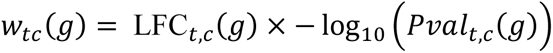

The final marker detection score was a weighted sum of the uniqueness and power scores, with adjustments for snATAC-associated features on the power score computation (weighted by 0.3 to account for differences in feature statistics between snRNA and snATAC data).

### Defining Cell-Type-Specific CREs Using Accessibility and Transcription

CREs were defined using scATAC-seq as described. Accessibility at CREs in single cells was quantified as described. Transcription within CREs in individual cells was quantified using SCAFE^73^ v1.0. Briefly, the 5’ scRNA-seq read alignments were converted to capped TSS (CTSS) BED files and the number of UMIs within each CRE was counted in a strand-agnostic manner. CTSS signals from all 5’ scRNA-seq samples were clustered into TSS clusters using default SCAFE parameters. These clusters were subsequently used to identify transcribed CREs. Cell types in scATAC-seq nuclei and 5’ scRNA-seq cells were annotated as described. The specificity of CREs in each level 2 cell type were calculated using Wilcoxon-rank sum tests implemented in the *scanpy.tl.rank_genes_groups* function in Scanpy^66^, based on either accessibility or transcription. Three sets of cell-type-specific CREs were defined, including Set 1: CREs ranked by accessibility, Set 2: Transcribed CREs ranked by accessibility, and Set 3: Transcribed CREs ranked by transcription. CREs were ranked by Wilcoxon p-values (log_2_ fold change > 0) and the top 3000 CREs were defined as cell-type-specific. Cell types with fewer than 3,000 CREs in each set were discarded. Set 1, 2 and 3 contains 34, 27 and 13 cell-types, respectively.

### Enrichment of Trait Heritability in Cell-Type-Specific CREs

Enrichments of heritability of two traits, i.e., hypertension^42^ and estimated glomerus filtration rate (eGFR)^43^, were assessed in the three sets of cell type specific CREs using partitioning linkage disequilibrium score regression implemented in LDSC^74^ v1.0.1. GWAS summary statistics for hypertension were obtained from UK Biobank^75^ (phenotype code 6150), and those for eGFR were obtained from the GWAS catalog (study GCST90428446). The summary statistics of eGFR were munged using the “munge_sumstats.py” script and that of hypertension were used as it is. Annotation files and LD score files for each set of CREs were generated using the “make_annot.py” and “ldsc.py” scripts with default parameters. Each set of CREs was added to the 97 annotations of the baseline-LD model v2.2, and heritability enrichment (i.e., the ratio of the proportion of heritability to the proportion of SNPs) for each trait in each set of CREs was estimated using the “ldsc.py” script with the “--h2” flag under default parameters. All cell types from Set 1 were analyzed in Fig. 5F and overlapping cell types across all sets were analyzed in Fig. 5G.

## Supplementary material

### Comparison of gene regulatory network inference across single-cell platforms

Understanding cell-type-specific gene regulatory networks (GRNs) is essential for unraveling the mechanisms that drive cellular identity and function^76^. Single-cell sequencing technologies, such as 3’ and 5’ scRNA-seq as well as joint snRNA-seq and snATAC-seq data, provide distinct insights into transcriptional regulation^77,78^. However, the robustness, reproducibility, and biological relevance of GRNs inferred from these platforms are not well understood. To address this, we developed a systematic framework to compare GRNs across platforms based on predictive accuracy, reproducibility of transcription factor (TF) identification, cross-platform complementarity, and biological validation with independent datasets.

We analyzed data from three donors (“lib_09,” “lib_10,” and “lib_36”) across all three platforms, encompassing 25,669 nuclei/cells (4,393 from 3’ scRNA, 6,416 from 5’ scRNA, and 14,860 from multiome). After filtering for features expressed in at least 30 cells, gene expression was log(1+x)-scaled, and peak accessibility was binarized. The top 1,000 most highly variable genes were selected per platform using Scanpy’s^66^ *sc.pp.highly_variable_genes* with the “batch_key” set to the biological sample identifier.

### Regulatory network inference

We inferred gene regulatory networks within each sequencing platform and biological sample independently. This entailed fitting a gradient-boosting-machine (GBM) tree regression model to each gene to predict its expression, similarly to the approach used by SCENIC^77^. We used the LightGBM^79^ package, with an ensemble of 20 estimators per model, learning rate of 0.5 and early stopping. For each gene, we used as input features: (1) gene expression data for all transcription factors (selected using JASPAR 2024^80^); and for only the multiome data, (2) chromatin accessibility of peaks proximal to the gene (within +/- 5kb), using the PyRanges^81^ package and TSS loci from RefTSS^82^ v4.1). This approach allowed us to treat “snRNA” as an additional modality, which is simply the multiome data excluding the chromatin accessibility features. To identify regulators for each gene, we assigned TFs based on their importance scores (total information gains from tree splits using that TF). A TF was considered a regulator if its importance score for a given gene exceeded the 95th percentile of scores globally across all genes within the sample and platform. Since the models were trained on cells pooled from all cell types within each sample, the resulting GRNs did not inherently include “cell type specificity” for network edges. To address this, we assigned cell type specificity to TF-target links that passed the importance filtering by evaluating the Pearson correlation between the TF and target gene’s ground-truth expressions in the training set. A link was deemed active in a particular cell type if the Pearson correlation exceeded 0.2.

### Cross-platform prediction accuracy

To assess the cross-sample generalization ability of models trained on each platform, we evaluated each model trained for that platform on the remaining held-out biological replicates. Specifically, for each pairing of training and held-out samples, we computed the Pearson correlation between the predicted and true log-transformed expression values for each gene (across cells), stratified by cell type (Supp. Fig. 14A). The 5’ scRNA-seq model demonstrated slightly higher but statistically significant predictive accuracy compared to 3’ scRNA-seq and multiome-derived models (Supp. Fig. 14B, left). However, the overall distributions of correlations revealed considerable variability across cell types, suggesting that biological and technical heterogeneity and cell-type-specific differences within patients may influence model performance for the gene expression inference through GRN (Supp. Fig. 14B, right).

### Reproducibility of TF identification

To estimate robustness of TF identifications within platforms, we calculated, per cell type, two metrics: (1) mean number of TFs identified per cell type across replicates; (2) consensus TF count: defined as TFs identified consistently in all replicates within each cell type. We defined a “reproducibility ratio” for each (platform, cell type) combination, as the latter quantity divided by the former. Overall, snMultiome exhibited the highest reproducibility (Supp. Fig. 14C, right), underscoring its robustness in detecting cell-type-specific regulators. While 5’ scRNA-seq also performed well in certain cell types, it exhibited greater variability compared to snMultiome. By contrast, 3’ scRNA-seq demonstrated the lowest reproducibility ratios, suggesting that RNA-only approaches may be less reliable for consistent TF inference.

### Cross-Platform complementarity in TF-target associations

To evaluate potential complementarity of cell-type-specific transcriptional regulators detected across sequencing platforms, we represented each consensus TF-target regulatory network as a binary indicator vector, where each entry denoted the presence or absence of a TF-target link. Pairwise Jaccard similarity scores were then computed between these vectors for each (platform, cell-type) pair, representing the intersection-over-union of shared regulatory elements across platforms. Hierarchical clustering revealed that platform-specific effects were more prominent than biological differences, with GRNs clustering primarily by platform rather than cell type (Supp. Fig. 14D). To characterize cross-platform overlap at the aggregated (cell type, TF) level, we also calculated the number of (cell type, TF) identifications shared by consensus networks between each pair of platforms. As expected, multiome and snRNA exhibited significant overlap in TFs due to their common RNA-sequencing foundation and potential representation of the same cells (Supp. Fig. 14E). However, their overlap with 3’ and 5’ scRNA, was considerably lower (Supp. Fig. 14E). These differences appear to be influenced by the RNA sequencing protocols rather than the inclusion of chromatin accessibility data, as multiome and snRNA exhibit much higher log-odds ratios (LORs) compared to the others scRNA-seq platforms (Supp. Fig. 14E, right). This indicates that complementary regulatory elements can be identified by scRNA-seq, with 5’ scRNA-seq detecting the highest number of unique TF across cell types (Supp. Fig. 14E, left). Altogether, this underscores the role of platform-specific technical factors in shaping GRN architecture.

### Validation using independent datasets

A TF needs to localize to the nucleus to act as a regulator. To test whether regulatory inferences from each platform satisfy this property, we cross-referenced consensus TFs identified per platform against localization data from v23 of the Human Protein Atlas^83^ (HPA). Specifically, we queried the HPA for genes with immunohistochemical staining patterns annotated as “detected” in the kidney, with a reliability score of “Enhanced”, and intersected these against TFs listed in JASPAR 2024. Intersecting the inferred TFs from each consensus network yielded a set of TFs per platform with independent evidence of nuclear localization in kidney tissue. With this approach, we found that a subset of inferred TFs was supported by experimental evidence (Supp. Fig. 14F). While the proportion of nuclear-localized TFs was consistent across platforms, 3’ scRNA-seq exhibited the highest validation fraction.

We also validated whether predicted regulatory TF-target associations were corroborated by independent evidence of TF binding or chromatin accessibility. To do so, we cross-referenced our consensus networks with bulk epigenomic data in kidney tissue. Using ChIPAtlas 2021^84^, we retrieved bulk ChIP-Seq peaks for transcription factors in all kidney cell lines, and intersected those with bulk ATAC-Seq peaks from kidney cortex (All peaks were filtered at reported significance level >= 50). We then linked each peak to the closest promoter in RefTSS v4.1 within a ±500 bp window. This yielded a set of potential TF-gene associations based on promoter binding in bulk samples. Altogether, this validation of TF-target associations using bulk ChIP-Seq data revealed low validation rates across all platforms (Supp. Fig. 14G). However, while the 3’ scRNA-seq platform seems to recover more consensus edges in ChIP-Seq promoters, the 5’ scRNA-seq platform, along with multiome, demonstrated the highest total average TF detection, suggesting that 5’ scRNA-seq and multiome may capture a broader repertoire of transcription factors, potentially enhancing the scope of inferred regulatory networks despite lower promoter-level validation in bulk ChIP-Seq data.

## Supplementary Figures

**Supplementary Figure 1.**
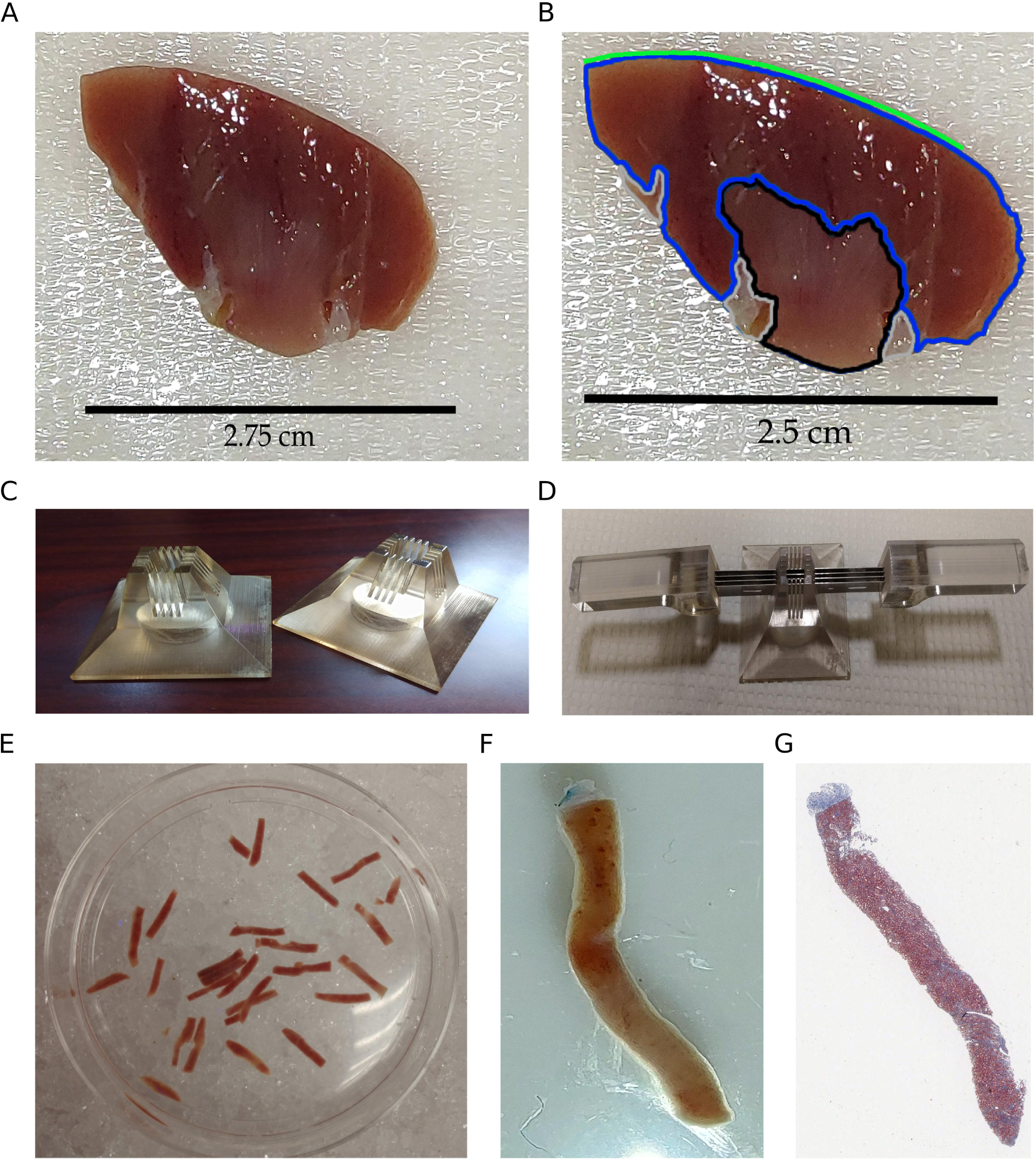
A) Live normal kidney tissue is obtained directly from the operating room at Michigan Medicine. Ideal procurements are a minimum of 1.5 cm x 1.0 cm x 1.0 cm in dimension and represent both renal cortex and medulla. B) Cortex (blue), medulla (black), capsule (green), corticomedullary junction (blue-black), and large arteries (white) are annotated. Tissue sections are chosen which contain complete transitions from renal capsule to corticomedullary junction. C) Using a 3D printed device (PRECISE Pyramid), nephrectomy specimens are cut into mock 16 gauge biopsy cores. D) The Precise Pyramid creates 25 identically proportioned mock 16 gauge biopsy cores per procurement. E) Biopsy cores are preserved across a variety of media, including but not limited to FFPE, CryoStor, RNALater, OCT, and liquid nitrogen (LN_2_). F) Biopsy core preserved in hypothermosol/CryoStor10. Note capsule, cortex, corticomedullary junction, and medulla. G) FFPE preserved core, stained with Masson Trichrome and digitally scanned at 40x from the same case.

**Supplementary Figure 2.**
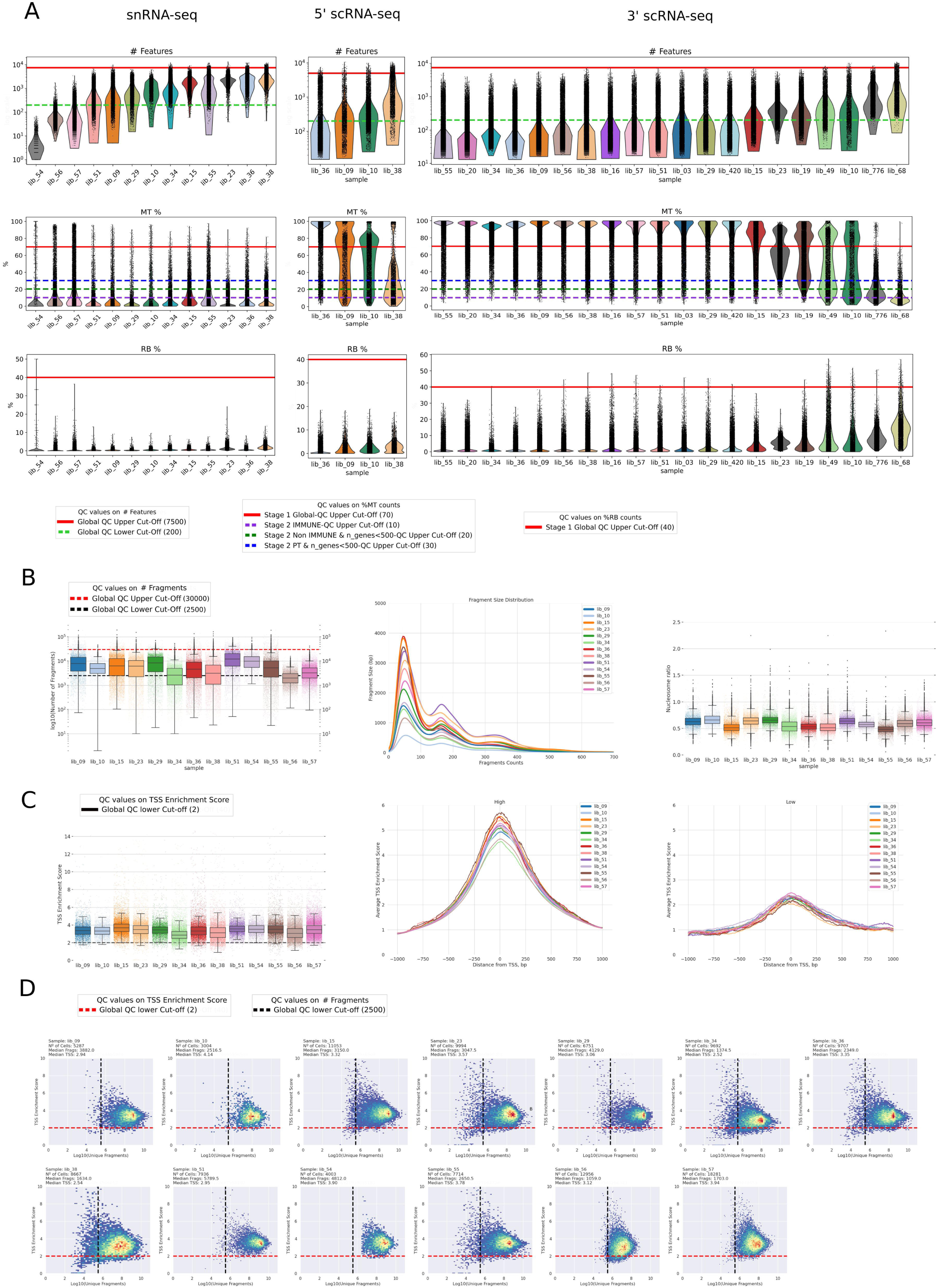
A) Violin plots showing quality control metrics for RNA-based single-cell technologies: number of unique features (genes) detected per cell (top row), percentage of mitochondrial reads (MT %) (middle row), and percentage of ribosomal protein genes (RB %) (bottom row). Metrics are displayed for snRNA from the Multiome assay (left column), 5’ scRNA-seq (middle column), and 3’ scRNA-seq (right column). Upper and lower thresholds for quality filtering are indicated by solid and dashed lines, respectively. B) Quality control metrics for snATAC-seq related to fragment count and length. Boxplot (box center represents the median, box size represents the interquartile range (IQR), and whiskers extend the smallest and largest value within 1.5 times the IQR) of the number of fragments per cell for each sample (left), fragment size distribution per sample (middle), and boxplot of nucleosome signal ratios per cell for each sample (right). C) Quality control metrics for scATAC-seq related to transcription start site (TSS) enrichment. Boxplot of TSS enrichment score distributions per sample (left), average TSS enrichment score distribution centered around the TSS site for cells with high TSS enrichment (middle), and average TSS enrichment score distribution across TSS sites for all high-enrichment cells (right). D) Bivariate histogram of TSS enrichment scores and the number of unique fragments per cell, illustrating the relationship between these two quality control parameters.

**Supplementary Figure 3.**
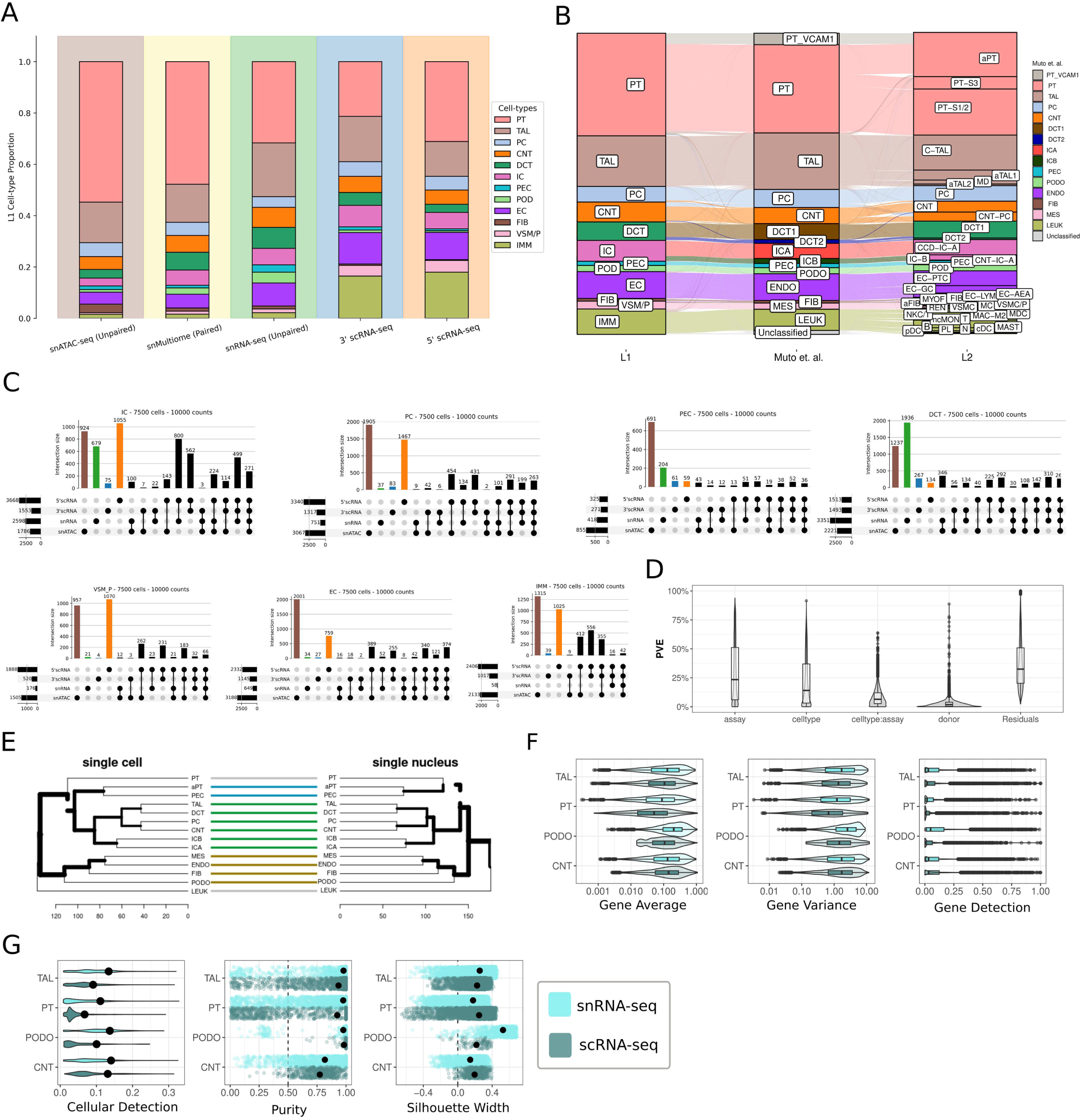
A) Stacked barplot of L1 cell-type composition across single cell data modalities. B) Alluvial plots displaying the distribution of label transfer annotation from external references Lake et al.^5^ - L1 and Muto et al.^17^ C) Upset plot of statistically significant genes on downsampled data across mBDRC protocols for Parietal Epithelial cells (PEC), Intercalated cells (IC), Distal Connecting Tubule cells (DCT), and Principal cells (PC), Vascular Smooth Muscle/Pericytes (VSM-P), Endothelial cells (EC) and Immune cells (IMM). D) The partial variance explained (PVE) by assay, cell type and donor using pseudobulk samples of 14 cell types with matched 3’ single-cell and single-nucleus libraries of 5 donors. The distributions of PVEs for the 22,707 genes are plotted as violin plots with median (bar), IQR (box), and outliers (dots). E) Tanglegram of Cell Type Dendrograms. Dendrograms are derived from hierarchical clustering using Euclidean distances and complete agglomeration of cell types per assay whereas thicker lines represent more confident clustering. The connecting lines between the dendrograms represent the local optimal solution of similarity based on a stepwise greedy forward selection algorithm (entanglement = 0.01), resulting in a nearly congruent tree structure. Some lines are not connected due to differences in clustering structures and the algorithm not forcing matches when similarity is insufficient. Dashed vs. full thick lines indicate slight differences in internal branch node splitting from the greedy algorithm, but the branch tips maintain the same similarity order. F) Mean gene expression, variance of mean expression and detection rate weighted by number of cells per library for single cell and single nuclei assays. The distributions of all genes with nonzero values in both assays (average and variance) as well as all 22,707 genes detected in either assay (detection) are plotted as violin plots with median (bar), interquartile range (box) and outliers (dots). G) Cellular detection rate per assay weighted by number of reads and cells per library for single cell and single nuclei assay, purity and silhouette index per assay. The dot represents the weighted median per assay.

**Supplementary Figure 4.**
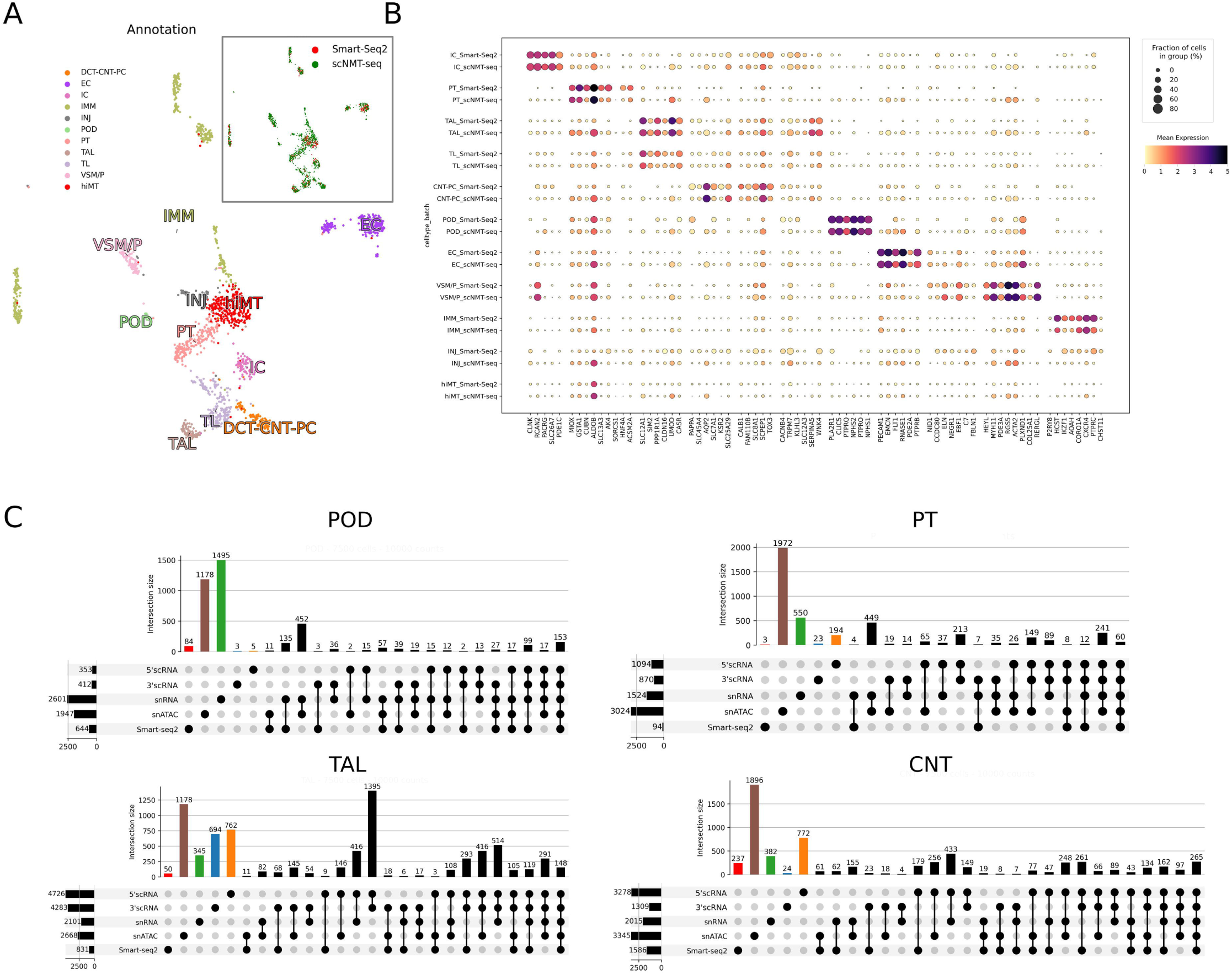
A) UMAP embedding of 1,842 cells from the Smart-seq2 and scNMT-seq transcriptomics dataset, showing L1 cell type populations, high mitochondrial (hiMT) cells, and injured state (INJ) cells. The top-right inset displays the same UMAP with colors indicating the scRNA-seq technology used for the experiment. B) Average expression values across Smart-seq2 and scNMT-seq cell populations in the mBDRC consensus marker analysis, consistent with Fig. 2D. C) Upset plots displaying statistically significant genes identified in downsampled data across mBDRC protocols, including Smart-seq2/scNMT-seq data, for podocytes (POD), proximal tubule (PT), thick ascending limb (TAL), and connecting tubule (CNT) populations markers.

**Supplementary Figure 5.**
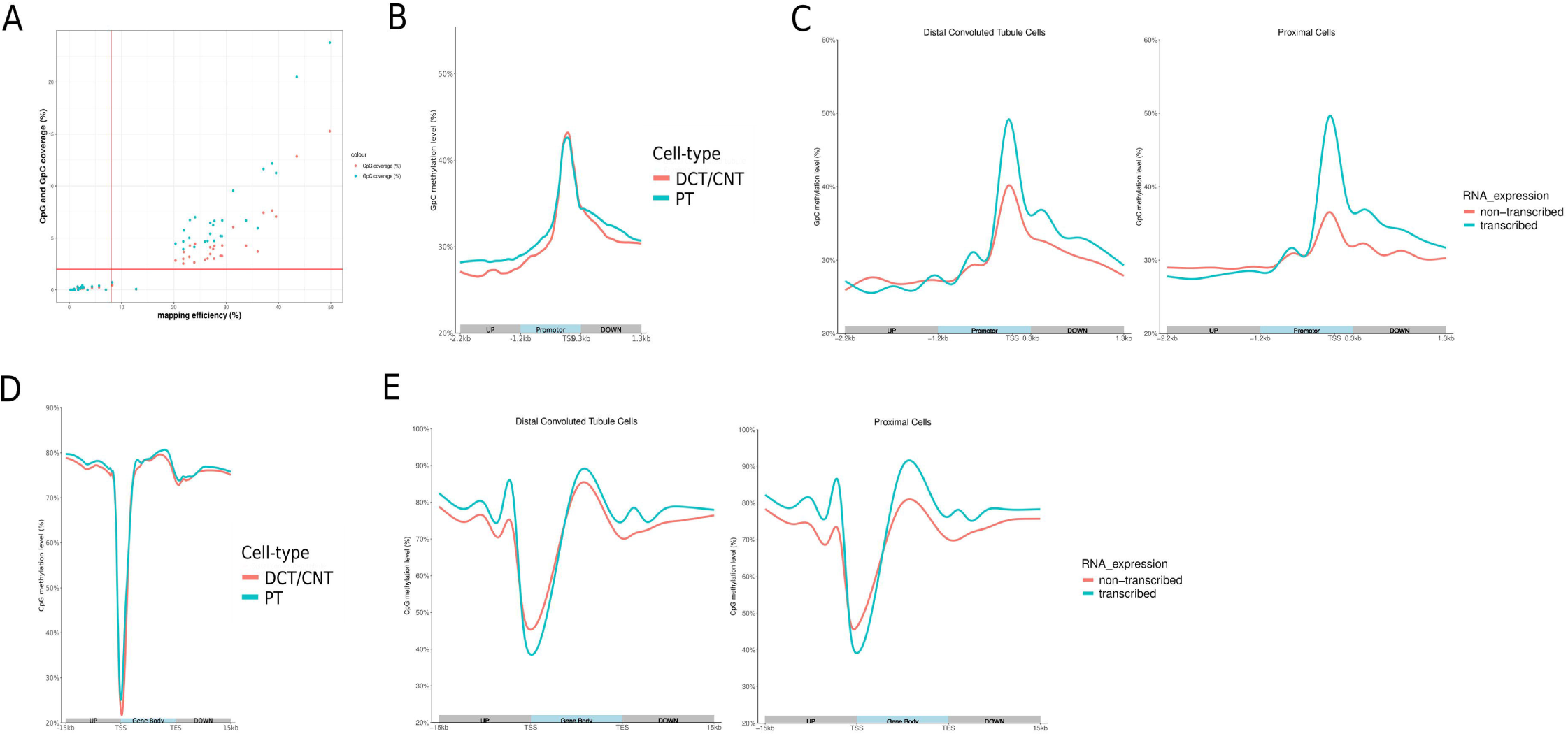
A) Percentage CpG and GpC sites covered by reads vs mapping efficiency (%). B) Promoter accessibility confirmed by GpC methylation of genes with open chromatin based on snATAC data common between PT and DCT. C) Promotor accessibility determined by GpC methylation of genes transcribed vs non-transcribed. D) CpG promoter and gene body methylation of genes with open chromatin based on ATAC data common between PT and DCT. E) CpG promoter and gene body methylation of genes transcribed vs non-transcribed.

**Supplementary Figure 6.**
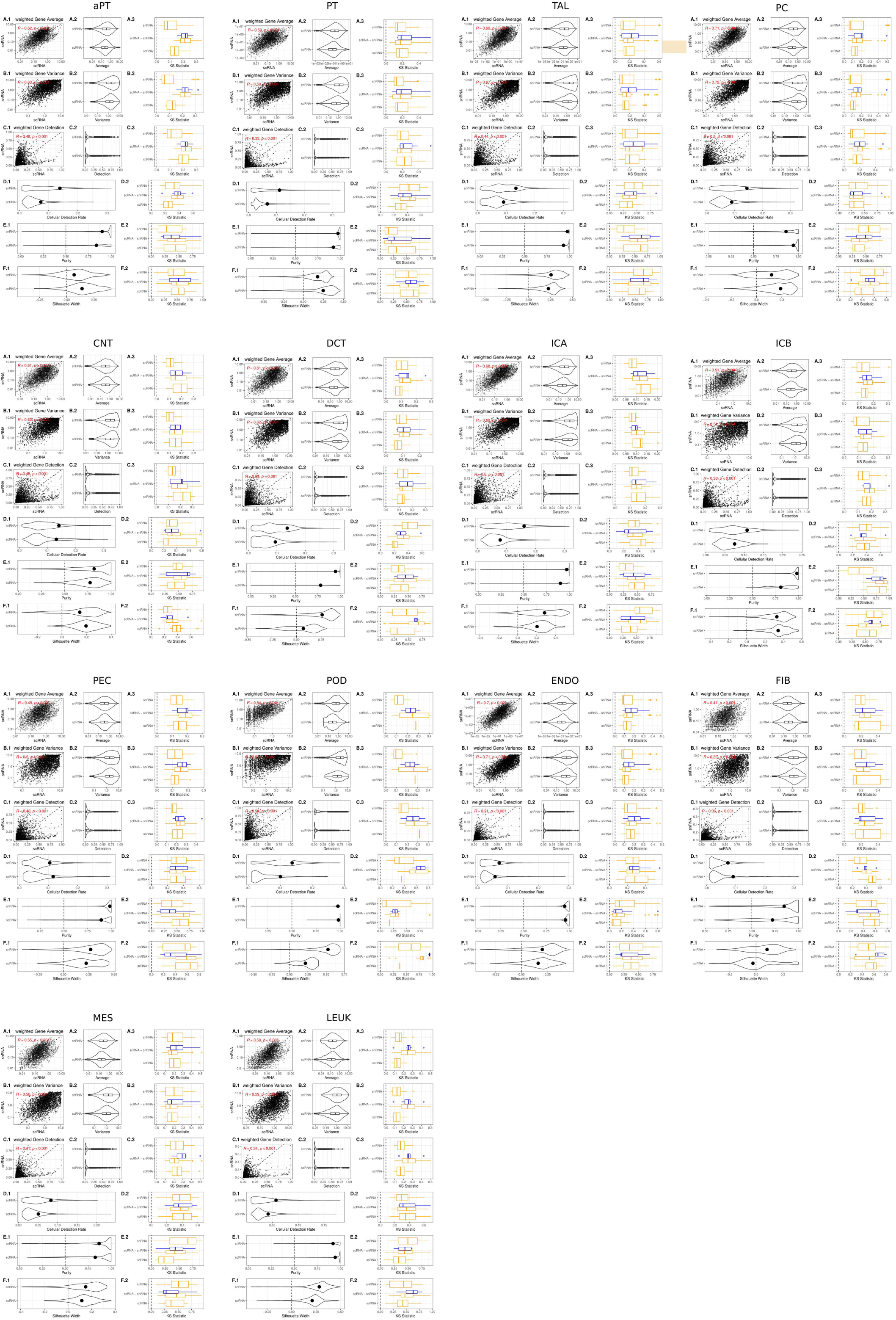
For each cell-type panel: A) Mean gene expression. A.1 Logarithmic weighted mean gene expression of snRNA-seq against scRNA-seq using log-normalized counts with dashed bisecting line and Pearson correlation coefficient in red. A.2 Logarithmic weighted mean gene expression values of snRNA-seq and scRNA-seq. A.3 KS Statistic comparing the distribution of mean gene expression values within snRNA-seq, between sc- and sn-RNA-seq and within scRNA-seq. Blue indicates within the same donor, orange across donors. B) Variance of gene expression. B.1 Logarithmic weighted gene expression variance of snRNA-seq against scRNA-seq using log-normalized counts with dashed bisecting line and Pearson correlation coefficient in red. B.2 Logarithmic weighted gene expression variance of snRNA-seq and scRNA-seq. B.3 KS Statistic comparing the distribution of gene expression variance within snRNA-seq, between sc- and sn-RNA-seq and within scRNA-seq. Blue indicates within the same donor, orange across donors. C) Detection of gene expression. C.1 Gene expression detection of snRNA-seq against scRNA-seq using log-normalized counts with dashed bisecting line and Pearson correlation coefficient in red. C.2 Gene expression detection of snRNA-seq and scRNA-seq. C.3 KS Statistic comparing the distribution of gene expression detection within snRNA-seq, between sc- and sn-RNA-seq and within scRNA-seq. Blue indicates within the same donor, orange across donors. D) Cellular Detection Rate. D.1 Proportion of nonzero gene counts in snRNA-seq and scRNA-seq. D.2 KS Statistic comparing the distribution of cellular detection rate within snRNA-seq, between sc- and snRNA-seq and within scRNA-seq. Blue indicates within the same donor, orange across donors. E) Purity. E.1 Purity of cell type cluster neighborhoods in snRNA-seq and scRNA-seq, dashed line indicates 50% pure neighborhood so half of the cells belong to the same cell type cluster. E.2 KS Statistic comparing the distribution of purity values within snRNA-seq, between sc- and snRNA-seq and within scRNA-seq. Blue indicates within the same donor, orange across donors. F) Silhouette. F.1 Silhouette values of cells within a cell type cluster in snRNA-seq and scRNA-seq, dashed line indicates 0 so below this value cells are closer to another cell type cluster or suited for forming their own cluster. F.2 KS Statistic comparing the distribution of silhouette values within snRNA-seq, between sc- and sn-RNA-seq and within scRNA-seq. Blue indicates within the same donor, orange across donors.

**Supplementary Figure 7.**
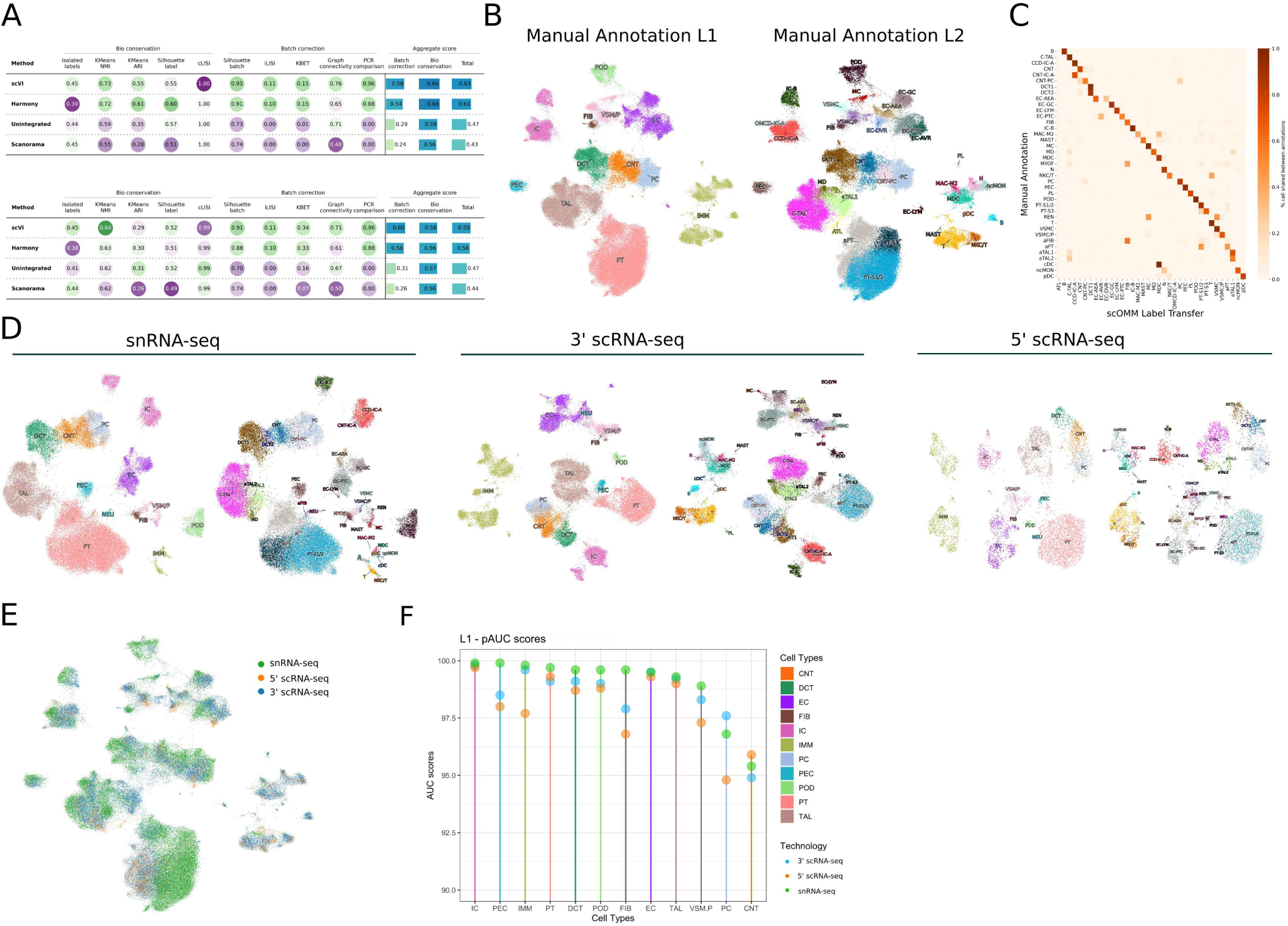
A) Benchmarking tables for Horizontal integration across mBDRC RNA protocols on the different cell type annotation granularity levels (L1 left, L2 right) showing methods ranked from best to poorest performance for bio conservation (preserving biological differences) and batch correction. B) scVI associated UMAP embedding (97,125 cells) of RNA protocols integration showing clustering-based annotation for the two cell type granularity levels L1 and L2. C) Heatmap showing the confusion matrix between Label transfer from HCA external resource (X axis) and Clustering-based annotation on L2 cell type annotation (Y axis). D) scVI associated UMAP embedding for individual RNA protocols (53,799 / 35,513 / 7813 cells respectively) showing transferred labels from HCA external resources for the two cell type granularity levels L1 and L2. E) scVI associated UMAP embedding (97,125 cells) of sn/scRNA protocol integration showing cells’ respective protocol. F) AUC score between label transfer annotation from HCA external resource and clustering-based annotation on L1 annotation level.

**Supplementary Figure 8.**
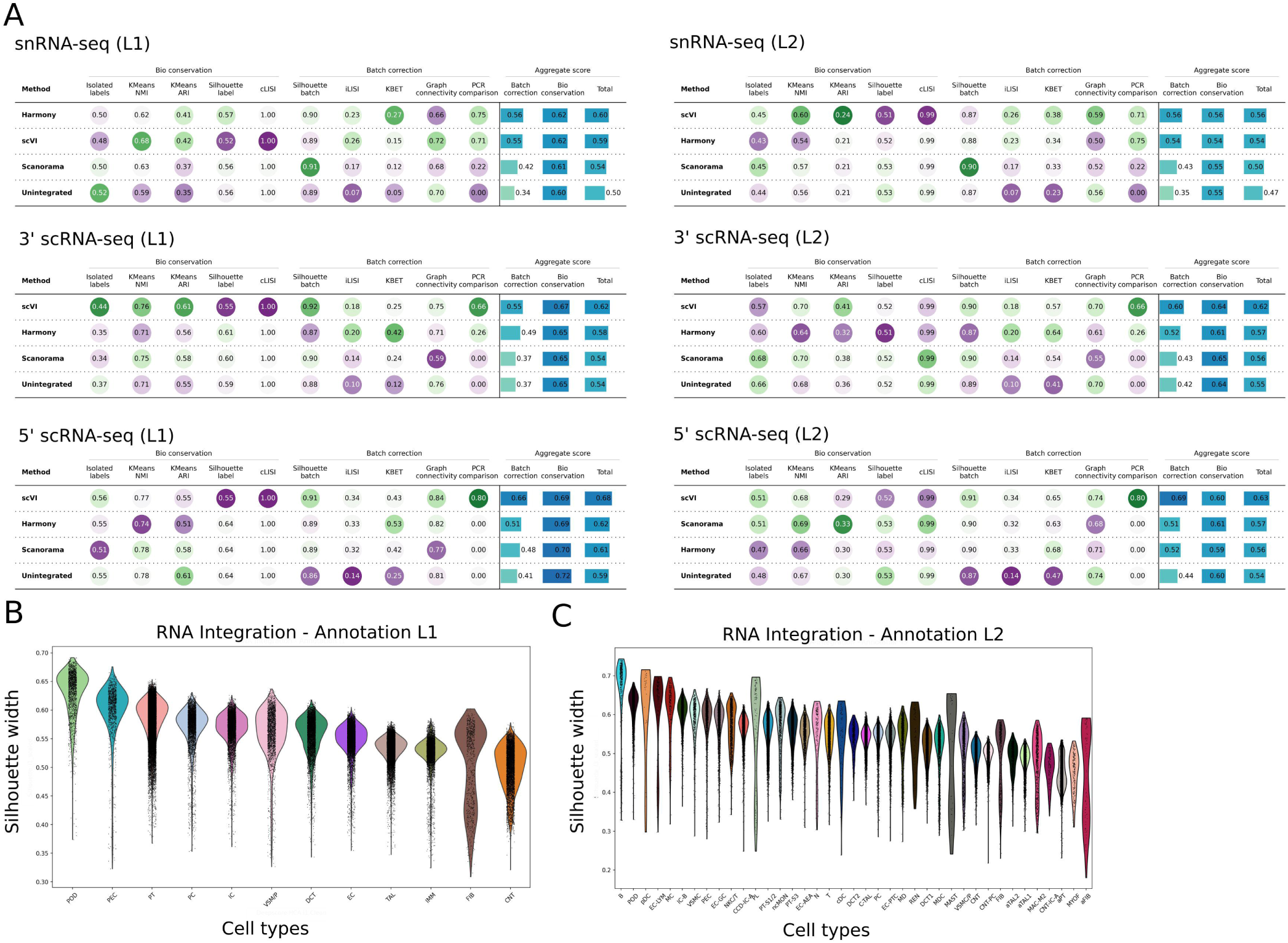
A) Benchmarking tables for Horizontal Integration of each mBDRC RNA protocols (Top to bottom: snRNA, 3’ scRNA, 5’ scRNA) on the different cell type annotation levels (L1 left, L2 right) showing methods ranked from best to poorest performance. B) Violin plots of silhouette width cell wise values by L1 transferred labels from HCA on scVI RNA Integration embedding. C) Violin plots of silhouette width cell wise values by L2 transferred labels from HCA on scVI RNA Integration embedding.

**Supplementary Figure 9.**
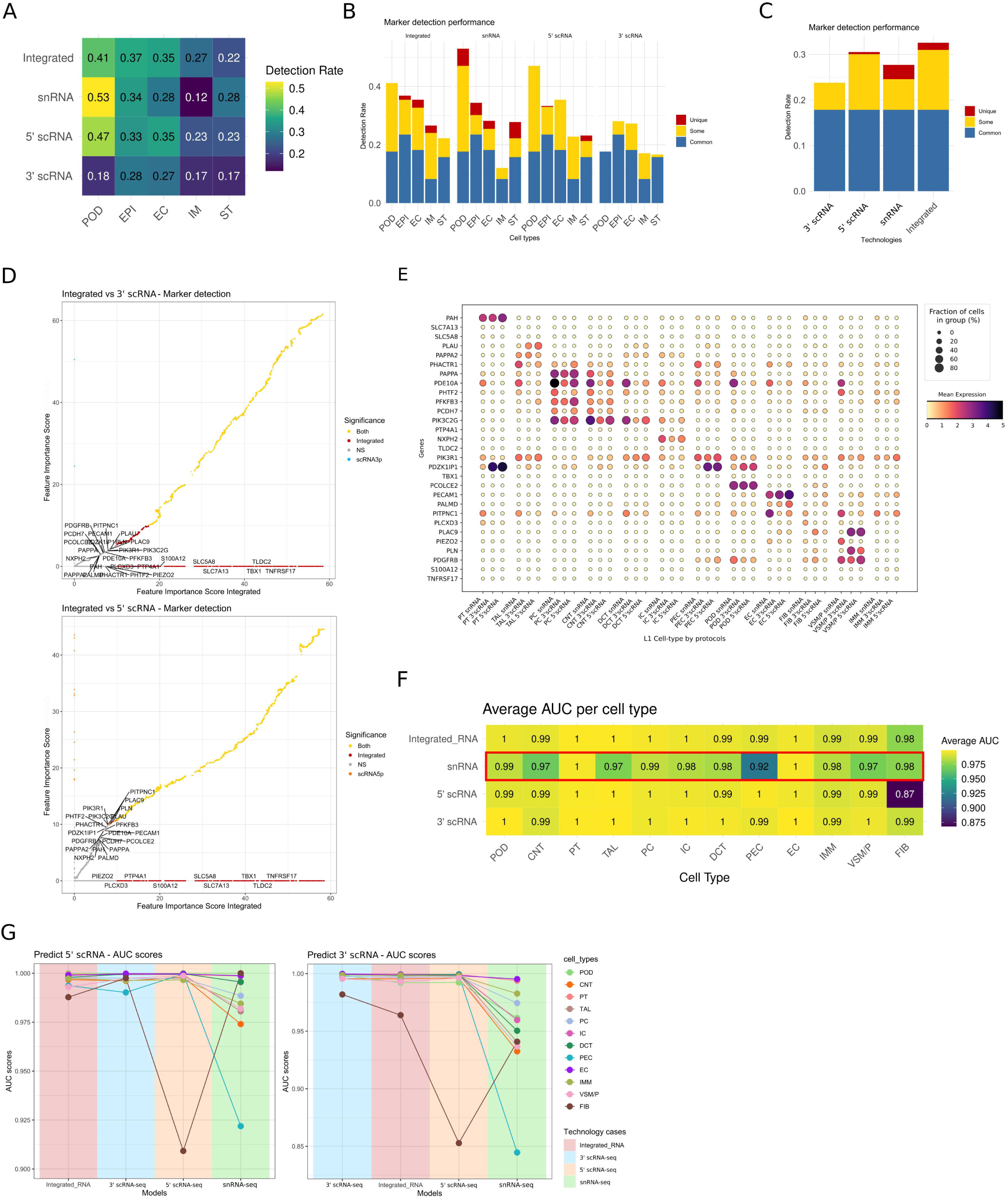
A) Marker detection rate comparison based on the feature importance of L1 groups across data models. B) Bar plots for Integrated model, snRNA, 5’ scRNA, and 3’ scRNA (left to right) showing fraction of markers composing detection rate metric in L1 groups that are common for every data model, shared by some, or uniquely detected by one data model. C) Bar plots for Integrated model, snRNA, 5’ scRNA, and 3’ scRNA (right to left) showing fraction of markers composing detection rate metric over every L1 group that are common for every data model, shared by some, or uniquely detected by one data model. D) Feature importance score comparison between 3’ scRNA-seq (left) / 5’ (right) and the integrated dataset for markers in Fig. 3G, which were found significant in snRNA-seq and integration models. E) Dot plot showing the log-normalized expression of highlighted genes in panel D across L1 cell-types per RNA protocol F) Heatmap showing AUC score between label transfer annotation from HCA external resource and clustering-based annotation on L1 annotation level across protocols. G) AUC scores for the predictability of each data model in predicting the 3’ scRNA-seq (top) / 5’ (bottom) data type for L1 cell-type annotations. Colored bars indicate distinct models for each technology, and dots represent L1 cell types.

**Supplementary Figure 10.**
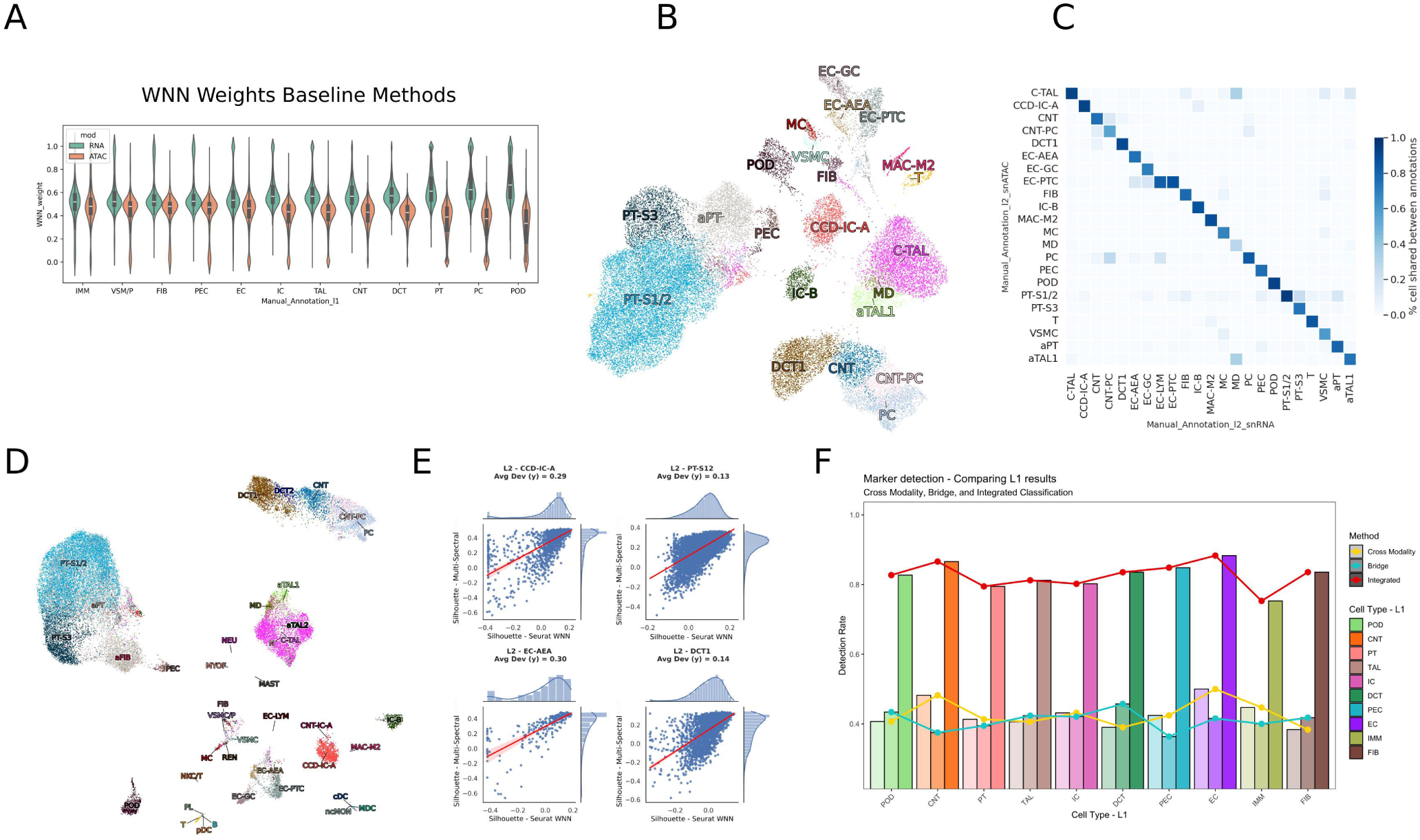
A) Violin plots of WNN modality weights per L1 cell type, obtained using the baseline embedding method implemented in Seurat. B) Spectral MNN- associated UMAP embedding (37,717 cells) showing snATAC-seq data clustering-based annotation at L2 resolution. C) Heatmap displaying the confusion matrix comparing clustering-based annotations from the best-performing horizontal integrations of both snRNA-seq (X-axis) and snATAC-seq (Y-axis) datasets using the same nuclei. D) WNN-associated UMAP embedding (37,717 cells) showing scOMM annotations obtained in the snRNA-seq data by projecting onto the HCA external reference data at L2 cell type resolution. E) Scatter plots comparing silhouette scores from optimized WNN (X-axis) and multi-spectral (Y-axis) integration methods across various kidney cell types and subtypes. Each blue dot represents a single cell, while the red line indicates the trend, with pink shading showing the confidence interval. Marginal histograms illustrate the distribution of silhouette scores along each axis. The “Avg Dev (y)” value represents the average deviation of Y-axis (multi-spectral scores) for each cell type or subtype. F) Marker detection rates from the scOMM model across different classification scenarios.

**Supplementary Figure 11.**
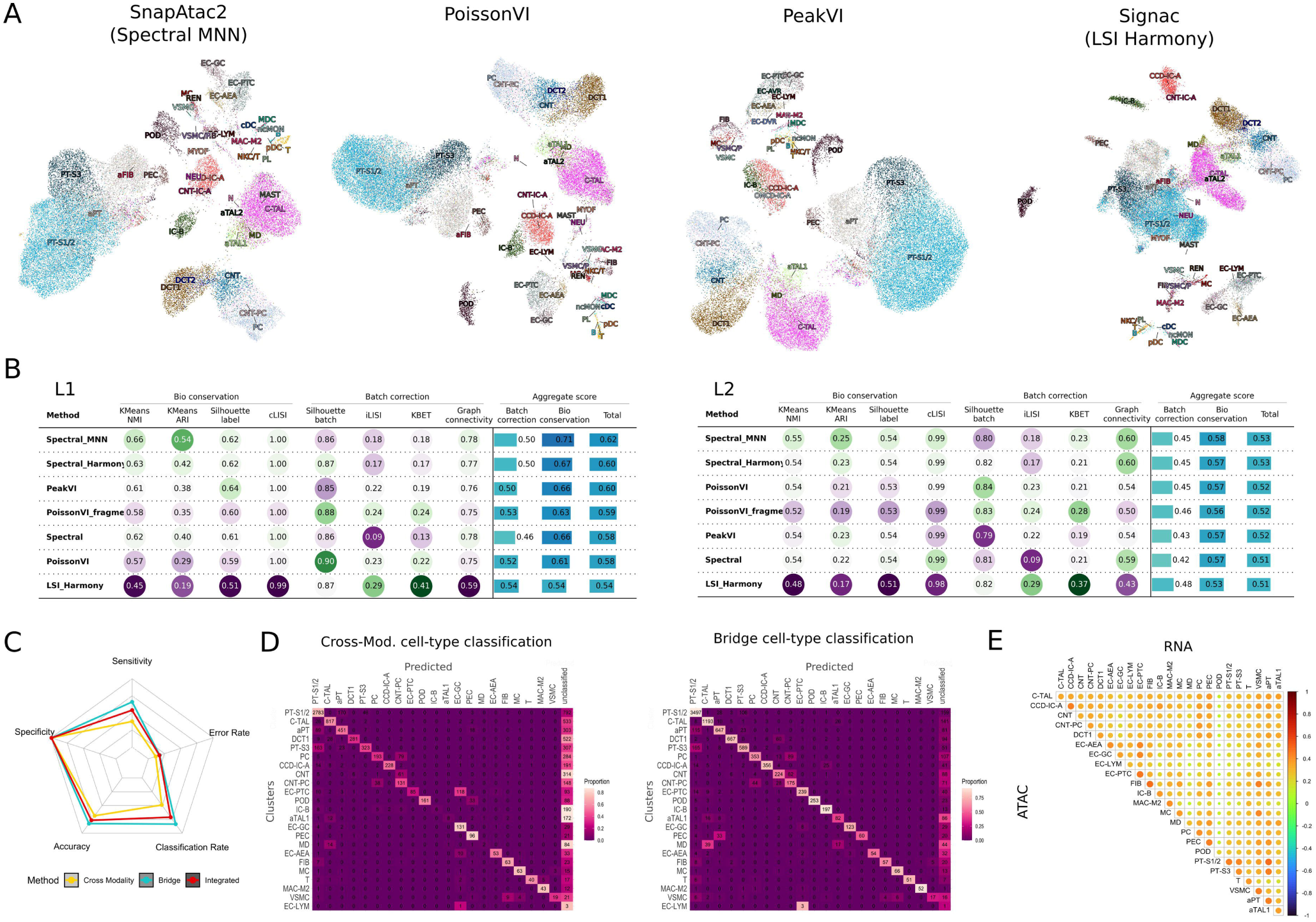
A) Benchmarking of methods associated with UMAPs embeddings (60,336 cells) of snATAC-seq showing label transferred annotations from HCA external resource for the L2 cell type granularity. B) Benchmarking tables for Vertical Integration of Multiome ATAC-RNA modalities on the different cell type annotation granularity levels (L1 & L2) showing methods ranked from best to poorest performance (top to bottom) C) Radar chart for model performance metrics (Sensitivity, Error Rate, Classification Rate, Accuracy and Sensitivity) across classification scenarios. D) Heatmaps showing the confusion matrix between clustering-based annotation on L2 cell type annotation and scOMM predictions for cross-modality classification (left) and bridge classification (right). E) Pearson correlation values across L2 groups between paired snRNA-seq gene expression profiles and gene activity profiles derived from snATAC-seq data from the same cells.

**Supplementary Figure 12.**
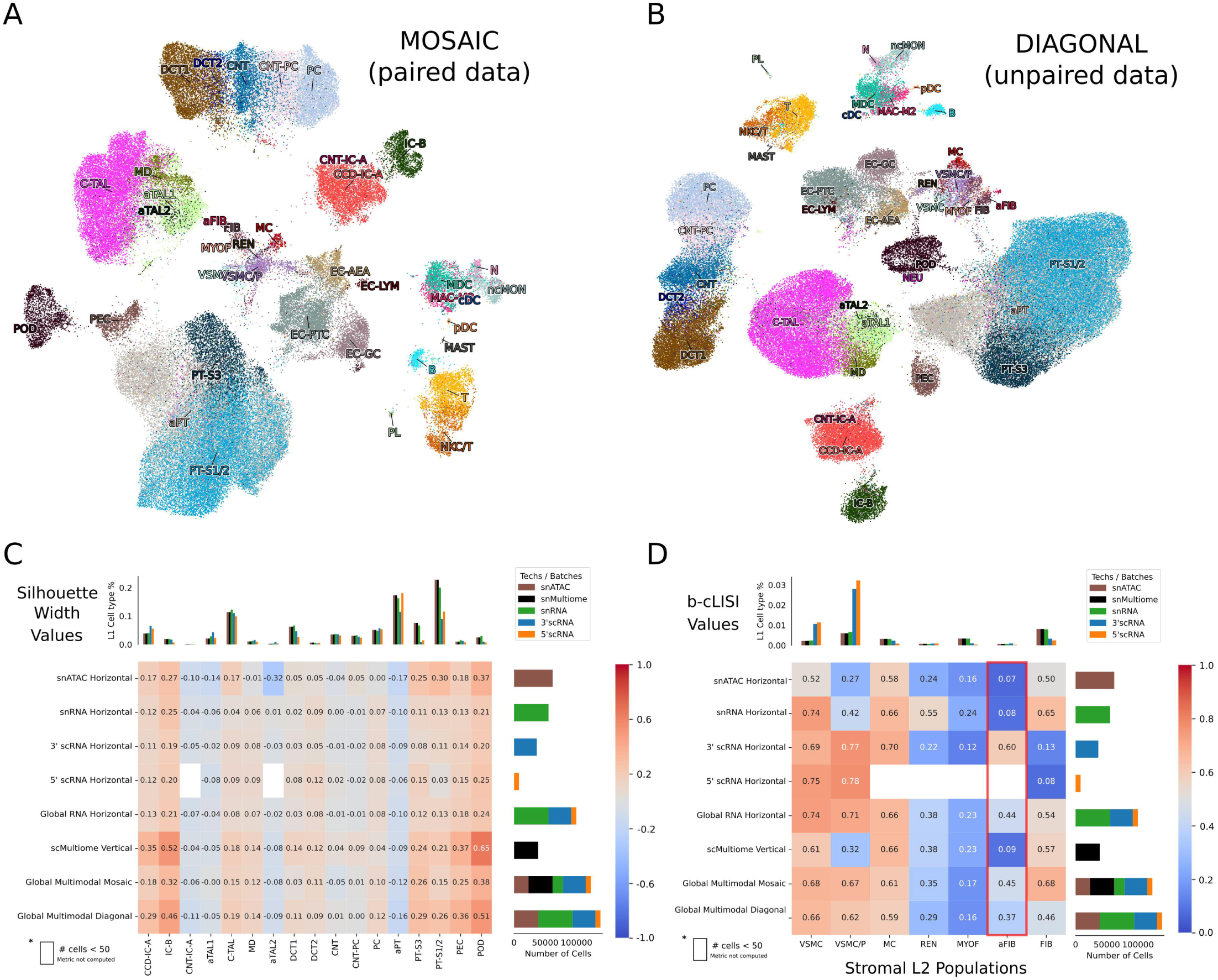
A) MultiVI associated UMAP embedding (119,744 cells) for mosaic integration showing scOMM annotations from HCA external resource with L2 cell type resolution. B) GLUE associated UMAP embedding (134,842 cells) for diagonal integration showing scOMM annotations from HCA external resource with L2 cell type resolution. C) Summary heatmap showing L2 cell type silhouette score over L1 cell type populations across every embedding scenario, unimodal and multimodal integrations. D) Summary heatmap showing L2 cell type silhouette score over L1 cell type populations for the stromal compartment across every embedding scenario, unimodal and multimodal integrations; aFIB population is highlighted with a red box.

**Supplementary Figure 13.**
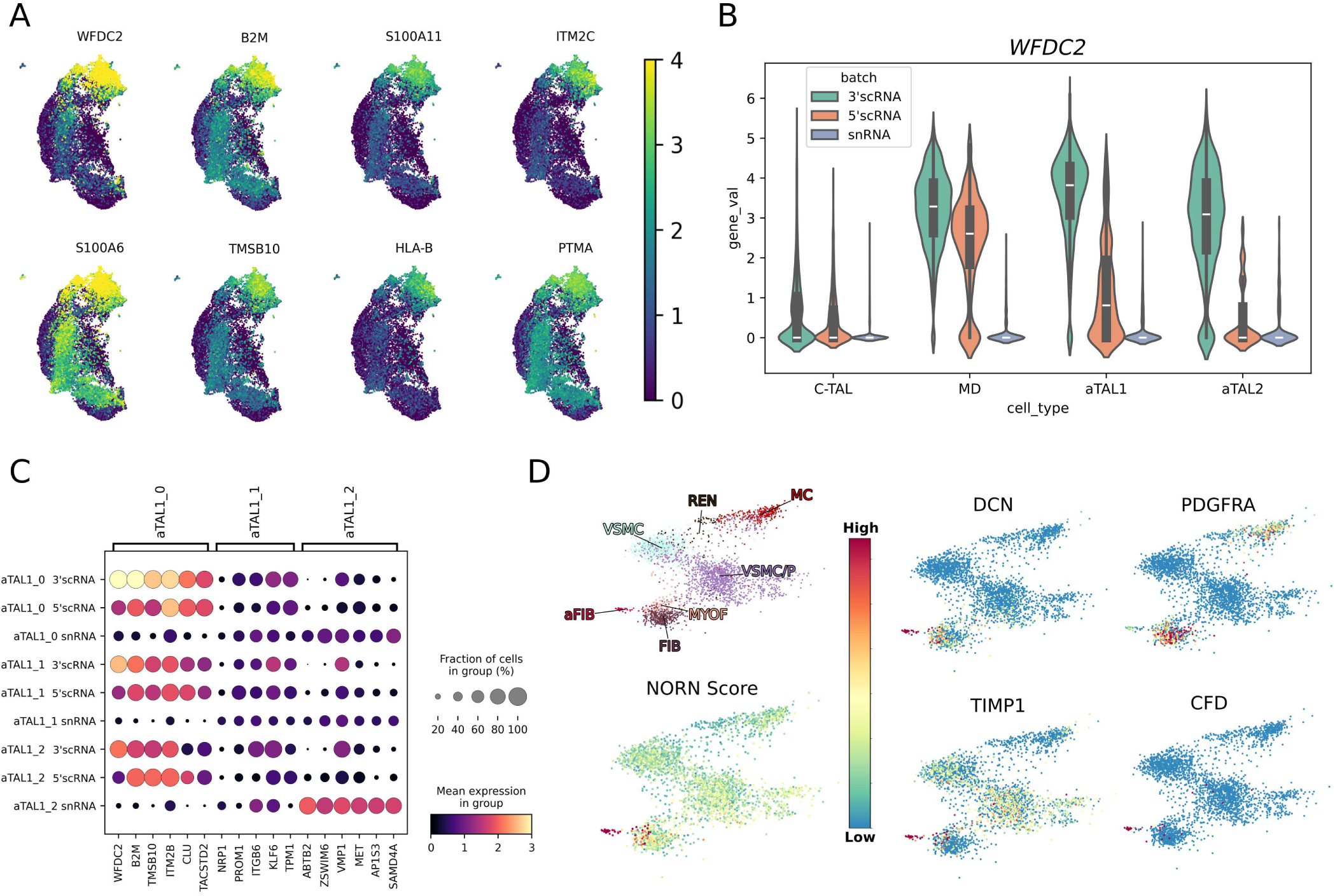
A) TAL compartment recomputed UMAP embedding (16,301 cells) as in Fig. 5B showing the log-normalized expression of top markers for aTAL1_0 cluster B) Violin plot of *WFDC2* log-normalized expression on TAL substates split by sn/scRNA-seq data. C) Average expression values of top marker genes across aTAL1 subclusters, split by RNA technology. D) Stromal compartment on UMAP embedding (3386 cells) of Horizontal Integration across mBDRC RNA protocols displaying scOMM L2 labels, together with Norn cells associated markers and Norn signature score.

**Supplementary Figure 14.**
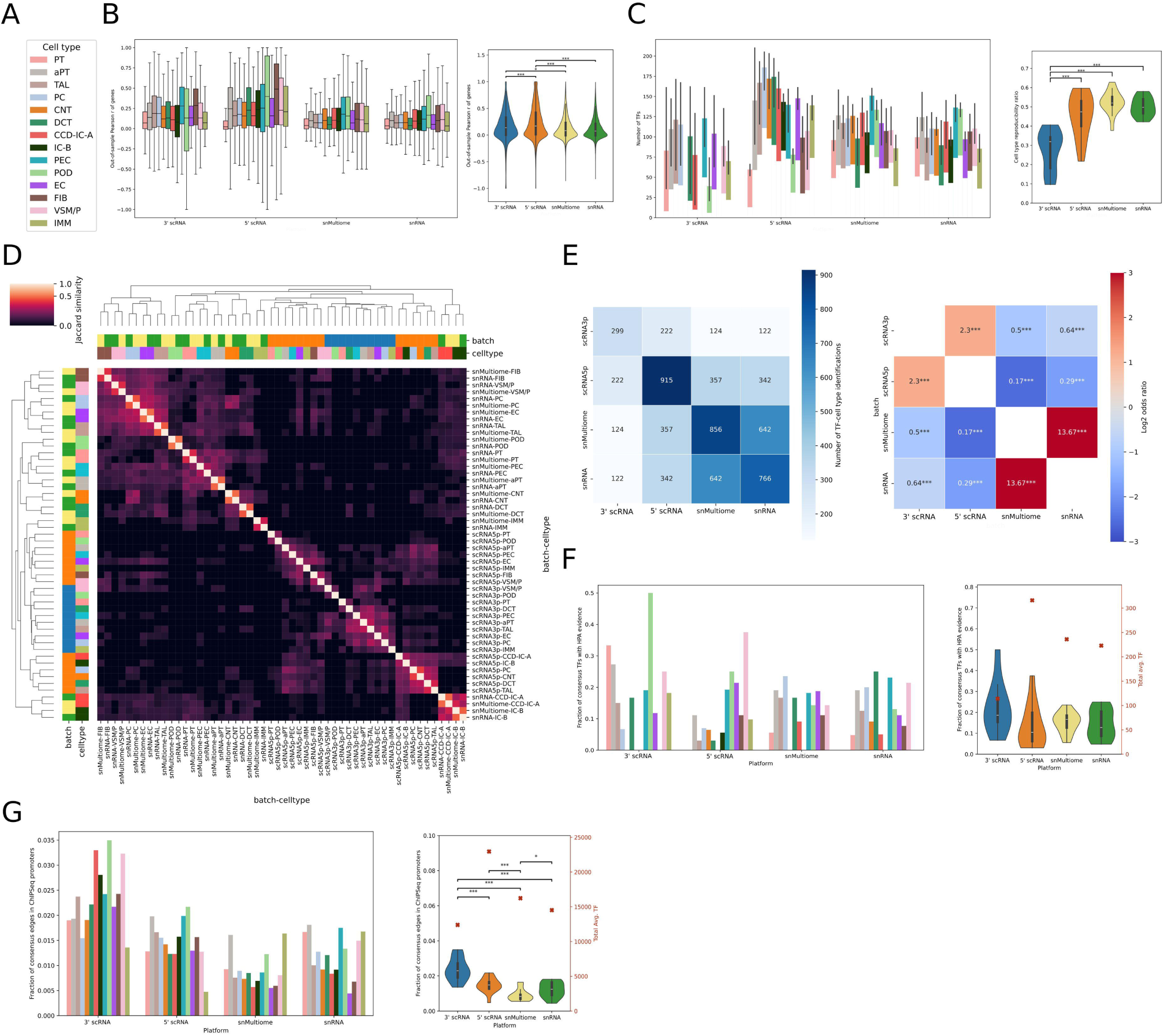
A) Cell-type annotation legend. B) Left panel: Pearson correlations between observed and out-of-sample predicted log-expression per gene, stratified by cell type. Right panel: Distribution of Pearson correlations, aggregated across cell types. Asterisks indicate significant differences in predictive accuracy across platforms (independent two-tailed T-test; *** = p < 0.001; * = p < 0.05). C) Left panel: Median number of transcription factors (TFs) identified per cell type across biological replicates, represented by the top of each bar, with whiskers indicating the minimum and maximum TF counts. The bottom of each bar represents the number of consensus TFs, found consistently in all replicates within each cell type. Right panel: Distribution of cell-type-specific “reproducibility ratios” per platform, calculated as the ratio of the number of TFs in the cross-sample consensus network to the mean number of TFs per replicate (independent two-tailed T-test; *** = p < 0.001). D) Ward’s hierarchical clustering of consensus regulatory networks per platform and cell type, based on Jaccard similarity of TF-target indicator vectors. Row and column annotations indicate platform (batch) and cell type. E) Left panel: number of cell type-TF assignments identified in each consensus network (diagonal) and shared between consensus networks across platforms (off-diagonal). Right panel: log-odds ratios of shared cell type-TF assignments across platforms (Fisher’s exact test; *** = p < 0.001). F) Left panel: Fraction of consensus TFs per platform that are also nuclear-localized in kidney tissue, per HPA immunohistochemistry annotations. Right panel: Distribution of these fractions across platforms, aggregated over cell types; no significant differences were observed between platforms (independent two-tailed T-test). G) Left panel: Fraction of consensus TF-target links per cell type with supporting evidence from intersected bulk ChIP-Seq and ATAC-Seq peaks at kidney promoters. Right panel: Distribution of these fractions across platforms, aggregated over cell types, with statistical significance between platforms indicated (independent two-tailed T-test; *** = p < 0.001).

